# Protein sequence evolution underlies interspecies incompatibility of a cell fate determinant

**DOI:** 10.1101/2025.08.02.668269

**Authors:** Emily L. Rivard, John R. Srouji, Anastasia Repouliou, Cassandra G. Extavour

## Abstract

Novel and rapidly evolving genes can integrate into conserved gene networks and play critical roles in development. Understanding how sequence variation in the orthologs of such genes influences functional interactions with the molecular products of older, more conserved genes requires investigation at the level of protein function. Here, we ask this question using the *oskar* gene as a case study. *oskar* is essential for primordial germ cell specification and embryonic patterning in fruit flies. Previous work showed functional incompatibility between *oskar* orthologs from the congeneric fruit fly species *Drosophila melanogaster* and *Drosophila virilis*. However, the region(s) of Oskar in which protein-coding sequence evolution led to this incompatibility, and the molecular mechanisms underlying it, remained unknown. To address these questions, we generated chimeric versions of *oskar* comprising different combinations of Oskar protein domains from each species, expressed these chimeric *oskar* sequences in *D. melanogaster*, and quantified their ability to assemble functional germ line and abdominal patterning determinants (germ plasm). We found that a specific portion of *D. virilis* Oskar, namely the OSK domain, was primarily responsible for the cross-species incompatibility of Oskar. In the absence of endogenous *D. melanogaster* Oskar, chimeras containing the *D. virilis* OSK domain could not localize posterior germ plasm well enough to generate primordial germ cells, but were sufficient to specify the anteroposterior axis. We also found that the *D. virilis* OSK domain had dominant-negative effects on *D. melanogaster* Oskar’s ability to localize germ plasm mRNAs, resulting in severe axial patterning defects. Both *in vivo* and *in silico* analyses support the hypothesis that evolved changes in the ability of the OSK domain to interact with conserved germ plasm mRNAs played a significant role in the evolution of interspecies incompatibility of Oskar. Under this hypothesis, an essential germ line determinant evolved to be incompatible across species of the same genus in less than 50 million years, while retaining functional within-species molecular interactions. Thus, by investigating *in vivo* protein function, we link genomic and molecular evolution with phenotypic variation and fitness at higher scales.

## Introduction

Genes with essential roles in development often exhibit strong sequence conservation ^1–3^. However, others undergo rapid sequence evolution despite being critical for development ^4–7^. While *cis*-regulatory sequence changes frequently drive phenotypic evolution ^8–11^, evolution in protein-coding sequences can also have dramatic functional consequences for development ^12–14^. Unfortunately, simply identifying amino acid substitutions that arise during evolution cannot tell us whether or how they generate phenotypic change ^15–17^. Studying the evolution of protein function *in vivo* at a molecular scale—through altered interactions with other molecules, localization patterns, or enzymatic activity—can provide data needed to link sequence changes with biological consequences at cellular and organismal levels.

Germ cell specification is critical not only for the development of an organism, but also for the continuity of its lineage. A suite of ancient, conserved genes function in germ line segregation across metazoans ^18,19^. However, multiple convergently evolved lineage-specific and quickly evolving genes have become integrated into this core germ line gene network and acquired essential roles in germ line specification. The best studied example of such a gene is the gene *oskar*, which arose in insects at least 325 million years ago ^20,21^ and subsequently gained a role in germ cell specification in some species, including *Drosophila melanogaster*. Oskar protein is the only factor necessary and sufficient for assembly of the germ plasm, the specialized cytoplasm at the egg posterior comprised of maternally provided germ line and abdominal patterning molecules ^22^. Specifically, Oskar nucleates formation of granules containing these molecules, and the cytoplasm containing these granules acts as a germ line determinant. Thus, cells inheriting these granules, called pole cells, are specified as the primordial germ cells ^23^.

*D. melanogaster* Oskar protein structure and function have been interrogated with genetic and biochemical assays over the last several decades. Oskar has two ordered domains, an N-terminal LOTUS domain and a C-terminal hydrolase-like OSK domain (Fig. 1B) ^24–26^. The LOTUS domain binds to Vasa protein *in vitro*, and Vasa recruitment to the germ plasm by Oskar is required for germ cell formation ^27–29^. The LOTUS domain of Oskar dimerizes *in vitro*, a capacity that could be relevant for its function in *D. melanogaster* ^24–26^. The OSK domain binds germ plasm mRNAs *in vitro* ^24,25^ and is required for proper recruitment of multiple germ plasm mRNAs *in vivo* ^30^. An intrinsically disordered region linking LOTUS and OSK may mediate interactions with several additional germ line molecules ^31–33^. Finally, while the aforementioned regions comprise the shorter translational isoform of Oskar that nucleates germ granules, there is a longer Oskar isoform with an additional N-terminal disordered domain called the Long Osk domain (Fig. S1B) ^34^. Rather than interact with germ granule components, the Long Oskar isoform is required for correct anchoring of *oskar* RNA/protein and mitochondria at the posterior ^35,36^. The Long Osk domain also modulates F-actin dynamics and endocytic activity at the posterior cortex of the oocyte, both of which influence how well germ plasm is anchored ^37–40^.

**Figure 1.**
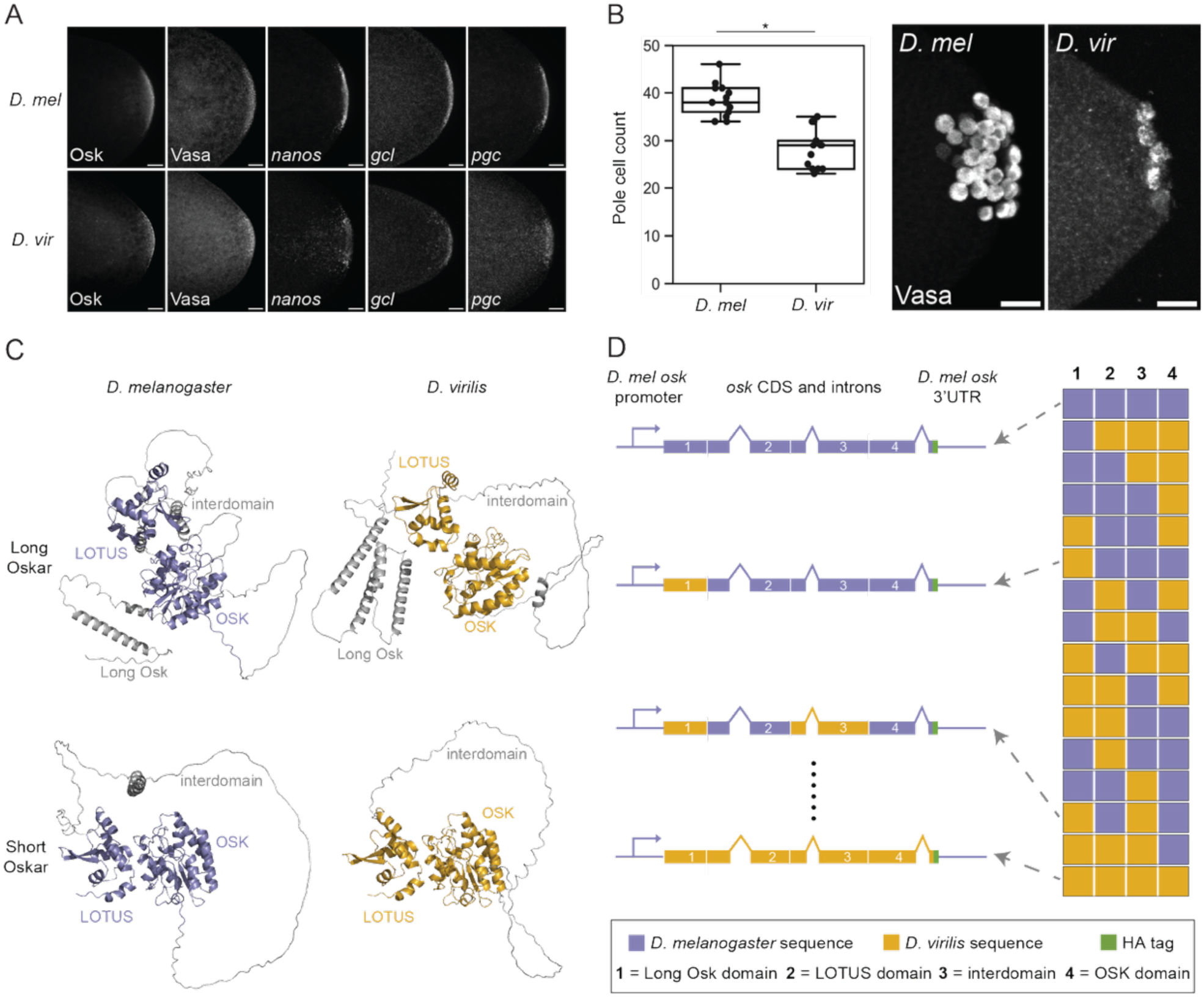
*D. virilis* exhibits posterior localization of several germ plasm molecules, including Oskar, which shares a similar domain architecture with *D. melanogaster* Oskar. A) Posterior poles of *D. melanogaster* (*D. mel*, top row) and *D. virilis* (*D. vir*, bottom row) stage 2 embryos with Oskar, Vasa, *nanos*, *germ cell-less* (*gcl*), and *polar granule component* (*pgc*) labeled. **B)** Pole cell counts for *D. virilis* and *D. melanogaster* stage 5 embryos. Asterisk designates significance (p-value < 0.01, estimated from the distribution of difference of means between sample and control from 100,000 simulated samples). n = 13 embryos for each species. B’) *D. melanogaster* and *D. virilis* stage 5 embryos with germ cells labeled with anti-*D. melanogaster* Vasa antibody. All micrographs are maximum intensity projections of 20 optical sections (step size = 0.5 *μ*m) oriented with posterior to the right and scale bars = 20 *μ*m. **C)** AlphaFold3 ^49^ structural predictions of Long and Short Oskar isoform orthologs from *D. melanogaster* and *D. virilis* with the Long Osk, LOTUS, interdomain, and OSK domains labeled. **D)** Schematic of *D. melanogaster–D. virilis oskar* chimeric constructs. *D. melanogaster* (purple) and *D. virilis* (gold) genomic DNA sequences encoding the four domains shown in (**C**) were fused in frame in all possible combinations. The *D. melanogaster oskar* promoter, 5’UTR, and 3’UTR were used for all constructs. An HA tag (green) was added to the C-terminus of all sequences. The rightmost portion of the schematic represents the 16 different chimeras generated for this experiment.

Despite its essential role in germ plasm assembly, which largely relies on direct interactions with other germ plasm molecules, *oskar* exhibits rapid protein sequence divergence compared with other *Drosophila* germ line genes ^6,7^. Indeed, even within the genus *Drosophila*, *oskar* has diverged so much that some orthologs are no longer functionally interchangeable ^41,42^. The ortholog of *oskar* from *D. virilis*, a species 40-50 million years diverged from *D. melanogaster* ^43^, was previously reported to rescue abdominal patterning defects, but not the loss of germ cells, when expressed in *D. melanogaster oskar* protein null flies (*osk[2]* (also called *osk[84]*) or *osk[6]* (also called *osk[166]*)) ^42^. Further, the same study reported that *D. virilis oskar* induced dominant-negative patterning defects in a wild-type background ^42^. However, this study raised at least three open questions, which have remained unanswered to date. First, did the trans-species expression vector used yield correct translation or localization of *D. virilis* Oskar protein in *D. melanogaster*? Second, what was the mechanism underlying the dominant-negative impacts of *D. virilis oskar* expression? Third, which region(s) of *D. virilis* Oskar were responsible for its failure to drive pole cell formation in *D. melanogaster*?

While there is substantial evidence that all four major domains of the Oskar protein play important germ plasm assembly roles, the sequence changes that prevent *D. virilis oskar* from specifying germ cells in *D. melanogaster* are unknown. Although evidence suggests that the OSK domain interacts with germ plasm mRNAs ^24,25,30^, the LOTUS domain interacts with Vasa protein ^27–29^, the interdomain interacts with Actin-binding proteins ^32^, and the Long Oskar domain contributes to mitochondrial localization ^35^, none of these reported domain abilities in *D. melanogaster* indicate that evolved changes in their sequences should or should not be expected to prevent *D. virilis* Oskar from carrying out these functions in a *D. melanogaster* context. Thus, given the essential roles of all four domains of the Oskar protein, it is not clear *a priori* whether any one of these domains might be more of a contributor than others, to inter-species incompatibility.

In principle, inter-species incompatibility of Oskar could be due to evolutionary changes in the sequences of interacting germ plasm mRNAs, rather than in the Oskar protein itself. Indeed, mRNA sequence divergence between lineages can cause changes in germ granule composition for some mRNAs ^44^. However, several previous observations lead us to hypothesize that evolutionary divergence in the Oskar protein is more likely to explain why *D. virilis* Oskar cannot assemble functional germ plasm in *D. melanogaster*. First, *D. virilis nanos* mRNA can be effectively localized to the posterior in *D. melanogaster* embryos (presumably by *D. melanogaster* Oskar), and the 3’UTR of *nanos* mRNA shows high conservation between these two species ^45^. Furthermore, unlike *oskar*, *D. virilis nanos* can functionally substitute for *D. melanogaster nanos* ^46^. In addition, previous studies have failed to find conserved 3’UTR signatures of germ plasm localization shared across all known mRNA germ plasm components ^47,48^. Taken together, these data suggest that evolutionary divergence of germ plasm mRNA UTR sequences is unlikely to provide a simple answer to explain why *D. virilis* Oskar protein cannot assemble functional germ plasm in *D. melanogaster*.

We posit that investigating the basis of this incompatibility could shed light on the larger problem of the evolution of complex molecular regulatory networks whose members have very different molecular evolutionary dynamics, and potentially also on the functional significance of specific portions of *oskar*. Here, we elucidate how protein-coding sequence evolution in *oskar* between these two species influences its ability to assemble functional germ plasm and mediate germ cell specification and abdominal patterning. By expressing *D. melanogaster–D. virilis* chimeric *oskar* sequences in *D. melanogaster* and characterizing their ability to perform Oskar’s functions, we identified Oskar protein domains from *D. virilis* that cause failure to form germ cells and impede recruitment of some germ plasm mRNAs to the posterior. We also identified portions of the *D. virilis* Oskar protein with dominant-negative effects on *D. melanogaster* Oskar’s ability to recruit germ plasm mRNA and mediate germ cell formation and axial patterning. Finally, we characterized Oskar-driven changes in how germ granules form and localize at the posterior in *D. virilis* and *D. melanogaster*. This evolution-informed mutagenesis strategy sheds new light on structure-function relationships *in vivo* for an important developmental gene. Furthermore, our findings suggest that even within a conserved gene network, molecular interactions among gene products can evolve in species-specific ways. This highlights the molecular flexibility of developmental programs and underscores the importance of continued investigation at the molecular mechanistic studies to understand interspecies variation in development.

## Results

### Germ plasm molecules recruited by Oskar in D. melanogaster are also localized to the posterior in D. virilis

We first asked how germ plasm assembly, primordial germ cell formation, and Oskar’s predicted structure compare between wild-type *D. virilis* and *D. melanogaster*. We used hybridization chain reaction (HCR) *in situ* hybridization and immunofluorescence to determine whether several key germ plasm molecules in *D. melanogaster* localize to the germ plasm of oocytes and embryos of *D. virilis*. We found that, as in *D. melanogaster*, *D. virilis oskar* mRNA and protein localize to the posterior end of oocytes and cleavage stage embryos (Fig. 1A, Fig. S1A). We also observed Vasa protein, mitochondria (detected with an anti-ATP5A antibody), and *nanos, polar granule component* (*pgc*), and *germ cell-less* (*gcl*) mRNAs enriched at the posterior of *D. virilis* eggs (Fig. 1A, Fig. S1A). These data demonstrate that several of the molecules that Oskar recruits to germ plasm in *D. melanogaster* are also found in *D. virilis* germ plasm.

We also used anti-Vasa antibody staining to label and count pole cells in stage 5 embryos from both species. We found that *D. virilis* has significantly fewer pole cells at this stage than *D. melanogaster* (Fig. 1B). This may result from the reduced expression of some germ line determinants in this species ^44^ or from differences in the composition, organization, or function of germ plasm.

Having established that *D. virilis* germ plasm contains known Oskar interaction partners, and that germ cells appear to be spatially and temporally specified in the same way, albeit in different quantities, we next investigated differences in Oskar primary sequence and predicted secondary and tertiary structure between these species. We found that the AlphaFold3 ^49^ structural predictions for *D. melanogaster* and *D. virilis* Oskar share a domain architecture consisting of four distinct regions (Fig. 1C, Fig. S1A–B). The longer of Oskar’s two translational isoforms starts with an N-terminal disordered region called Long Osk, which is 22 amino acids longer in *D. virilis* than in *D. melanogaster* and contains two additional predicted alpha helices (Fig. 1C, Fig. S1A–B). The first folded domain, LOTUS, exhibits a high degree of structural similarity between these species (RMSD = 0.263) and shares 69% amino acid identity (Fig. S1B–C). The disordered interdomain that links Oskar’s two folded domains is 33 amino acids shorter in *D. virilis* than in *D. melanogaster* and lacks much secondary structure (Fig. S1B–C). The C-terminal folded domain, OSK, is also structurally similar between the two species (RMSD = 0.297) and shares 76% identity (Fig. S1B–C). Thus, the modular organization of Oskar is similar between *D. melanogaster* and *D. virilis,* with higher sequence and structural similarity within the folded regions and greater variability in the disordered portions of the protein.

This shared organization suggested that designing chimeric *D. melanogaster–D. virilis* Oskar proteins with different combinations of the four domains could be an effective strategy for determining the region(s) of Oskar where evolutionary divergence was responsible for the interspecies incompatibility (Fig. 1D). We therefore generated sixteen chimeric versions of Oskar by swapping its four domains between the two species in all possible combinations. To test the function of these chimeric Oskar proteins in *D. melanogaster,* we expressed these constructs in *D. melanogaster* under the control of the *D. melanogaster* genomic regulatory region. We evaluated the abilities of these chimeric Oskar proteins in two genetic backgrounds:

1) an *oskar* RNA null background to evaluate the chimera’s ability to rescue the *oskar* null, and
2) a background with endogenous *oskar* present to assess potential interactions with *D. melanogaster oskar*.

We performed several experiments to ask whether the behaviors of our chimeric Oskar proteins were sufficiently similar to those of endogenous *D. melanogaster* Oskar, to allow us to make inferences about the functional impacts of interspecies protein sequence divergence. First, we quantified enrichment of each chimera at the posterior, and found that despite some variation in detectable expression levels (Figs. S2, S3), this did not correlate with any of the germ plasm component, pole cell formation, or axial patterning phenotypes that we assessed. Further, we determined that chimeric *oskar* mRNAs and proteins localized correctly to the oocyte posterior (Fig. S2, S4, Table S1). In *D. melanogaster*, *oskar* mRNA splicing contributes to its correct localization ^50–52^. Although we did not explicitly analyze chimeric *oskar* mRNA processing, we confirmed that correct localization, as well as localization and translation of chimeric Oskar proteins, occurred in all chimeras (Figs. S2–S4). Moreover, all chimeras permitted oogenesis to proceed normally, which does not happen when *D. melanogaster oskar* splicing is disrupted ^50,53,54^. Thus, we were able to attribute the germ plasm, pole cell, and body patterning phenotypes we assessed, to species-specific protein domain abilities in the experiments described below.

### The D. virilis OSK domain impedes germ plasm assembly in D. melanogaster

We first tested whether the chimeric Oskar proteins could induce pole cell formation at the posterior of stage 5 embryos. Consistent with previous studies using a *D. melanogaster oskar* protein null (*osk[2]* (also called *osk[84]*) or *osk[6]* (also called *osk[166]*)) background ^42^, we found that *D. virilis* Oskar completely failed to specify pole cells in *D. melanogaster oskar* RNA null embryos *(y[1] sc[*] v[1]; Kr[If-1]/CyO; TI{w[+mC]=TI}osk[0.EGFP]*^55^; Fig. 2A–B). Oskar (marked by the HA tag on our transgenes) and Vasa proteins were detected in posterior somatic blastoderm cells (Fig. 2A). Only nine chimeric Oskar proteins could induce any pole cells to form, and only three of those could induce as many pole cells as *D. melanogaster* Oskar (Fig. 2B, Table S2). Notably, inclusion of the *D. virilis* OSK domain caused near (two of eight) or complete (six of eight) failure to generate pole cells (Fig. 2B), suggesting that this domain is an important contributor to *D. virilis oskar*’s functional incompatibility in *D. melanogaster.* We also observed some reduction in pole cell number by chimeras containing the LOTUS domain from *D. virilis* in combination with the interdomain from *D. melanogaster* (Fig. 2B). These reductions did not occur in chimeras that had the *D. virilis* LOTUS and *D. virilis* interdomains together (Fig. 2B), suggesting that the functional properties of Oskar’s individual domains may be influenced by neighboring protein sequence.

**Figure 2.**
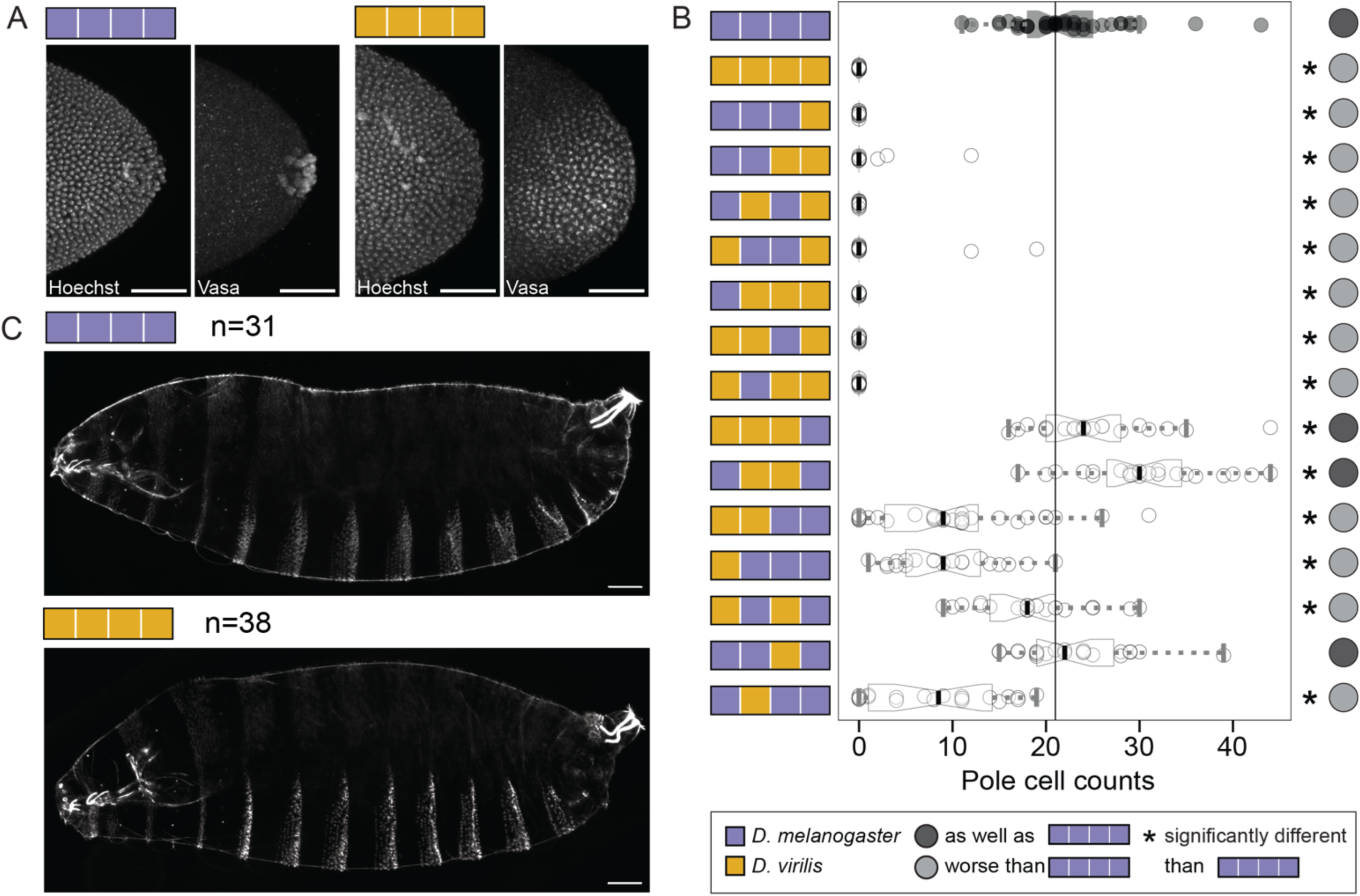
*D. virilis* Oskar and chimeras with the *D. virilis* OSK domain fail to rescue pole cell formation but rescue embryonic patterning defects. A) Posterior poles of stage 5 embryos from *D. melanogaster* mothers expressing *D. melanogaster* (purple, left) or *D. virilis oskar* (gold, right) in an *oskar* null background. Embryos are stained with a nuclear marker (Hoechst) and anti-*D. melanogaster* Vasa antibody. Micrographs are maximum intensity projections of 60 optical sections (step size = 1 *μ*m) and scale bars = 50 *μ*m. **B)** Pole cell counts in stage 5 embryos expressing chimeric *oskar*. Distribution for embryos from flies expressing the *D. melanogaster oskar* chimera (positive control) is in black with the median marked with a black line. Significant difference from positive control distribution (p-value < 0.05, estimated from the distribution of difference of means between sample and control from 100,000 simulated samples) indicated by asterisk. Circles to the right of the plots indicate data interpretation as follows: dark gray = specifies pole cells as well as *D. melanogaster oskar*; light gray = specifies pole cells worse than *D. melanogaster oskar*. Sample sizes ≥15 for each genotype; see Table S2. **C)** Cuticles of first instar larval progeny of mothers expressing *D. melanogaster* (purple, left) and *D. virilis oskar* (gold, right) exhibit wild-type patterning. Micrographs are projections of 15 optical sections (step size = 1.47 *μ*m). Scale bars = 50 *μ*m. Posterior is to the right in all micrographs.

As previously reported ^42^, we found that although *D. virilis oskar* cannot mediate pole cell formation, it can perform *oskar*’s axial patterning role. Specifically, first instar larval progeny of mothers expressing *D. virilis oskar* exhibited wild-type abdominal patterning (Fig. 2C). This result suggests that *D. virilis oskar* can successfully recruit abdominal patterning determinants to the posterior for abdominal specification.

### D. virilis Oskar and chimeric Oskar proteins effectively recruit Vasa but not mRNAs to germ plasm

To better understand how the chimeras either succeeded or failed to perform Oskar’s roles in cell fate specification, we examined the localization of a selection of germ plasm molecules that are normally recruited by *D. melanogaster* Oskar to the posterior. Specifically, we measured the integrated enrichment of signal for the 15% posterior-most portion of the egg in immunofluorescence and HCR experiments designed to detect particular germ plasm components (Figs. 3, 5, S5, S6). We also assessed colocalization of the chimeric Oskar proteins with each germ plasm molecule by calculating the Pearson correlation for the signal intensity between HA and another germ plasm molecule, for the pixels in the 15% posterior-most end of the embryo that had HA intensity values two standard deviations over the mean (Fig. S7).

**Figure 3.**
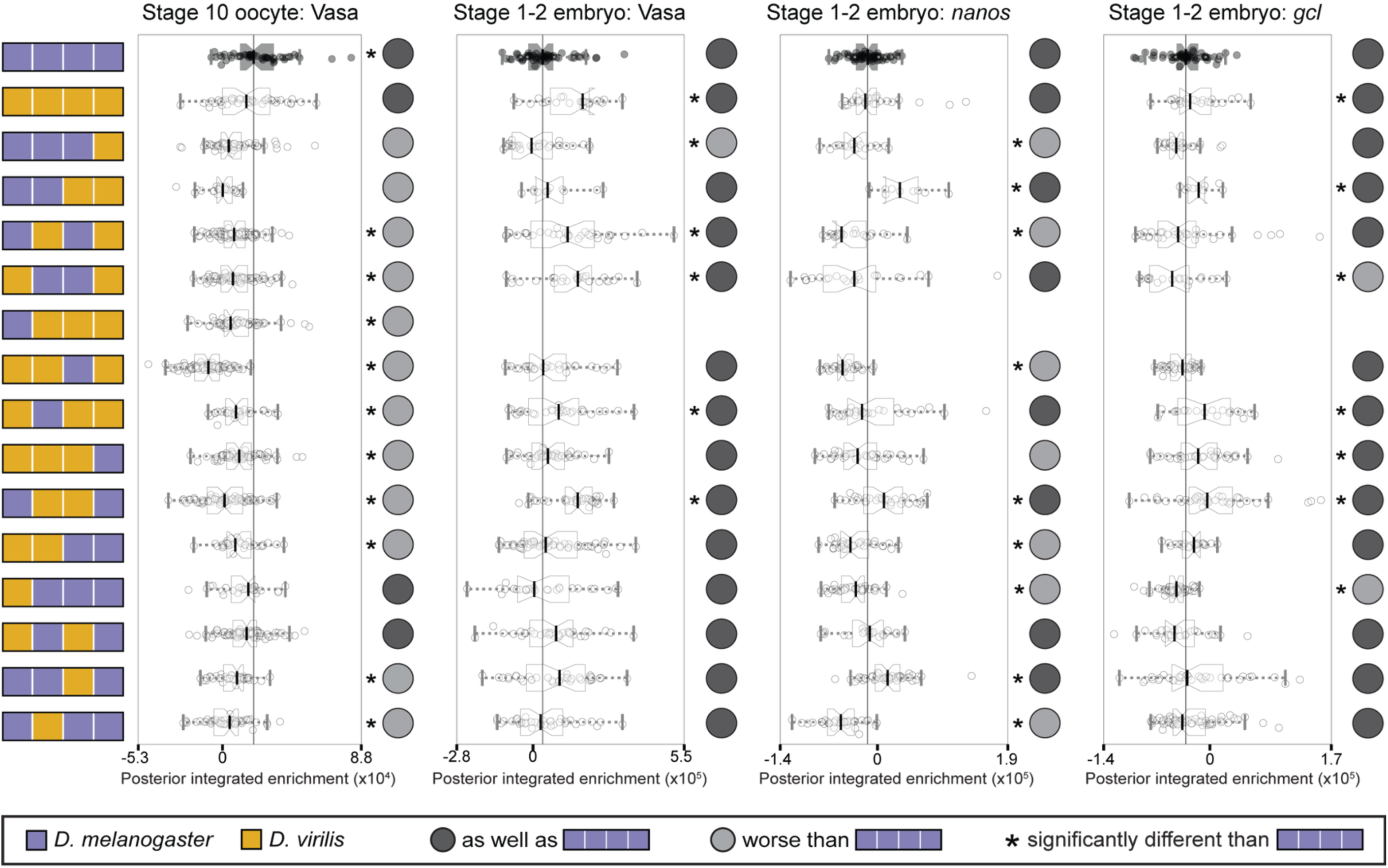
*D. melanogaster–D. virilis oskar* chimeras effectively recruit Vasa protein but not germ plasm mRNAs in *D. melanogaster oskar* null embryos. Posterior integrated enrichment of Vasa, *nanos*, and *gcl* in stage 10 oocytes and stage 1–2 embryos from *D. melanogaster* mothers expressing chimeric Oskar proteins. For the chimeric genotype with the Long Osk domain from *D. melanogaster* and the other three domains from *D. virilis*, sufficient posterior HA enrichment was found in oocytes to pass our threshold for analysis of germ plasm enrichment, but no embryos had sufficient enrichment to be included in the analysis. Distribution for eggs from flies expressing D*. melanogaster oskar* (positive control) is in black with the median marked with a black line. Significant difference from positive control distribution (p-value < 0.05, estimated from the distribution of difference of means between sample and control from 100,000 simulated samples) indicated by asterisk. Circles to the right of the plots indicate data interpretation as follows: dark gray = enriches the germ plasm molecule at least as well as *D. melanogaster oskar*; light gray = enriches the germ plasm molecule worse than *D. melanogaster oskar*. Sample sizes ý9 for each condition; see Table S3.

**Figure 4.**
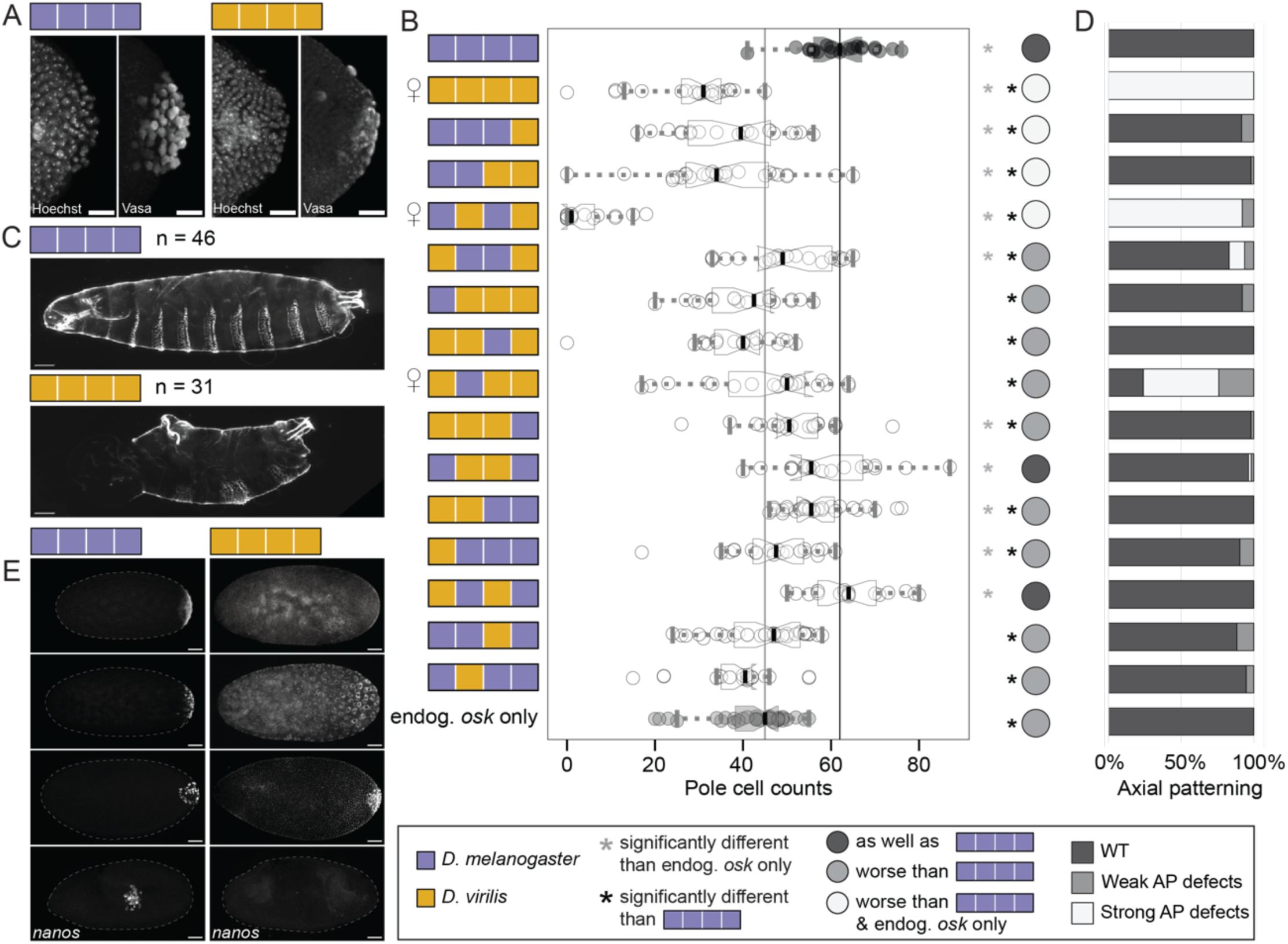
*D. virilis* Oskar and several Oskar chimeras disrupt axial patterning and impair germ cell formation in the presence of endogenous *oskar*. A) Stage 5 embryos from *D. melanogaster* mothers *expressing D. melanogaster* (purple, left) or *D. virilis oskar* (gold, right). Pole cells are marked by anti-*D. melanogaster* Vasa antibody and nuclei are marked with Hoechst. Micrographs are maximum intensity projections of 60 optical sections (step size = 1 *μ*m) and scale bars = 20 *μ*m. **B)** Pole cell counts in stage 5 embryos expressing chimeric *oskar*. Distribution from flies expressing only endogenous *oskar* (endog. *osk* only, negative control) is in gray with the median marked with a gray line. Distribution from flies expressing *D. melanogaster oskar* (positive control) is in black with the median marked with a black line. Significant difference from positive and negative control distributions (p-value < 0.05, estimated from the distribution of difference of means between sample and control from 100,000 simulated samples) indicated by black or gray asterisks respectively. Circles to the right of the plots indicate data interpretation as follows: dark gray = specifies pole cells as well as *D. melanogaster oskar*; light gray = specifies pole cells worse than *D. melanogaster oskar*; white = specifies pole cells worse than both *D. melanogaster oskar* and endogenous *oskar* alone. Sample sizes ≥20 for each genotype; see Table S4. Mirror of Venus symbol to the left of the plot indicates chimeric genotypes that exhibited dominant female sterility. **C)** Cuticles of embryos/first instar larval progeny of *D. melanogaster* mothers expressing *D. melanogaster oskar* (left) have wild-type anterior patterning, while the cuticles of *D. melanogaster* animals whose mothers expressed *D. virilis oskar* (right) are bicaudal. Micrographs are projections of 15 optical sections (step size = 1.47 *μ*m) and scale bars = 50 *μ*m. **D)** Frequency of defects in anterior-posterior (AP) patterning in progeny from mothers expressing each chimeric Oskar protein. Dark gray (WT) = wild-type AP patterning; light gray = weak AP defects: some duplication of posterior segments in the anterior but not completely bicaudal; white = strong AP defects: bicaudal or complete mirror image duplications of the posterior. Sample sizes ≥31 for each genotype; see Table S4. **E**) Representative examples of the range of *nanos* localization patterns observed in wild-type *D. melanogaster* embryos expressing either *D. melanogaster* (left) or *D. virilis oskar* (right). Micrographs are displayed with increasingly advanced stages of embryogenesis going top to bottom. Dotted lines outline the outer edge of embryos. Micrographs are maximum intensity projections of 30 optical sections (step size = 2 *μ*m) and scale bars = 50 *μ*m. Posterior is to the right in all micrographs.

**Figure 5.**
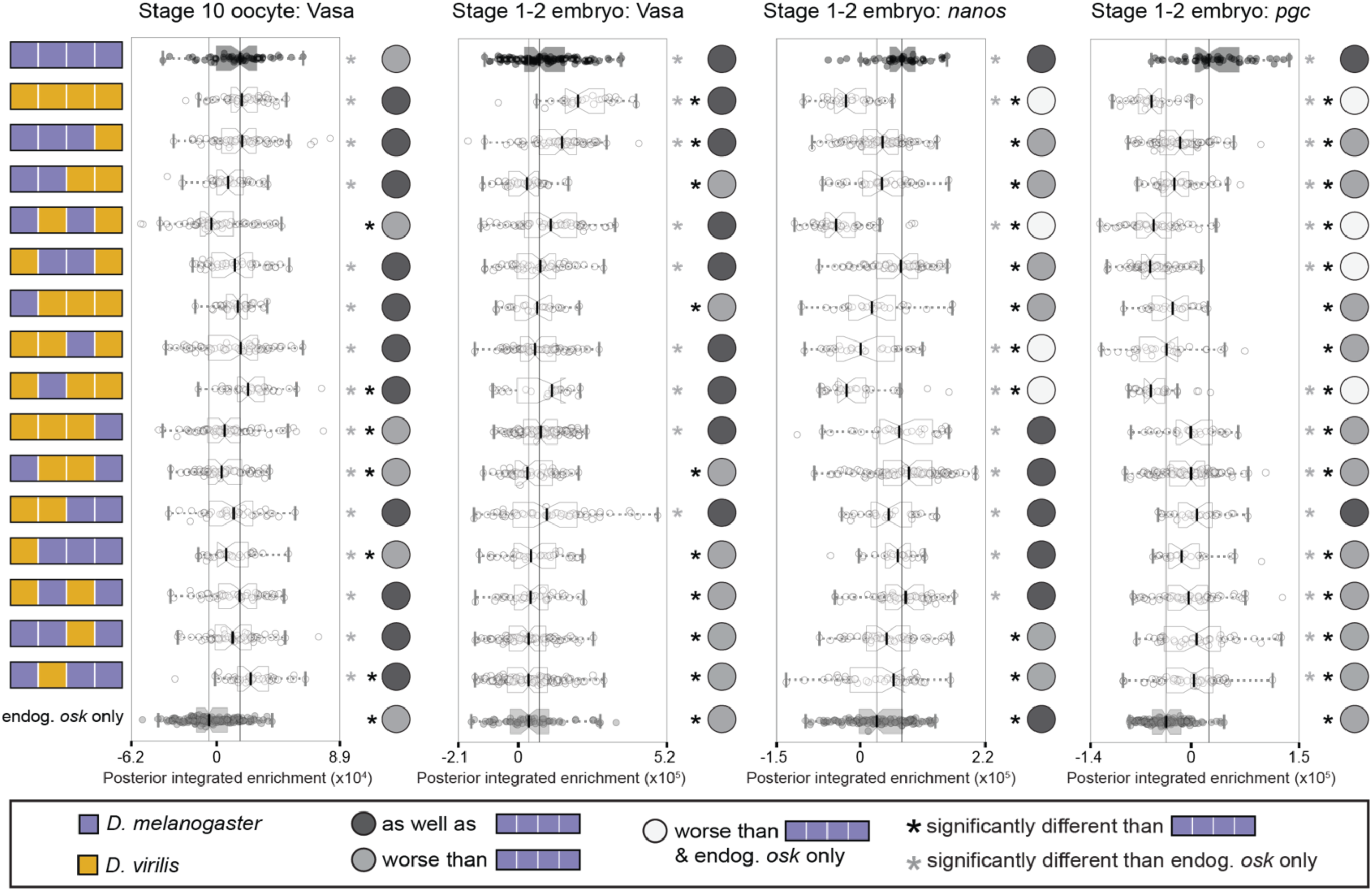
*D. melanogaster–D. virilis oskar* chimeras recruit Vasa protein but have dominant-negative effects on germ plasm mRNA localization in the presence of endogenous Oskar. Distributions of posterior integrated enrichment for Vasa, *nanos*, and *pgc* in stage 10 oocytes and stage 1–2 embryos from *D. melanogaster* mothers expressing chimeric *oskar*. Distribution from flies expressing *D. melanogaster oskar* (positive control) is in black with the median marked with a black line. Distribution from flies expressing only endogenous *oskar* (endog. *osk* only, negative control) is in gray with the median marked with a gray line. Significant difference from positive and negative control distributions (p-value < 0.05, estimated from the distribution of difference of means between sample and control from 100,000 simulated samples) indicated by black or gray asterisks respectively. Circles to the right of the plots indicate data interpretation as follows: dark gray = enriches the germ plasm molecule at least as well as *D. melanogaster oskar*; light gray = enriches the germ plasm molecule worse than *D. melanogaster oskar*; white = enriches Vasa worse than both *D. melanogaster oskar* and endogenous *oskar* alone. Sample sizes ≥16 for each condition; see Table S5.

**Figure 6.**
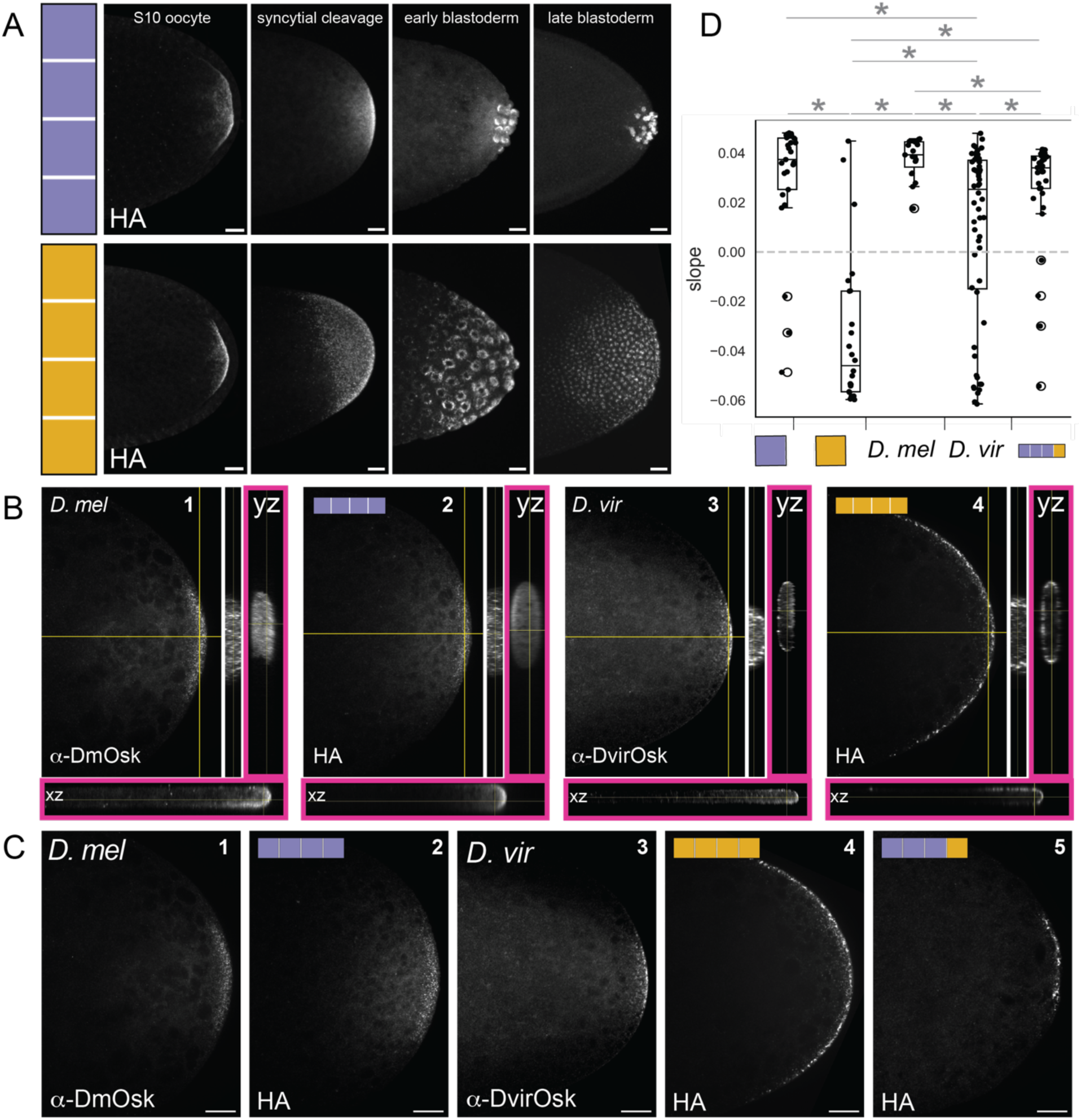
The OSK domain contributes to regulating species-specific granule size and localization. **A)** Oocytes and embryos from *D. melanogaster* mothers expressing *D. melanogaster* (purple, top row) or *D. virilis oskar* (gold, bottom row). Images are oriented with posterior to the right and are shown in order of increasing developmental time going left to right. Anti-HA antibody staining marks the localization of Oskar protein. Micrographs are maximum intensity projections of 30 optical sections (step size = 2 *μ*m) and scale bars = 20 *μ*m. **B)** Cleavage stage embryos of wild-type *D. melanogaster* (1: Oregon R) and *D. virilis* (3) stained with species-specific anti-Oskar antibodies, and *D. melanogaster* embryos from mothers expressing *D. melanogaster oskar* (2) or *D. virilis oskar* (4) stained with anti-HA antibody. Orthogonal views in the yz and xz orientations flank each image. Main images and orthogonal projections immediately to the right were captured with a 60x objective; orthogonal views bounded in pink were captured with a 20x objective, included here to more clearly visualize the distinct cortical localization of *D. virilis* and *D. melanogaster* Oskar proteins. **C)** Posterior pole of cleavage stage embryos from Oregon R (1), wild-type *D. virilis* (4), *D. melanogaster* expressing *D. melanogaster oskar* (2), *D. melanogaster* expressing *D. virilis oskar* (4), and *D. melanogaster* (bottom right) expressing *D. melanogaster oskar* with the OSK domain swapped with *D. virilis* sequence (5). *D. melanogaster* and *D. virilis* embryos are stained with species-specific Oskar antibodies and transgenic animals are stained for HA. Micrographs are single optical sections and scale bars = 20 *μ*m. Posterior is to the right in all micrographs. **D)** Plot of the slopes for lines of best fit that were drawn for curves of the fluorescent signal intensity when eroding yz projections of HA signal in embryos (see Fig. S13). Positive slope indicates more signal towards the middle of the embryo than the edges, and negative slope indicates more signal at the edges of the embryo. Significant difference in distributions (p-value < 0.05, estimated from the distribution of difference of means between sample and control from 100,000 simulated samples) indicated by asterisk; n.s. = not significant.

The germ line protein Vasa requires Oskar for its posterior localization ^56,57^ and directly interacts with *D. melanogaster* Oskar via the LOTUS domain based both on crystal structures and on *in vitro* assays ^27,28^. We wished to determine whether failure to accumulate sufficient Vasa in the germ plasm could account for the inability of some of the tested chimeras to induce pole cell formation. In stage 10 oocytes, we found no significant difference in Vasa posterior enrichment between animals expressing *D. melanogaster* or *D. virilis oskar* (Figs. 3, S5A, Table S3). In contrast, for most chimeras we observed reduced Vasa posterior enrichment in oocytes (Fig. 3), suggesting that combinations of sequences from the two species have a deleterious effect on initial recruitment of Vasa to the posterior of the oocyte. However, in cleavage stage embryos, we found that nearly all chimeras enriched Vasa at least as well as *D. melanogaster* Oskar, suggesting that sufficient Vasa protein can accumulate with additional time (Figs. 3, S5B, Table S3). Indeed, embryos expressing several chimeras, as well as full length *D. virilis oskar,* had significantly higher Vasa enrichment at the posterior than that observed with *D. melanogaster oskar* (Figs. 3, S5B, Table S3). Furthermore, we found that nearly all chimeras had correlation coefficients at least as high as those observed between *D. melanogaster* Oskar and Vasa in embryos (Fig. S7). Structural predictions of *D. virilis* Oskar protein interacting with *D. melanogaster* Vasa generated with AlphaFold3 ^49^ suggest these proteins would form the same interface observed in both predicted and crystallography-resolved structures of *D. melanogaster* Oskar LOTUS in complex with the RecA-like domain of Vasa (Fig. S8A) ^27^. These results suggest that failure to interact with Vasa is unlikely to explain why *D. virilis oskar* cannot rescue pole cell formation in a *D. melanogaster oskar* null background.

We then examined the distribution of the germ plasm mRNAs *nanos* and *germ cell-less* (*gcl*), suggested based on *in vitro* evidence to directly interact with the OSK domain of Oskar as a mechanism for their localization to the posterior ^24,25^. Despite the inability of most chimeras to form pole cells (Fig. 2), and the known role of *gcl* in pole cell formation ^58^, we did not find evidence of reductions in posterior enrichment of *gcl* in embryos from mothers expressing most of the chimeras (Figs. 3, S5D, Table S3). However, we did find significantly reduced enrichment of *nanos* in embryos laid by mothers expressing several chimeras with one or both of the *D. virilis* LOTUS or OSK domains (Figs. 3, S5C, Table S3). Thus, *D. virilis* protein sequence from either of the two folded Oskar domains may prevent *nanos* mRNA from accumulating at the posterior to wild-type levels in *D. melanogaster*.

Furthermore, we found that the *D. virilis oskar* line showed significantly reduced correlation between HA and *nanos* (Fig. S7, Table S3), suggesting that *D. virilis* Oskar may not interact with *nanos* mRNA in the same way as *D. melanogaster* Oskar. All chimeras with the *D. virilis* OSK domain also showed this reduced co-localization, as well as most chimeras with the *D. virilis* LOTUS domain, and the chimera with both folded domains from *D. melanogaster* but the two intrinsically disordered domains from *D. virilis* (Fig. S7, Table S3).

The OSK domain has long been considered a putative mRNA binding domain based on *in vitro* experiments ^24,25^, and an *oskar* allele lacking the OSK domain entirely (ΔOSK) expressed at the anterior pole of fly embryos fails to recruit germ plasm mRNAs ^26,30^. To generate hypotheses about the potential RNA-binding ability of this domain in each species, we generated electrostatic maps of the OSK domain (Fig. S8B). These maps suggest that the *D. virilis* OSK domain is less positively charged on the surface than the *D. melanogaster* OSK domain, perhaps indicating reduced RNA-binding ability. However, swaps of the OSK domain alone were not the only chimeric features that led to reduced mRNA enrichment or co-localization, again highlighting the complex and potentially not strictly modular nature of Oskar’s molecular interactions.

Together, these data suggest that failure to recruit Vasa is not responsible for the lack of germ cells in *D. virilis oskar* rescue experiments. Instead, differential interactions with germ plasm mRNAs may contribute to functional differences in Oskar protein between *D. virilis* and *D. melanogaster*.

### D. virilis Oskar has dominant-negative effects on D. melanogaster Oskar function

Oskar-mediated germ plasm assembly requires multiple complex molecular interactions, both homotypic ^26^ and heterotypic ^59^. We therefore hypothesized that the presence of *D. virilis* Oskar and/or chimeric Oskar proteins might disrupt the germ plasm assembly abilities of endogenous *D. melanogaster* Oskar. To test this hypothesis, we introduced chimeric Oskar proteins into a genetic background with endogenous *oskar*.

We first assessed the impact of these chimeras on wild-type germ plasm function by counting the number of pole cells specified. While over-expressing *D. melanogaster oskar* at the posterior led to significantly more pole cells compared to wild-type controls (expressing only endogenous *oskar*)*, D. virilis oskar* caused fewer pole cells to be specified by embryonic stage 5 (Fig. 4A–B, Table S4). All but two chimeras caused reductions in pole cell number compared to *D. melanogaster oskar*, suggesting that all of Oskar’s domains can contribute to pole cell formation in this background (Fig. 4A–B, Table S4). Notably, all chimeras containing the *D. virilis* OSK domain had significantly fewer pole cells than the *D. melanogaster oskar* positive control, and half of these chimeras exhibited dominant-negative effects on endogenous Oskar’s ability to specify pole cells (Fig. 4B, Table S4). In particular, the chimera with just the LOTUS and OSK domains from *D. virilis* generated significantly fewer pole cells than any other chimera, and even fewer than full length *D. virilis* Oskar (Fig. 4A–B, Table S4), suggesting that this combination of domains has a particularly potent negative effect on endogenous *oskar*’s function.

Strikingly, expression of *D. virilis oskar* or of two of the chimeric Oskar proteins in the presence of *D. melanogaster oskar* caused dominant female sterility (Fig. 4B). This finding is consistent with previous work showing sterility in *D. melanogaster* lines expressing full-length *D. virilis oskar* ^42^. We collected unhatched first instar larvae produced by females expressing *D. virilis oskar* and found that these embryos were bicaudal ^60^, a severe patterning defect in which the head and thoracic segments are replaced by a mirror image duplication of the posterior abdominal segments and telson (Fig. 4C). Examination of the chimeric *oskar* genotypes revealed three additional chimeras with these strong anterior defects (Fig. 4D, Table S4): the chimera with LOTUS and OSK from *D. virilis,* the chimera with all domains from *D. virilis* except LOTUS, and the chimera with OSK and Long Osk from *D. virilis.* We observed low, but not statistically significant, levels of partial posterior duplications in eight other chimeric genotypes, three of which contained the *D. virilis* OSK domain (Fig. 4D, Table S4).

Bicaudal phenotypes can result from deficient or insufficient posterior localization of multiple determinant molecules ^61–65^ , including the germ plasm component *nanos* mRNA ^66^. We therefore hypothesized that *nanos* mRNA might be mislocalized in the embryos derived from mothers expressing chimeras that yielded bicaudal larvae, thus causing the patterning defects described above (Fig. 4C–D). To test this, we compared the localization of *nanos* mRNA in embryos derived from mothers expressing either *D. virilis* or *D. melanogaster oskar* (Fig. 4E).

When expressing *D. melanogaster oskar*, *nanos* was tightly localized at the posterior in cleavage stage embryos, and was specifically detected in pole cells later in embryogenesis (Fig. 4E). In contrast, *nanos* was mislocalized in *D. virilis oskar* embryos. Specifically, there was reduced *nanos* enrichment at the posterior of preblastoderm embryos, low levels of *nanos* expression throughout the blastoderm, and loss of *nanos* expression in germ cells later in embryogenesis (Fig. 4E).

We then examined the localization of several additional germ plasm molecules recruited in the presence of Oskar chimeras in a background with endogenous *D. melanogaster oskar*. We noted that, in contrast to our observations in an *oskar* null background (Fig. 2), most chimeras were able to produce at least some pole cells in a background with endogenous *D. melanogaster oskar* (Fig 4). In addition to Vasa protein and *nanos* mRNA, we therefore chose to quantify the mRNA of germ plasm component *pgc*, rather than that of *gcl*, which we had chosen for the null background experiments based on its known role in pole cell formation ^58^. As in the *oskar* null rescue experiments (Fig. 3), we found that both *D. melanogaster* Oskar and *D. virilis* Oskar increased Vasa enrichment at the posterior in stage 10 oocytes and stage 1–2 embryos (Figs. 5, S6A–B, Table S5). However, *D. virilis* Oskar failed to localize the germ plasm mRNAs *nanos* and *pgc* to the posterior as well as *D. melanogaster* Oskar (Figs. 5, S6C–D, Table S5). In fact, enrichment of both *nanos* and *pgc* mRNAs was significantly reduced by the expression of *D. virilis* Oskar relative to enrichment in animals only expressing endogenous *oskar*. We therefore hypothesize that expression of *D. virilis oskar* leads to dominant-negative effects on *D. melanogaster oskar*’s ability to recruit germ plasm mRNAs. Chimeras containing the *D. virilis* OSK domain also exhibited significant defects in both *nanos* and *pgc* mRNA enrichment at the posterior, and half of these chimeras had reduced *nanos* and *pgc* enrichment at the posterior relative to endogenous *oskar* alone (Figs. 5, S6C–D, Table S5). These observations may implicate the OSK domain specifically in *D. virilis* Oskar’s possible dominant-negative effect on *D. melanogaster* Oskar-mediated germ plasm assembly. In addition, the finding that *D. virilis oskar* cannot enrich germ plasm mRNAs at the posterior is consistent with the observed strong axial patterning defect (Fig. 4C–D) and may contribute to the reduction in pole cells (Fig. 4A– B).

We wished to gain further insight into the potential mechanisms underlying the dominant-negative patterning defect. To this end, we quantified the accumulation and localization of chimeric Oskar protein, endogenous Oskar protein, and Nanos protein in wild-type oocytes and embryos (containing endogenous *oskar*) expressing full length *D. virilis oskar*, or expressing chimeras containing the *D. virilis* OSK domain. In these conditions, we detected at least as much chimeric or *D. virilis* Oskar protein at the posterior, as in controls (Figs. S3, S9, S10; Tables S6, S7). Moreover, the localization of the chimeric or *D. virilis* Oskar proteins closely matched that of endogenous Oskar protein, being tightly localized to the posterior in both stage 10 oocytes and embryos (Figs. S9, S10). As noted above, chimeric *oskar* mRNAs, including chimera Dm123Dv4 containing the *D. virilis* OSK domain, also localized correctly to the oocyte posterior (Fig. S4, Table S1). This suggests that chimeric *oskars* were translated and localized correctly, and that the observed patterning defects are unlikely to be due to insufficient quantity or localization of the chimeric Oskar proteins.

Endogenous Oskar protein was enriched at the posterior of stage 10 oocytes to at least the same degree as in controls (Fig. S9; Table S6). The posterior localization of endogenous Oskar in stage 10 oocytes in the presence of these chimeras was also similar to controls (Fig. S9). By stages 1–2 of embryogenesis, posterior enrichment of endogenous Oskar was statistically indistinguishable from controls in flies containing both endogenous *D. melanogaster* Oskar and full length *D. virilis* Oskar, as well as in flies containing both endogenous *D. melanogaster* Oskar and chimera Dm13Dv24 (Fig. S9; Table S6). This chimera, which contains the *D. virilis* LOTUS and OSK domains, induced the highest proportion (92%) of dominant-negative bicaudal defects, of all chimeric Oskar proteins (Fig. 4D). The second highest proportion (52%) of dominant-negative bicaudal defects was induced by chimera Dm2Dv134, which contains the *D. virilis* Long Oskar, linker and OSK domains (Fig. 4D). Stage 1–2 embryos containing this chimera as well as endogenous *D. melanogaster oskar*, had significantly less endogenous Oskar protein than controls, even though they had contained significantly more endogenous Oskar when they were stage 10 oocytes (Fig. S9; Table S6). Based on these observations, it could be the case that reduction of endogenous Oskar at the posterior could contribute to the dominant-negative AP patterning defects observed in the presence of chimera Dm2Dv134.

As noted above, full-length *D. virilis oskar*, and chimeras containing the *D. virilis* OSK domain, were unable to accumulate normal levels of *nanos* mRNA at the posterior in a wild-type background (Figs. 5, S10, Table S7). We examined the quantities and localization of Nanos protein in these genotypes, and found that Nanos protein enrichment did not always scale with *nanos* mRNA enrichment in a simple way. Specifically, we detected at least as much Nanos protein at the posterior as in controls, in the presence either of full-length *D. virilis* Oskar, or of chimeras containing the *D. virilis* OSK domain (Fig. S10B; Table S7). In accordance with our observations that ectopic *nanos* mRNA was sometimes detectable anterior to the normal posterior localization site (Fig. 4E), we detected similar regions of ectopic Nanos protein further anterior than in controls, especially at earlier embryonic stages (Fig. S10J). Further, we detected higher levels of Nanos protein at more anterior positions than in controls, in embryos with both endogenous *D. melanogaster oskar* and either full length *D. virilis* Oskar or chimeras containing the *D. virilis* OSK domain (Fig. S10I; Table S7). We observed strong colocalization of these chimeric proteins and *nanos* mRNA (Fig. S10D, J), but weaker colocalization of these chimeras and Nanos protein (Fig. S10E, J). We did not find evidence for mislocalization or absence of Smaug protein in embryos expressing *D. virilis* OSK-containing chimeras in a background with endogenous *D. melanogaster oskar*, suggesting that the observed patterning defects were not explained by the absence of Smaug-dependent degradation of repression of *nanos* (Fig. S11; Table S8). Taken together, these observations are consistent with the interpretation that inappropriately anteriorly localized Nanos protein contributed to the dominant-negative bicaudal patterning defects observed with these chimeras.

In *D. melanogaster*, Oskar is involved in a positive feedback loop between polarity maintenance, Oskar mRNA localization, and Oskar protein translation ^67,68^. We therefore asked whether defects in polarity maintenance might contribute to some of the apparent functional failures of chimeric Oskar. To do this, we assessed the behavior of the Par-1 protein, a kinase with a conserved role in eukaryotic cell polarity ^69^, which is normally enriched at the posterior pole of *D. melanogaster* oocytes and contributes to Oskar localization ^70–76^. We quantified the enrichment of Par-1 protein, its co-localization with chimeric Oskar, and its localization pattern in *oskar* null oocytes expressing selected Oskar chimeras, as well as in wild-type oocytes (containing endogenous *D. melanogaster* Oskar) expressing either full length *D. melanogaster oskar*, or full length *D. virilis oskar*. We found that in a wild-type background, there were no significant alterations to Par-1 enrichment, co-localization with Oskar, or localization pattern in the presence of *D. virilis* Oskar (Fig. S12D–E, Table S9). Par-1 also appeared unaffected in *oskar* null oocytes expressing chimeras containing the *D virilis* LOTUS or interdomains (Fig. S12A–C, Table S10). Interestingly, however, we noted that *oskar* null oocytes expressing chimeras containing the *D. virilis* Long Oskar or OSK domains did not accumulate as much Par-1 at the posterior as controls (Fig. S12B, Table S10). However, in *oskar* null embryos, four out the seven chimeras containing the *D. virilis* Long Oskar domain were able to support pole cell formation (Fig. 2), and all of them showed at least as much Vasa enrichment as controls (Fig. 3). Further, although none of the seven chimeras containing the *D. virilis* OSK domain were able to support pole cell formation in *oskar* null embryos (Fig. 2), most of them nevertheless accumulated appropriate levels of Vasa protein, *nos* and *gcl* mRNAs (Fig. 3). Moreover, we did not observe defects in *oskar* mRNA or protein localization in the chimeras we examined (Figs. S4, S9). Given these data, we cannot exclude the interpretation that some of the germ plasm and patterning defects we report here, may be partially caused by a disruption of the positive feedback loop that maintains oocyte polarity ^67,68^. However, we speculate that in many cases, even if polarity is initially disrupted in oocyte stages, this defect is corrected by the time of embryonic germ plasm accumulation.

### D. virilis Oskar nucleates granules with distinct localization and size compared to those nucleated by D. melanogaster Oskar

In addition to regulating germ plasm assembly and localization ^59^, previous reports suggest that Oskar also regulates the specific morphology and localization of germ granules ^41^. Germ granule morphology, including size, shape and localization, are strikingly variable across insects ^77,78^ including within the genus *Drosophila* ^79–82^, but the molecular mechanistic basis of Oskar’s regulation of these granule properties remains unknown. Previous histological reports described *D. virilis* germ plasm as having fewer, larger granules than *D. melanogaster* ^79^. We developed an antibody against *D. virilis* Oskar, and then used species-specific anti-Oskar antibodies to compare the localization pattern of wild-type *D. virilis* and *D. melanogaster* Oskar in their native context. We found that endogenous *D. melanogaster* Oskar in wild-type *Oregon R* embryos was distributed through the posterior cytoplasm (Fig. 6B–C, Table S11), similar to the distribution of electron-dense germ plasm granules reported for this species ^80,83^. However, in its native context *D. virilis* Oskar was significantly more enriched in the posterior cortical cytoplasm of the embryo, and appeared to form fewer, larger granules, than those observed for *D. melanogaster* Oskar (Fig. 6B–C, Table S11), again consistent with previous observations ^79^.

We then compared the localization of *D. melanogaster* Oskar and *D. virilis* Oskar expressed in *D. melanogaster oskar* null (*y[1] sc[*] v[1]]; Kr[If-1]/CyO; TI{w[+mC]=TI}osk[0.EGFP]*) oocytes and embryos by imaging embryos stained for HA, the tag we used for our chimeric *oskar* constructs (Fig. 6A). In mid-oogenesis (stage 10), *D. melanogaster* and *D. virilis* Oskar had indistinguishable HA localization patterns, suggesting that *D. virilis oskar* mRNA was successfully localized and then translated at the posterior (Fig. 6A). However, in cleavage stage embryos, *D. virilis* Oskar was detected over a broader region of the embryo than *D. melanogaster* Oskar (Fig. 6A). In addition, we saw *D. virilis* Oskar protein accumulation far from the posterior in apparently random ectopic locations in 36% of embryos (Fig. S13A–B). In the cellular blastoderm, *D. melanogaster* Oskar was localized to the pole cells, while *D. virilis* Oskar was detected in the cytoplasm of presumably somatic blastoderm cells at the posterior end of the embryo (Fig. 6A).

Examining the granular distribution of *D. virilis* Oskar within the posterior of *D. melanogaster* oocytes and early embryos revealed a fascinating pattern. In stage 10 oocytes, *D. melanogaster* and *D. virilis* Oskar localized similarly along the edge of the *D. melanogaster* oocyte membrane (Fig. S13D). However, we observed a difference in the HA signal distribution between species in cleavage stage embryos. *D. virilis* Oskar was tightly associated with the posterior embryonic membrane at this stage, while the signal for *D. melanogaster* Oskar was distributed throughout the posterior cytoplasm (Fig. 6B–C, Fig. S13C).

Finally, we saw gross morphological differences in the Oskar-containing granules of *D. melanogaster* expressing *D. virilis* or *D. melanogaster* Oskar. Specifically, HA signal appeared in fewer, larger foci when expressing *D. virilis* Oskar compared to the numerous small puncta of *D. melanogaster* Oskar-containing granules (Fig. 6D). Chimeric Oskar proteins possessing the OSK domain from *D. virilis* formed granules resembling, but not identical to, those formed by *D. virilis* Oskar (Fig. 6D), In other words, granules generated in *D. melanogaster* by *D. virilis* Oskar protein more closely resembled those of wild-type *D. virilis* than those of *D. melanogaster* (Fig. 6D). Further, granules generated by chimeric Oskar proteins containing the *D. virilis* OSK domain were also similar to the *D. virilis* granule type, suggesting that evolved changes to this domain may drive species-specific granule size and localization (Fig. 6D).

### The potential for evolved changes in Oskar protein to influence mRNA or protein interactions

We found that *D. virilis* Oskar successfully enriched the germ plasm protein Vasa at the posterior of embryos but exhibited defects in its recruitment of some, but not all, germ plasm mRNAs (Fig. 3; Fig. S2). This result led us to consider whether Vasa and germ plasm mRNAs might compete for Oskar binding sites. To explore this possibility *in silico*, we compared AlphaFold3 ^49^ predictions of *D. melanogaster* or *D. virilis* Oskar in complex with *D. melanogaster* Vasa, and with the *D. melanogaster nanos* 3’UTR (Fig. S8C–D). We found that regions of the *D. melanogaster* OSK domain that potentially interact with the *nanos* 3’UTR were not predicted to do so in *D. virilis* Oskar (Fig. S8C–D).

As a second *in silico* approach to asking whether evolved changes in RNA interacting ability might explain the differences between *D. virilis* and *D. melanogaster* proteins *in vivo*, we used GPSite ^84^, a machine-learning model for predicting protein binding sites with various other molecules, to predict binding residues for *D. melanogaster* and *D. virilis* Oskar protein with protein and RNA. We considered residues with predicted binding scores greater than a threshold of 0.5 to be putative binding sites. We observed that while the overall pattern of predicted residue binding affinity was similar between the two orthologs, *D. virilis* Oskar exhibited higher protein-binding scores than *D. melanogaster* Oskar within the LOTUS domain, while *D. melanogaster* Oskar protein-binding affinity exhibited higher peaks than *D. virilis* within the interdomain (Fig. S14A, C). *D. melanogaster* Oskar exhibited higher RNA-binding scores than *D. virilis* Oskar throughout the protein (Figs. S14B, D, S15B). Indeed, *D. virilis* Oskar was not predicted to have any residues over the threshold for confident RNA-binding in its Long Oskar isoform, and only two residues over the threshold for its Short Oskar isoform, compared to the thirteen in *D. melanogaster* Short Oskar. (Fig. S14B, D).

We compared the overall protein-binding and RNA-binding scores for each of our chimeric Oskar protein sequences. We found that all chimeric Oskar proteins had overall protein-binding scores over the threshold, but chimeras with their LOTUS domain from *D. virilis* had consistently higher protein-binding scores (Fig. S15A, C). In terms of predicted RNA-binding ability, while *D. melanogaster* Oskar and several additional chimeras had RNA-binding scores over 0.5, *D. virilis* Oskar and a host of other chimeric proteins were below this threshold for the Long Oskar isoform (Fig. S14B, D). Intriguingly, the three chimeric proteins with the lowest RNA-binding scores for short Oskar exhibited patterning defects and severe defects in mRNA enrichment in our experiments conducted in a background with endogenous Oskar (Fig. S15D).

Our AlphaFold3 predictions suggested that the *D. virilis* OSK domain should exhibit a greater interaction with Vasa than the OSK domain of *D. melanogaster* (Fig. S8C–D). Using GPsite ^84^ to examine the relationship between RNA-binding and protein-binding scores, we found a clear pattern of *D. virilis* LOTUS leading to high protein-binding score, regardless of the other domains present (Fig. S15E, F). The predicted RNA-binding score was less tightly associated with any specific protein domain. However, the *D. virilis* OSK domain appeared in nearly all of the lowest ranked short Oskar chimeras (Fig. S15F). We notably observed that the short Oskar chimeras with all *D. melanogaster* and all *D. virilis* sequence were found in the most extreme corners of the correlation plot, with *D. melanogaster* Oskar displaying high RNA-binding and lower protein-binding scores, while *D. virilis* Oskar has high protein-binding and low RNA-binding scores (Fig. S15). Taken together, these *in silico* observations are consistent with our *in vivo* experiments in supporting the hypothesis that evolved differences in the OSK domain change this domain’s ability to interact with germ plasm mRNAs in species-specific ways, which are a major cause of interspecies incompatibility of Oskar’s germ line determinant role.

## Discussion

We have used domain swapping between orthologous genes to obtain insight into the evolutionary changes that can affect protein function, even between closely related species, as well as the functional roles of that protein’s domains in its native context. We uncovered the basis for the functional incompatibility of *D. virilis oskar* in *D. melanogaster* by expressing sixteen chimeric versions of *oskar* in *D. melanogaster* and testing their ability to perform *oskar* function (Fig. 1D). Our data suggest that the *D. virilis oskar* and chimeras containing the *D. virilis* OSK domain fail to rescue germ cell formation (Fig. 2A–B), but that all chimeras can successfully restore abdominal patterning in *oskar* null embryos (Fig. 2C). We observed a complex relationship between Oskar sequence and its ability to recruit germ plasm molecules. *D. virilis oskar* can successfully enrich Vasa at the posterior but exhibits defects in its interactions with some germ plasm mRNAs, and these defects are often associated with the presence of the *D. virilis* OSK domain (Fig. 3). In addition, *D. virilis oskar* has a dominant-negative effect on *D. melanogaster oskar* function that is also at least partially due to the *D. virilis* OSK domain (Figs. 4–5), which appears to lead to inappropriately anterior accumulation of Nanos protein (Fig. S10), resulting in patterning defects including bicaudal embryos (Fig. 4). Finally, we identify differences in the localization and size of *D. virilis* and *D. melanogaster* Oskar-containing granules that are linked at least partially to the presence of the OSK domain, and that we speculate may influence their ability to mediate downstream functions (Fig. 6). Thus, our results shed light on the evolution of a key germ line protein.

### Maintenance of Oskar at the posterior

In the first reported test of the function of a non-*D. melanogaster* oskar ortholog, Webster and colleagues ^42^ observed reduced maintenance of *D. virilis oskar* mRNA at the posterior of *D. melanogaster* embryos (their Fig. 3D) relative to *D. melanogaster oskar* mRNA. They also reported more diffuse localization of *D. melanogaster oskar* mRNA (their Fig. 3C) and protein (their Fig. 3J) when both *D. melanogaster* and *D. virilis oskar* were expressed in the same *D. melanogaster* embryo. The authors proposed a model for Oskar protein function in which one domain of Oskar anchors it to the posterior cortex, while one or more distinct domains bind to germ plasm molecules ^42^. They speculated that *D. virilis* Oskar could interact with *D. melanogaster* posterior patterning determinants but could not anchor itself to the posterior, thereby dragging these determinants away from the pole ^42^.

In the three decades since that work was published, we have learned that Oskar indeed has a domain with a distinct role in anchoring itself to the posterior, namely the Long Osk domain ^36^. However, our data do not suggest that swapping the *D. melanogaster* Long Osk domain with *D. virilis* Long Osk sequence leads to severe defects in the posterior maintenance of *oskar* or other germ plasm molecules (Fig. 3). In fact, Webster and colleagues ^42^ concluded their study by describing a *D. melanogaster–D. virilis oskar* chimeric protein that they expressed at the anterior pole of *D. melanogaster* embryos. This chimera was described as having 188 amino acids of N-terminal *D. melanogaster* sequence, which corresponds to the Long Osk domain and a portion of the LOTUS domain (Fig. S1B), with the remainder of the sequence derived from *D. virilis*. They reported (as “data not shown”) that that this chimera failed to specify pole cells at the anterior of the embryo ^42^. This finding is consistent with our results, which show that the chimera possessing only *D. melanogaster* Long Osk cannot specify pole cells in *D. melanogaster*, nor can most other chimeras that possess the *D. virilis* OSK domain (Fig. 2B). Together, these data suggest that *D. virilis* Oskar’s failure to localize tightly to the posterior in *D. melanogaster* is not solely based on a difference in the Long Osk domain itself. However, it is possible that changes to another domain within the Long Oskar isoform may influence its activity, given that Long Oskar contains all the domains found in the shorter isoform.

Alternatively, Short Oskar’s function in germ granule assembly may also affect Oskar localization at the posterior. For instance, Oskar failing to interact with germ plasm mRNA could in principle influence granule size or biophysical properties in a way that allows diffusion away from the posterior more readily. To our knowledge, no other domain of Oskar has been directly implicated in posterior localization, and further work will be needed to test this new hypothesis about how Oskar might be retained at the posterior pole.

### Oskar’s inferred interactions with germ granule components

Although the LOTUS domain is classically associated with Vasa interactions and the OSK domain with mRNA binding, *in vivo* studies challenge the strict modularity of Oskar’s domains, suggesting the LOTUS domain may also contribute to mRNA interactions ^26^. If Vasa and germ plasm mRNAs indeed interact with Oskar via some of the same residues, then reduced interaction with mRNA could in principle enable more interaction with Vasa. Conversely, greater affinity for Vasa could preclude interaction with mRNA. Our data cannot distinguish between these possibilities, but are consistent with the hypothesis that Oskar’s interactions with germ plasm molecules *in vivo* are complex and potentially non-modular in nature. Future work could examine the specific intermolecular interactions of cross-species germ plasm components using quantitative analysis of single-molecule imaging, or assessing RNA translation dynamics.

Recent work demonstrated that degrading germ plasm mRNAs increases the size of Oskar-labeled germ granules, suggesting a relationship between protein and mRNA incorporation in granules ^30^. We found that the Oskar-labeled granules from *D. virilis* appeared in larger foci than the small Oskar-labeled granules nucleated by *D. melanogaster* Oskar (Fig. 6D). We speculate that the reduced mRNA co-localization and increase in Vasa co-localization effected by *D. virilis* Oskar (Fig. S7) could contribute to the apparent difference in Oskar-labeled germ granule size.

The germ granules that form upon expression of *D. virilis* Oskar in *D. melanogaster* resemble those detectable in *D. virilis* using an anti-*D. virilis* Oskar antibody (Fig. 6D)*. D. virilis* germ granules are distinct from those of *D. melanogaster* both morphologically and in terms of molecular content. In her classic comparative study of *Drosophila* germ granules, Sheila Counce described *D. virilis* granules as fused into short strings, in contrast to the small, individual droplets observed in *D. melanogaster* ^79^. In addition, a recent study found that *D. virilis* exhibits lower expression levels of several germ plasm mRNAs and has fewer transcripts within granules than *D. melanogaster* ^44^. This difference in germ plasm mRNA abundance might explain why *D. virilis* embryos form fewer pole cells than *D. melanogaster* (Fig. 1B). It is also possible that variable germ plasm mRNA amount, or other differences in the composition of *D. virilis* germ granules relative to those of *D. melanogaster*, may explain the morphological differences in the granules between these species. Further examination of pole bud formation, pole cell divisions, and the contents and biophysical properties of germ granules in both species, will be needed to rigorously test this hypothesis.

The origin and functional consequences of the variation in germ granule morphology and composition across *Drosophila* species remain unclear ^44,79,80^. While previous work suggested that mRNA position within germ granules is independent of the onset of translation or degradation ^85^, a recent study found a correlation between the positioning of mRNAs within germ granules and their translational status ^86^. It is possible that species-specific differences in germ granule structure or morphology could influence the post-transcriptional regulation of germ granule components. In the case of the germ granules nucleated by *D. virilis* Oskar in *D. melanogaster*, possible changes in the proportions or spatial organization of germ plasm components within the granule could disrupt the onset of gene expression patterns that regulate pole cell formation and other precisely timed developmental processes.

### Different germ plasm requirements for pole cell specification and axial patterning

We found that *D. virilis* Oskar was able to rescue the abdominal patterning but not the pole cell specification defects of *D. melanogaster oskar* null embryos (Fig. 2), consistent with previous reports ^42^. This may be because *D. virilis* Oskar can interact sufficiently with abdominal patterning determinants but insufficiently with germ cell determinants in *D. melanogaster*. Alternatively, this finding may reflect a difference in the dose dependency or sensitivity of these processes to changes in posterior localization of germ plasm.

Oskar directs the localization of the posterior patterning determinant *nanos*, which establishes a posterior gradient of Nanos protein that inhibits translation of *hunchback* at the posterior and enables proper axial patterning ^87,88^. However, *nanos* is not required for germ cell specification ^66,89^, although it is required for their subsequent migration and survival ^90–93^. We observed no striking differences in the predicted interactions between *D. melanogaster* and *D. virilis* Oskar and the three germ plasm proteins Lasp, Valois, and Smaug (Figs. S11, S16), for which direct interactions have been proposed based on yeast two hybrid, GST-pull down and cell culture-based assays ^31–33^. It is possible that other, untested molecules required for germ cell specification fail to assemble in the *D. melanogaster* germ plasm with *D. virilis oskar* expression.

It may also be the case that pole cell formation requires more stringent posterior localization and a higher concentration of germ granule components than is necessary for *nanos* to mediate proper axial patterning. The role of germ granules in pole cell specification is tightly associated with the posterior pole, where granules mediate complex processes allowing pole buds to form, including changes in the phospholipid composition of the membrane and the actin cytoskeleton network ^94,95^. The species-specific differences in *D. virilis* Oskar localization at the posterior (Fig. 6) may impact the dynamic process transporting germ granules into the germ cells as they form ^96^. Future studies deploying high-resolution live imaging of embryos at the time of pole budding may offer new insights into how germ cell formation is disrupted upon expression of *D. virilis oskar*.

Jongens and colleagues ^97^ reported that ectopic expression of *gcl* at the anterior led to formation of ectopic anterior pole buds . This might initially appear incompatible with our observation that some chimeras with levels of *gcl* that were comparable to controls, were nonetheless unable to form pole cells (Figures 2, 3). However, Jongens and colleagues ^97^ noted that the ectopic pole buds formed by anteriorly localized *gcl* did not pinch off to form pole cells. This was consistent with their report that other germ plasm components did not ectopically localize to the anterior in response to anterior *gcl* expression (a report described in the text of their results section, but with data indicated as “not shown” ^page^ ^21^^28^ ^in^ ^97^. They interpreted their results to mean that “the level of *gcl* activity required for the induction of pole bud formation may be much less than that required to complete pole cell formation” ^page^ ^21^^32^ ^in^ ^97^ . We did not, as did Jongens and colleagues ^97^, quantify pole bud formation by examining embryos at stages earlier than the end of stage 5. Instead, we only included late stage 5 embryos with well-formed pole cells in our quantification (Fig. 2). Many chimeras that failed to drive pole cell formation despite apparently unaffected *gcl* levels, nevertheless showed alterations in the levels of at least some of the other germ plasm components that we quantified. Our observation is thus consistent with the interpretation of Jongens and colleagues, that “*gcl* activity alone is not sufficient for pole cell formation or maintenance ^page^ ^21^^33^ ^in^ ^97^. In sum, while many of the individual molecules found in wild-type germ plasm clearly play important roles in establishing some of the molecular and cellular characteristics of pole cells (e.g. *gcl* for pole bud formation, *nanos* for somatic state repression, *oskar* for granule nucleation), establishing robustly cellularized PGCs that can maintain germ cell fate, requires a complex interaction of many, if not all, of these components.

### *Dominant-negative effect of* D. virilis oskar *on* D. melanogaster oskar *function*

We observed that expression of *D. virilis oskar* in a background with endogenous *D. melanogaster oskar* had dominant-negative effects on axial patterning, pole cell number, and germ plasm mRNA enrichment (Fig. 4–5). In fact, the mRNA enrichment defects were more severe in a background with both *D. melanogaster* and *D. virilis oskar* than in an *oskar* null background, leading to development of bicaudal embryos, (Fig. 4–5) a phenotype which does not occur when expressing *D. virilis oskar* in *oskar* RNA null embryos (Fig. 2). There is precedent for dominant-negative effects arising from expression of a wild-type gene from another species in a non-native context. For example, expression of homologs of the *matrimony* gene from multiple *Drosophila* species in wild-type *D. melanogaster* embryos causes severe developmental defects ^98^. Furthermore, expression of the *D. melanogaster* Serum Response Factor ortholog (*blistered*) in mammalian cells has a dominant-negative effect on the activity of the endogenous mammalian factor, which appears to occur via dimerization between the endogenous and transfected proteins ^99^.

One possible molecular explanation for the strong dominant-negative effect of *D. virilis* Oskar could be the formation of heterodimers between *D. virilis* and *D. melanogaster* Oskar that have a deleterious effect on Oskar function. *D. melanogaster* Oskar’s LOTUS domain dimerizes in crystal structures ^24–26^, and AlphaFold3 ^49^ consistently predicts interactions between *D. melanogaster* Short Oskar monomers at the same interface observed in the reported crystal structures (Fig. S17A). We found that *D. virilis* Short Oskar is predicted to form the same dimerization interface observed in homodimers of *D. melanogaster* Short Oskar (Fig. S17B). Furthermore, a heterodimer forming through interaction at the LOTUS domain is also predicted between *D. melanogaster* and *D. virilis* Short Oskar (Fig. S17C). These predictions are consistent with the hypothesis that *D. melanogaster* and *D. virilis* may directly interact in the trans-species transgenic *D. melanogaster* lines studied herein.

Unlike the predicted homodimers, the heterodimer also exhibits moderately high confidence interactions between the OSK domains of *D. melanogaster* Oskar and *D. virilis* Oskar proteins (Fig. S17C). These OSK-OSK interactions are also apparent in higher-order predicted oligomeric structures containing both *D. melanogaster* and *D. virilis* full length Oskar (Fig. S17C). Further, when both Oskar orthologs are present, *D. melanogaster* Oskar is consistently predicted to heterodimerize with *D. virilis* rather than forming homodimers (Fig. S17C). We speculate that the predicted OSK-OSK interactions between *D. melanogaster* and *D. virilis* Oskar might preclude interactions between *D. melanogaster* germ plasm mRNA and the *D. virilis* Oskar and chimeric Oskars studied here. AlphaFold3 ^49^ predicts structures of *D. melanogaster* Oskar proteins with the *D. melanogaster nanos* 3’UTR that suggest multiple weak potential interactions between Oskar and the mRNA (Fig. S17D). Similar interactions are also predicted between *D. virilis* Oskar and the *D. melanogaster nanos* 3’UTR (Fig. S17E). However, these predicted interactions are lost in structures that include both *D. melanogaster* and *D. virilis* Oskar proteins with the *D. melanogaster nanos* 3’UTR. Instead, weak interactions appear between the OSK domains of *D. virilis* and *D. melanogaster* Oskar (Fig. S17F). These predictions further support the hypothesis that the dominant-negative effect of *D. virilis* Oskar in *D. melanogaster* arises through detrimental direct interactions with *D. melanogaster* Oskar. Future empirical tests of this hypothesis may provide new insights into the nature and functional significance of Oskar dimerization *in vivo*.

### Evolution of oskar isoforms and the Long Oskar domain

One element of Oskar function that our experiments cannot speak to, is the relative amounts of the Long and Short isoforms of Oskar. To date, the mechanisms that regulate the stoichiometry of these Oskar isoforms in *D. melanogaster*, and the dynamics and biological significance of the stoichiometry, remain unknown. The C-terminal HA-tagged design of our chimeric transgenes means we are unable to distinguish between chimeric Long and Short isoforms in our transgenic lines. It is possible that alteration of these ratios for one or both of endogenous or chimeric Oskars, contributes to the trans-species incompatibility that we explore here. We also note that the polyclonal antibody we raised against *D. virilis* Oskar cannot distinguish between the two isoforms. Consequently, we cannot comment on whether the *D. virilis* Long isoform, or chimeras containing the *D. virilis* Long Oskar domain, are localized cortically and/or to endosomes, as is the case for *D. melanogaster* Long Oskar ^36–38,100^.

However, some evidence suggests that *oskar* mRNAs and proteins from these two fly species do not have identical subcellular behaviors. We observed that *D. virilis* Oskar protein is more prone to localize closer to the embryonic cortex, than *D. melanogaster* Oskar, both in the endogenous *D. virilis* context and in the trans-species transgenic one. In addition, we (this study) and others ^42^ have observed that *D. virilis oskar* gene products may be less strongly restricted to the posterior at certain stages than *D. melanogaster* Oskar, both in the trans-species context of the *D. melanogaster* embryo (protein in Fig. 6A, mRNA in Webster and colleagues’ Figure 3 and described as “data not shown” in their Figure 4 ^42^) and in the endogenous context of the *D. virilis* oocyte (mRNA in Webster and colleagues’ Figure 2^42^). Similarly, transcripts of the *oskar* homologs of the jewel wasp *Nasonia vitripennis* and the ant *Messor pergandi* are also localized to the posterior of developing oocytes at early stages of oogenesis, but appear somewhat more diffuse than in *D. melanogaster* (Lynch and colleagues’ Figures 3 and 8 ^101^). These hymenopteran *oskar* sequences lack a Long Oskar domain ^101^, showing that despite its important role in germ plasm anchoring in *D. melanogaster*, this domain can be dispensable for functional germ plasm assembly in other species.. This could mean that the anchoring function demonstrated for the *D. melanogaster* Long Oskar domain is species-specific, and an evolutionary innovation relative to Oskar’s conserved germ plasm assembly role. Consistent with this hypothesis, we previously provided evidence that the wasp example is not an isolated case: the Long Oskar domain itself is entirely absent from Oskar proteins in most insects, and evolved only approximately 70 million years ago, around the time of divergence of the lineage leading to the Pinpunculidae (the big-headed flies) ^20^. This also indicates that evolving the Long Oskar domain is not what allowed *oskar*, which likely originally played a role in somatic development ^20,102,103^, to evolve its germ line role, since at least some *oskar* orthologs without the Long Oskar domain can still assemble germ plasm.

### Evolution-inspired analysis of protein structure-function relationships

Functional characterization of classical *oskar* alleles generated with random mutagenesis offered the first insights into the biological processes regulated by Oskar ^87,104^. More recently, targeted mutagenesis has been used to determine the functional significance of specific regions of the Oskar protein, identifying residues that might be important for interaction with Vasa ^27^ or mRNA ^25,26^. However, designing such targeted experiments inherently relies on assumptions about which residues might be good candidates to test. Here, we leveraged the naturally occurring sequence divergence associated with the functional incompatibility of *D. virilis* Oskar in *D. melanogaster* to identify functionally significant regions of Oskar *in vivo.* By taking advantage of species-specific sequence variations, we were able to evaluate functionally significant portions of Oskar in an unbiased manner. This type of approach has offered insight into other structure-function relationships. For instance, the pea aphid (*Acyrthosiphon pisum*) homolog of Vasa fails to localize to the germ plasm when expressed in *D. melanogaster* ^105^. By generating a series of chimeras swapping domains between *D. melanogaster* and *A. pisum* Vasa, researchers demonstrated that specific residues within the C-terminal HELICc domain of Vasa are required for posterior localization *in vivo* ^105^. Going forward, our work on Oskar may provide a foundation for targeted screens, for instance within the OSK domain, to identify specific residues important for pole cell formation. This approach provides a powerful framework for dissecting the molecular evolution of developmental genes and identifying functionally significant residues in rapidly evolving proteins.

## Acknowledgements

We thank members of the Extavour Lab for helpful discussions, particularly Chandrashekar Kuyyamudi for assistance with image analysis. We also thank several Extavour lab members for assistance with experiments required for revision of the manuscript: Tarun Kumar for fly crosses and maintenance; Chandrashekar Kuyyamudi for embryo collection, staining, and imaging; Macy Ingersoll for dissections and embryo collection; and Evelyn Schwager, Damien Markus, Yu Shirai, Suhrid Ghosh, Daniella Haber and Petra Kovacikova for embryo collection. We thank Mariana Wolfner and members of her lab for assistance with Western Blots. This study was supported by NIH award # 1R01GM143611, funds from Harvard University, and from the Howard Hughes Medical Institute awarded to CGE. Additional support was provided by the NSF-Simons Center for Mathematical and Statistical Analysis of Biology at Harvard (award number #1764269), the Harvard Quantitative Biology Initiative, a Boehringer Ingelheim Fonds Ph.D. fellowship awarded to A.R., and a Herchel Smith Graduate Fellowship awarded to ELR.

## Author Contributions

CGE, JRS, and ELR conceived and designed the project. ELR and JRS performed the experiments. AR provided custom scripts for image analysis. CGE supervised the project and acquired funding. ELR and CGE wrote the manuscript. All authors reviewed and edited the manuscript.

## Methods

### Generation of chimeric oskar constructs

We removed the UAS, promoter, and *k10* 3’UTR elements from pVALIUM22 [51-53] via round-the-world polymerase chain reaction (RTW-PCR), leaving the *gypsy* insulator elements. We inserted a 6,577 kb ApaI–XhoI *D. melanogaster* genomic digest fragment corresponding to the genomic *oskar* locus (GenBank Acc. No. AE014297, region 8932758-8939335; gift from Dr. Anne Ephrussi, European Molecular Biology Laboratory, Heidelberg, Germany) into this linearized and modified pVALIUM22 via Gibson assembly. This genomic locus has previously been shown to provide wild-type *oskar* function in a loss-of-function background ^34^. We introduced a C-terminal hemagglutinin (HA) epitope tag (TACCCATACGATGTTCCGGATTACGCT, encoding YPYDVPDA) via RTW-PCR, generating the plasmid pVAL22-Dm1234-HA. A clone of the *D. virilis osk* genomic locus ^42^ (GenBank Acc. No. L22556; obtained from Dr. Paul Macdonald, University of Texas at Austin, Texas, U.S.A.) served as the template for fragment amplification.

To generate various combinations of *D. melanogaster* and *D. virilis osk* fragments, we applied RTW-PCR to pVAL22-Dm1234-HA to delete a specific region, PCR to amplify the corresponding region from the *D. virilis osk* genomic locus clone, and Gibson assembly to insert the *D. virilis osk* fragment into the truncated pVAL22-Dm1234-HA plasmid. This strategy was applied to four specific regions in pVAL22-Dm1234-HA, corresponding to the Long Osk, LOTUS, interdomain, and OSK domains (Fig. 1D), which led to the creation of 16 total constructs. All transgenic constructs possess the *D. melanogaster oskar* 5’UTR and 3’UTR sequences.

### Drosophila stocks

We prepared plasmid midi preps containing each *oskar* chimeric construct (Qiagen HiSpeed Plasmid Midi Kit) and sent them to BestGene (Chino Hills, CA, USA) for injection into the attP40 line (*y[1] w[67c23]; P{CaryP}attP40*). We crossed P_0_ adults to *y[1] v[1]* to screen for transformants based on *vermillion* rescue, balanced constructs over *CyO,* and introduced the recessive marker *scute* to distinguish transgenic lines from wild-type animals (*y[1] sc[*] v[1]; chimeric-osk/CyO.*) Heterozygotes for the transgene were used in experiments testing the effects of one copy of chimeric *oskar* expression in a background with endogenous *oskar*. We used *y[1] sc[*] v[1]; +/CyO* animals as a negative control in these experiments (referred to as “endogenous *oskar*” in the text).

For rescue experiments, we generated animals expressing the chimeric *oskar* constructs in the *oskar* RNA null allele background *y[1] sc[*] v[1]; chimeric-osk/CyO; TI{w[+mC]=TI}osk[0.EGFP]* (*oskar* allele obtained from Bloomington *Drosophila* Stock Center 67407), which contains a 3xP3-EGFP-tubulin 3’UTR cassette inserted 26 nucleotides upstream of the *oskar* transcriptional start site that leads to failure to transcribe *oskar* ^55^ (*/TM3*). Animals heterozygous for the chimeric *oskar* construct and homozygous for *osk[0.EGFP]* were used in experiments testing the effect of one copy of chimeric *oskar* expression in an *oskar* null background.

For experiments with wild-type animals, we used the *D. melanogaster* strain *Oregon R* (Bloomington *Drosophila* Stock Center 5) and *D. virilis* (gift from Rainbow Transgenics, Camarillo, CA, USA).

All flies were reared at 25°C at 60% humidity with standard fly food and yeast as previously described ^106^.

### Immunofluorescence and in situ hybridization

For antibody staining experiments in oocytes, we dissected ovaries from 3–5 day old females in 1xPBS (phosphate buffered saline), fixed for 20 minutes in 4% PFA (paraformaldehyde) in 1xPBS, and washed with 1xPBS. We permeabilized tissue twice for five minutes in 1xPBT (1xPBS + 0.2% Tween-20), once for two hours in 1xPBTT (1xPBT + 1% Triton X-100), and once more for five minutes in 1xPBT. We blocked for two hours in 1xPBT + 1% BSA (bovine serum albumin, Millipore Sigma) and incubated with primary antibody overnight at 4°C (see list of antibodies and concentrations in Table S12 and Key Resources Table). The following day, we washed three times for one hour in 1xPBT + 1% BSA and blocked for one hour in 1xPBT + 1% BSA + 2% NGS (normal goat serum, Jackson ImmunoResearch). We mixed secondary antibody solution in 1xPBST + 1% BSA + 4% NGS and incubated overnight at 4°C. The following day, we washed five times for 15 minutes in 1xPBT, mounted samples in Vectashield (Vector Laboratories), and stored slides at –20°C until imaging.

For antibody staining experiments in embryos, we collected embryos on apple juice plates (18 g agar + 20 g sucrose + 200 mL apple juice + 600 mL Milli-Q water) for one hour (stage 1–2 embryos) or two hours plus two hours incubation at 25°C (stage 5 embryos). We dechorionated embryos for one minute in 100% bleach (Chlorox), washed with distilled water, and fixed for 20 minutes in 1:1 heptane:4% PFA in 1xPBS on a horizontal rotating shaker. We devitellinized by manual shaking in 1:1 heptane:methanol and washed with 100% methanol (note that for experiments with phalloidin staining, we used 80% ethanol in place of methanol). We gradually stepped embryos out of methanol (or ethanol) into 1xPBS and permeabilized/blocked embryos in 1xPBT + 1 mg/mL BSA (1xPBS + 0.2% Triton X-100) three times for 30 minutes. We incubated embryos in primary antibody overnight at 4°C. The following day, we washed three times for one hour in 1xPBT + 1 mg/mL BSA and blocked two times for one hour in 1xPBT + 1mg/mL BSA + 5% NGS. We incubated in secondary antibody solution overnight at 4°C. The following day, we washed five times for 15 minutes in 1xPBT, mounted samples in Vectashield, and stored slides at −20°C until imaging.

For fluorescence *in situ* hybridization (FISH) experiments, we obtained hybridization chain reaction (HCR) probes and amplifiers from Molecular Instruments (Los Angeles, CA, USA; see Table S13 for list). We dechorionated and devitellinized stage 1–2 embryos as described above. For FISH experiments (in wild-type *D. melanogaster* and *D. virilis* tissues), we used the HCR RNA-FISH protocol for whole-mount fruit fly embryos from Molecular Instruments. For experiments combining HCR FISH with HA antibody staining (in chimeric *oskar* tissues), we gradually stepped embryos out of methanol in 1xPBS and permeabilized twice for five minutes in 1xPBT (1xPBS + 0.2% Tween-20), once for one hour in 1xPBTT (1xPBT + 1% Triton X-100), and once for five minutes in 1xPBT. We blocked twice for one hour in 1xPBT + 5% NGS and incubated in primary antibody overnight at 4°C. The following day, we washed four times for 30 minutes in 1xPBT and incubated in 2° antibody in 1xPBT + 5% NGS for three hours at room temperature. We washed five times for five minutes in 1xPBT and five times for five minutes in 5xSSCT (5xSSC (saline-sodium citrate) + 0.1% Triton X-100). We post-fixed in 4% formaldehyde (Millipore Sigma) and washed four times for five minutes in 5xSSCT. We incubated at 37°C for two hours in probe hybridization buffer (Molecular Instruments) and then added 16 nM of probe to incubate overnight at 37°C. The following day, we washed four times for 30 minutes at 37°C in probe wash buffer (Molecular Instruments) and then once for five minutes in 5xSSCT at room temperature. We incubated in amplification buffer (Molecular Instruments) for one hour and then added snap-cooled hairpins to incubate overnight at room temperature. The following day, we washed two times for five minutes and two times for 30 minutes in 5xSSCT. We incubated for one hour in Hoechst 33342 (Sigma Aldrich, 10 mg/mL stock solution) and then washed five times for five minutes in SSCT. We mounted samples in Vectashield and stored slides at −20°C before imaging.

### Antibody generation and validation

We generated a polyclonal antibody, chicken anti-*D. virilis* Oskar, using AbClonal (Woburn, MA, USA) custom antibody services. AbClonal synthesized the gene sequence of a partial protein antigen target (*Drosophila virilis* Oskar, UniProt ID: Q24741, M161-R260) and cloned it into an expression vector. They purified this protein and injected it into two chickens for immunization. Total IgY was precipitated and purified from 10 egg yolks of immunized chickens.

We validated antibodies with Western Blots using adult Oregon R ovary protein lysates generated from 30 pairs of ovaries squished in RIPA (radioimmunoprecipitation assay) buffer with protein amount estimated using a Pierce Detergent Compatible Bradford Assay (ThermoFisher Scientific) (Fig. S18A–B). We ran an SDS-PAGE gel with lysates mixed and boiled with 5% beta-mercaptoethanol and 5xPierce Lane Marker Non-Reducing Sample Buffer (ThermoFisher Scientific). We also loaded Precision Plus Dual Color (Bio-Rad, chemiluminescent blots). We conducted semi-dry transfer using a Bio-Rad Trans-Blot Turbo Transfer System (Bio-Rad) and the Trans-Blot Turbo RTA PVDF Transfer Kit (Bio-Rad).

We washed membranes once for five minutes in 1xTBS-T (1xTBS (Tris-buffered saline) + 0.1% Tween-20) and blocked for one hour in 1xTBST + 5% milk (Carnation Instant Nonfat Dry Milk). We washed twice for one minute, once for 15 minutes, and twice for five minutes in 1xTBST. We incubated membranes overnight at 4°C in 1:500 primary antibody in 1xTBST + 5% BSA + 0.03% sodium azide. The following day, we washed three times for ten minutes in 1xTSBT, incubated in 1:5000 anti-chicken HRP secondary antibody (ThermoFisher Scientific) in 1xTBST + 5% milk at room temperature, and washed three times for ten minutes in TBST. For detection, we used the SuperSignal West Pico PLUS Chemiluminescent Substrate (ThermoFisher Scientific). We imaged using a CCD camera on a Sapphire Biomolecular Imager.

### Larval cuticle preparations

We collected embryos for three hours on apple juice plates and incubated plates for 18 hours at 25°C. We dechorionated embryos/larvae for one minute in 100% bleach, washed with distilled water, and devitellinized by shaking in 1:1 heptane:methanol for five minutes. We removed the heptane and shook in 100% methanol for three minutes. We then washed once in methanol for three minutes, twice in 1xPBT (1xPBS + 0.1% Triton X-100) for three minutes, and once in 1xPBT for five minutes on a horizontal rotating shaker. We mounted samples in 3:1 lactic acid (Fisher Scientific):Milli-Q water, and we incubated slides at 65°C overnight. We stored slides at room temperature until imaging.

### Image acquisition and analysis

For germ plasm molecule enrichment experiments, we used a Nikon CSU-W1 and Plan Apo λ 20x/0.75 air objective to capture image stacks (60 µm total, 2 µm step size). We used Fiji to generate sum intensity projections of all stacks and masks of the tissue of interest within each projection. We used a previously described ^26^ Python script to calculate the relative signal enrichment for each pixel (z-score) in a projection, interpolate the relative signal enrichment along the anteroposterior axis, integrate the values for the 15% posterior-most positions, and plot the integrated posterior enrichment values. To ensure that we only tested for enrichment of germ plasm at the posterior in samples where the chimeric Oskar was localized at the posterior, we filtered our dataset to include only samples where the mean HA signal enrichment for the 15% posterior-most positions was greater than the mean plus three times the standard deviation of the HA signal enrichment observed in the genetic negative control for the experiments in the background with endogenous *oskar*. For the *oskar* RNA null experiments, we could not have a genetic negative control since *oskar* RNA null animals do not create late-stage oocytes or embryos ^54^, so we filtered for mean HA enrichment greater than the mean plus three times the standard deviation of signal in tissues of the same genotype with no primary antibody. We statistically compared each chimeric genotype’s distribution of integrated posterior enrichment values to those of the controls by bootstrapping, estimating p-values from the distribution of mean differences between sample and control from 100,000 simulated samples.

To supplement this analysis, we also measured colocalization of HA and each germ plasm molecule of interest as previously described ^26^. We found the Pearson correlation for the signal intensity between HA and another germ plasm molecule for the pixels in the 15% posterior-most end of the embryo with HA intensity values 2 standard deviations over the mean. We statistically compared distributions of Pearson correlation coefficients by bootstrapping, estimating p-values from the distribution of mean differences between sample and control from 100,000 simulated samples.

For the chimeric genotype with the Long Osk domain from *D. melanogaster* and the other three domains from *D. virilis*, sufficient posterior HA enrichment was found in oocytes to pass our threshold for analysis of germ plasm enrichment, but no embryos had sufficient enrichment to be included in the analysis (Fig. 3). We note that despite these low HA expression levels, sufficient *nanos* mRNA must be recruited for successful abdominal patterning, as we observed larval progeny of mothers expressing this chimera (data not shown).

For germ cell counting experiments, we used a Nikon CSU-W1 and Plan Apo λ 20x/0.75 air objective to capture image stacks (60 µm total, 1 µm step size). We used phalloidin staining to track the progress of cellularization and precisely stage embryos at the end of embryonic stage 5. We counted germ cells using the Fiji cell counter ^107^.

For axial patterning experiments, we used an EC Plan-Neofluar 20x/0.50 M27 on a Zeiss AxioImager Z1 and dark-field microscopy to capture image stacks (15 optical sections with a 1.47 µm step size). We used Helicon Focus (version 8.2.2) to render representative images with extended depth of field. We scored each cuticle patterning phenotype as “wild-type” (showing wild-type anterior-posterior (AP) patterning), “strong AP defects” (completely bicaudal, complete mirror image duplications of the posterior), or “partial AP defects” (some duplication of posterior segments in the anterior but not completely bicaudal). We calculated the Jensen– Shannon Divergence (JSD) for the distribution of “WT”, “strong anterior defects”, and “partial anterior defects” phenotypes in each chimeric background compared to the distribution of these phenotypes in the negative control (endogenous *oskar* only) and positive control (all *D. melanogaster oskar* sequence). A score of 0 indicates the distributions are identical, while a score of 1 indicates they are completely different.

For evaluation of the radial distribution of HA signal intensity in embryos, we analyzed images using a custom Python script. We rotated image stacks with the posterior of the embryo at the right, and used Cellpose ^108^ to segment the area containing the embryo (cyto3 model, diameter set to 1000). We cropped the image stacks and then rescaled in the z-dimension to account for differences in pixel dimensions. We computed the mean intensity along the yz axis to generate a 2D image and applied Otsu’s threshold ^109^ within the masked area. We then eroded over 20 iterations and computed the mean intensity within the eroded region with each iteration. We normalized each intensity profile and then fitted a straight line to each normalized profile. We plotted the slopes of these fitted lines and compared the distributions for *D. virilis* and *D. melanogaster* with statistical analysis by bootstrapping, estimating p-values from the distribution of mean differences between each pair of genotypes from 100,000 simulated samples. See Fig. S13C for an overview of this image analysis workflow.

### GPSite Analysis

Protein sequences for *D. melanogaster* and *D. virilis* Oskar, along with those of the chimeras, were input into the GPSite web server (https://bio-web1.nscc-gz.cn/app/GPSite) ^84^, and the output data were visualized with plots generated in Python. To generate the aligned line scores shown in Fig. S14, the *D. melanogaster* and *D. virilis* Oskar protein sequences were aligned with EMBOSS Needle ^110^.

## Key Resources Table

|  | SOURCE | IDENTIFIER |
|---|---|---|
| Antibodies |  |  |
| Rabbit polyclonal anti-Vasa | Laboratory of Akira Nakamura, Kumamoto University, Japan | N/A |
| Mouse monoclonal anti-HA | AbClonal | Cat# AE008 |
| Chicken polyclonal anti-Vasa | Extavour Lab<sup>26</sup> | N/A |
| Chicken polyclonal anti-Oskar (virilis) | This paper | N/A |
| Mouse monoclonal anti-ATP5A (15H4C4) | Abcam | Cat# ab14748 |
| Rat monoclonal anti-HA (3F10) | Millipore Sigma | Cat# 11867423001 |
| Rabbit polyclonal anti-Oskar | Laboratory of Anne Ephrussi, European Molecular Biology Laboratory, Germany | N/A |
| Guinea pig polyclonal anti-Smaug | Laboratory of Howard Lipshitz, University of Toronto, Canada | N/A |
| Rabbit polyclonal anti-Par-1 | Laboratory of Daniel St. Johnston, University of Cambridge, UK | N/A |
| Rabbit polyclonal anti-Nanos | Laboratory of Akira Nakmura, Kumamoto University, Japan | N/A |
| Rabbit polyclonal anti-GFP, conjugated to Alexa Fluor 488 | ThermoFisher Scientific | Cat# A-21311 |
| Alexa Fluor 488 AffiniPure goat anti-rabbit IgG (H+L) secondary antibody | Jackson ImmunoResearch Laboratories | Cat# 111-545-144 |
| Alexa Fluor 647 goat anti-mouse IgG (H+L) highly cross-adsorbed secondary antibody | ThermoFisher Scientific | Cat# A-21236 |
| Alexa Fluor 568 goat anti-chicken IgY (H+L) secondary antibody | ThermoFisher Scientific | Cat# A-11041 |
| Alexa Fluor 488 goat anti-mouse IgG (H+L) highly cross-adsorbed secondary antibody | ThermoFisher Scientific | Cat# A-11029 |
| Alexa Fluor 647 goat anti-rat IgG (H+L) highly cross-adsorbed secondary antibody | ThermoFisher Scientific | Cat# A-21247 |
| Alexa Fluor 647 donkey anti-rabbit IgG (H+L) highly cross-adsorbed secondary antibody | ThermoFisher Scientific | Cat# A-31573 |
| Alexa Fluor 568 goat anti-mouse IgG (H+L) highly cross-adsorbed secondary antibody | ThermoFisher Scientific | Cat# A-11004 |
| Alexa Fluor 568 goat anti-rabbit IgG (H+L) cross-adsorbed secondary antibody | ThermoFisher Scientific | Cat# A-11011 |
| Alexa Fluor 647 goat anti-chicken IgY (H+L) secondary antibody | ThermoFisher Scientific | Cat# A-21449 |
| Alexa Fluor 568 goat anti-rat IgY (H+L) secondary antibody | ThermoFisher Scientific | Cat# A-11077 |
| Alexa Fluor 488 goat anti-rat IgY (H+L) secondary antibody | ThermoFisher Scientific | Cat# A-11006 |
| Cy5 donkey anti-guinea pig IgG (H+L) secondary antibody | ThermoFisher Scientific | Cat # A-17648 |
| HRP goat anti-chicken IgY (H+L) secondary antibody | ThermoFisher Scientific | Cat# A-16054 |
| Hoechst 33342 | Sigma-Aldrich | Cat# B2261-25MG |
| Alexa Fluor 488 phalloidin | Invitrogen | Cat# A-12379 |
| TRITC phalloidin | Invitrogen | Cat# R415 |
| *D. melanogaster oskar* HCR probe, amp B1 (v3.0) | Molecular Instruments | N/A |
| *D. melanogaster nanos* HCR probe, amp B5 (v3.0) | Molecular Instruments | N/A |
| *D. melanogaster germ cell-less* HCR probe, amp B3 (v3.0) | Molecular Instruments | N/A |
| *D. melanogaster polar granule component* HCR probe, amp B4 (v3.0) | Molecular Instruments | N/A |
| *D. melanogaster oskar* 3’UTR HCR probe, amp B5 (v3.0) | Molecular Instruments | N/A |
| *D. virilis oskar* HCR probe, amp B1 (v3.0) | Molecular Instruments | N/A |
| *D. virilis nanos* HCR probe, amp B5 (v3.0) | Molecular Instruments | N/A |
| *D. virilis germ cell-less* HCR probe, amp B3 (v3.0) | Molecular Instruments | N/A |
| *D. virilis polar granule component* HCR probe, amp B4 (v3.0) | Molecular Instruments | N/A |
| HCR amplifier B1 488 (v3.0) | Molecular Instruments | N/A |
| HCR amplifier B5 555 (v3.0) | Molecular Instruments | N/A |
| HCR amplifier B3 594 (v3.0) | Molecular Instruments | N/A |
| HCR amplifier B4 647 (v3.0) | Molecular Instruments | N/A |
| Chemicals, peptides, and recombinant proteins |  |  |
| Chlorox Germicidal Bleach | Chlorox | Cat# 67619-32 |
| Heptane | Sigma-Aldrich | Cat# 32287 |
| Methanol | Sigma-Aldrich | Cat# 646377 |
| Ethanol | Sigma-Aldrich | Cat# E7023 |
| Paraformaldehyde, powder, 95% | Sigma-Aldrich | Cat# 158127 |
| Lactic acid, 90% solution in water | Fisher Scientific | Cat# AC189872500 |
| 2-mercaptoethanol, Omnipur | Avantor | Cat# EM-6050 |
| Formaldehyde solution | Sigma-Aldrich | Cat# 252549 |
| 5x Pierce Lane Marker Non-Reducing Sample Buffer | Thermo Scientific | Cat# 39001 |
| Precision Plus Protein Dual Color Standards | Bio-Rad | Cat# 1610394 |
| SuperSignal West Pico PLUS Chemiluminescent Substrate | Thermo Scientific | Cat# 34580 |
| Sodium azide | Sigma-Aldrich | Cat# S2002 |
| *Drosophila virilis* Oskar, (UniProt ID: Q24741, M161-R260 aa) | This paper | N/A |
| Critical commercial assays |  |  |
| HiSpeed Plasmid Kit | Qiagen | Cat# 12643 |
| QIAprep Spin Miniprep Kit | Qiagen | Cat# 27104 |
| Pierce Detergent Compatible Bradford Assay Kit | Thermo Scientific | Cat# 23246 |
| Trans-Blot Turbo RTA Mini 0.2 µm PVDF Transfer Kit | Bio-Rad | Cat# 1704272 |
| Deposited data |  |  |
| Code for analyzing radial distribution of HA signal | This paper | https://github.com/erivard/dmdv_osk_chimera.git |
| Sequences of *D. melanogaster–D. virilis oskar* chimeras | This paper | https://github.com/erivard/dmdv_osk_chimera.git |
| Experimental models: Organisms/strains |  |  |
| *y[1] sc\* v[1]; P{y[+] v[+]=(Dm1234-HA)attP40/CyO* | This paper | N/A |
| *y[1] sc\* v[1]; P{y[+] v[+]=(Dv1234-HA)attP40/CyO* | This paper | N/A |
| *y[1] sc\* v[1]; P{y[+] v[+]=(Dm123Dv4-HA)attP40/CyO* | This paper | N/A |
| *y[1] sc\* v[1]; P{y[+] v[+]=(Dm12Dv34-HA)attP40/CyO* | This paper | N/A |
| *y[1] sc\* v[1]; P{y[+] v[+]=(Dm13Dv24-HA)attP40/CyO* | This paper | N/A |
| *y[1] sc\* v[1]; P{y[+] v[+]=(Dm23Dv14-HA)attP40/CyO* | This paper | N/A |
| *y[1] sc\* v[1]; P{y[+] v[+]=(Dm1Dv234-HA)attP40/CyO* | This paper | N/A |
| *y[1] sc\* v[1]; P{y[+] v[+]=(Dm3Dv124-HA)attP40/CyO* | This paper | N/A |
| *y[1] sc\* v[1]; P{y[+] v[+]=(Dm2Dv134-HA)attP40/CyO* | This paper | N/A |
| *y[1] sc\* v[1]; P{y[+] v[+]=(Dm4Dv123-HA)attP40/CyO* | This paper | N/A |
| *y[1] sc\* v[1]; P{y[+] v[+]=(Dm14Dv23-HA)attP40/CyO* | This paper | N/A |
| *y[1] sc\* v[1]; P{y[+] v[+]=(Dm34Dv12-HA)attP40/CyO* | This paper | N/A |
| *y[1] sc\* v[1]; P{y[+] v[+]=(Dm234Dv1-HA)attP40/CyO* | This paper | N/A |
| *y[1] sc\* v[1]; P{y[+] v[+]=(Dm24Dv13-HA)attP40/CyO* | This paper | N/A |
| *y[1] sc\* v[1]; P{y[+] v[+]=(Dm124Dv3-HA)attP40/CyO* | This paper | N/A |
| *y[1] sc\* v[1]; P{y[+] v[+]=(Dm134Dv2-HA)attP40/CyO* | This paper | N/A |
| *y[1] sc\* v[1]; P{y[+] v[+]=(Dm1234-HA)attP40/CyO; osk[0.EGFP]/TM3 Sb[1]* | This paper | N/A |
| *y[1] sc\* v[1]; P{y[+] v[+]=(Dv1234-HA)attP40/CyO; osk[0.EGFP]/TM3 Sb[1]* | This paper | N/A |
| *y[1] sc\* v[1]; P{y[+] v[+]=(Dm123Dv4-HA)attP40/CyO; osk[0.EGFP]/TM3 Sb[1]* | This paper | N/A |
| *y[1] sc\* v[1]; P{y[+] v[+]=(Dm12Dv34-HA)attP40/CyO; osk[0.EGFP]/TM3 Sb[1]* | This paper | N/A |
| *y[1] sc\* v[1]; P{y[+] v[+]=(Dm13Dv24-HA)attP40/CyO; osk[0.EGFP]/TM3 Sb[1]* | This paper | N/A |
| *y[1] sc\* v[1]; P{y[+] v[+]=(Dm23Dv14-HA)attP40/CyO; osk[0.EGFP]/TM3 Sb[1]* | This paper | N/A |
| *y[1] sc\* v[1]; P{y[+] v[+]=(Dm1Dv234-HA)attP40/CyO; osk[0.EGFP]/TM3 Sb[1]* | This paper | N/A |
| *y[1] sc\* v[1]; P{y[+] v[+]=(Dm3Dv124-HA)attP40/CyO; osk[0.EGFP]/TM3 Sb[1]* | This paper | N/A |
| *y[1] sc\* v[1]; P{y[+] v[+]=(Dm2Dv134-HA)attP40/CyO; osk[0.EGFP]/TM3 Sb[1]* | This paper | N/A |
| *y[1] sc\* v[1]; P{y[+] v[+]=(Dm4Dv123-HA)attP40/CyO; osk[0.EGFP]/TM3 Sb[1]* | This paper | N/A |
| *y[1] sc\* v[1]; P{y[+] v[+]=(Dm14Dv23-HA)attP40/CyO; osk[0.EGFP]/TM3 Sb[1]* | This paper | N/A |
| *y[1] sc\* v[1]; P{y[+] v[+]=(Dm34Dv12-HA)attP40/CyO; osk[0.EGFP]/TM3 Sb[1]* | This paper | N/A |
| *y[1] sc\* v[1]; P{y[+] v[+]=(Dm234Dv1-HA)attP40/CyO; osk[0.EGFP]/TM3 Sb[1]* | This paper | N/A |
| *y[1] sc\* v[1]; P{y[+] v[+]=(Dm24Dv13-HA)attP40/CyO; osk[0.EGFP]/TM3 Sb[1]* | This paper | N/A |
| *y[1] sc\* v[1]; P{y[+] v[+]=(Dm124Dv3-HA)attP40/CyO; osk[0.EGFP]/TM3 Sb[1]* | This paper | N/A |
| *y[1] sc\* v[1]; P{y[+] v[+]=(Dm134Dv2-HA)attP40/CyO; osk[0.EGFP]/TM3 Sb[1]* | This paper | N/A |
| *w[1118]; Kr[If-1]/CyO; TI{w[+mC]=TI}osk[0.EGFP]/TM3, Ser[1]* | Bloomington Drosophila Stock Center | BDSC 67407 |
| *vas[PH165]/CyO; Dr/TM3* | Laboratory of Paul Lasko, McGill University, Canada | N/A |
| *Df(2L)b87e25/CyO* | Bloomington Drosophila Stock Center | BDSC 3138 |
| *Sco/CyO; osk-GFP[KI]/TM6B* | Laboratory of Graydon Gonsalvez, Augusta University, USA |  |
| *D. melanogaster* (*Oregon R*) | Laboratory of Cassandra Extavour | N/A |
| *D. virilis* | Gift from Rainbow Transgenics | N/A |
| Oligonucleotides |  |  |
| Primers for cloning chimeras, see Table S14 | This paper | N/A |
| Recombinant DNA |  |  |
| pVAL22_Dm1234-HA | This paper | N/A |
| pVAL22_Dv1234-HA | This paper | N/A |
| pVAL22_Dm123Dv4-HA | This paper | N/A |
| pVAL22_Dm12Dv34-HA | This paper | N/A |
| pVAL22_Dm13Dv24-HA | This paper | N/A |
| pVAL22_Dm23Dv14-HA | This paper | N/A |
| pVAL22_Dm1Dv234-HA | This paper | N/A |
| pVAL22_Dm3Dv124-HA | This paper | N/A |
| pVAL22_Dm2Dv134-HA | This paper | N/A |
| pVAL22_Dm4Dv123-HA | This paper | N/A |
| pVAL22_Dm14Dv23-HA | This paper | N/A |
| pVAL22_Dm34Dv12-HA | This paper | N/A |
| pVAL22_Dm234Dv1-HA | This paper | N/A |
| pVAL22_Dm24Dv13-HA | This paper | N/A |
| pVAL22_Dm124Dv3-HA | This paper | N/A |
| pVAL22_Dm134Dv2-HA | This paper | N/A |
| pVALIUM22 | Drosophila Genomics Resource Center | Cat# 1468 |
| *D. melanogaster oskar* genomic locus | Laboratory of Anne Ephrussi | GenBank Acc. No. AE014297 |
| *D. virilis oskar* genomic locus | Laboratory of Paul Macdonald | GenBank Acc. No. L22556 |
| Software and algorithms |  |  |
| Fiji | Schindelin et al.<sup>107</sup> | https://imagej.net/software/fiji/ |
| Anaconda Python Distribution | Anaconda | https://www.anaconda.com/ |
| Helicon Focus (v. 8.2.2) | HeliconSoft Limited | https://www.heliconsoft.com/heliconsoft-products/helicon-focus/ |
| AliView | Larsson<sup>111</sup> | https://ormbunkar.se/aliview/#DOWNLOAD |
| PyMOL | Schrödinger | https://www.pymol.org/ |
| GPSite | Biomedical HPC+AI Platform | https://bio-web1.nscc-gz.cn/app/GPSite |
| Other |  |  |
| Nikon CSU-W1 SoRA | Nikon | N/A |
| Zeiss AxioImager | Zeiss | N/A |
| Trans-Blot Turbo Transfer System | Bio-Rad | Cat# 1704150 |
| Sapphire Biomolecular Imager | Azure | N/A |

## Data Availability

Further information and requests for resources and reagents should be directed to and will be fulfilled by the lead contact, Cassandra G. Extavour.

Antibodies and fly strains developed for this study are available upon request. Source Data are provided with this paper in the associated file **dmdv_chim_rawdata.xlsx**.

## Code Availability

Custom scripts used for data analysis are available at https://github.com/erivard/dmdv_osk_chimera.git (commit ID e92a908).

