## Supplementary Information for "Protein sequence evolution underlies interspecies incompatibility of a cell fate determinant"

**This document contains the following materials:**

- Supplemental Figures S1 through S18
- Supplemental Tables S1 through S14
- Supplemental References

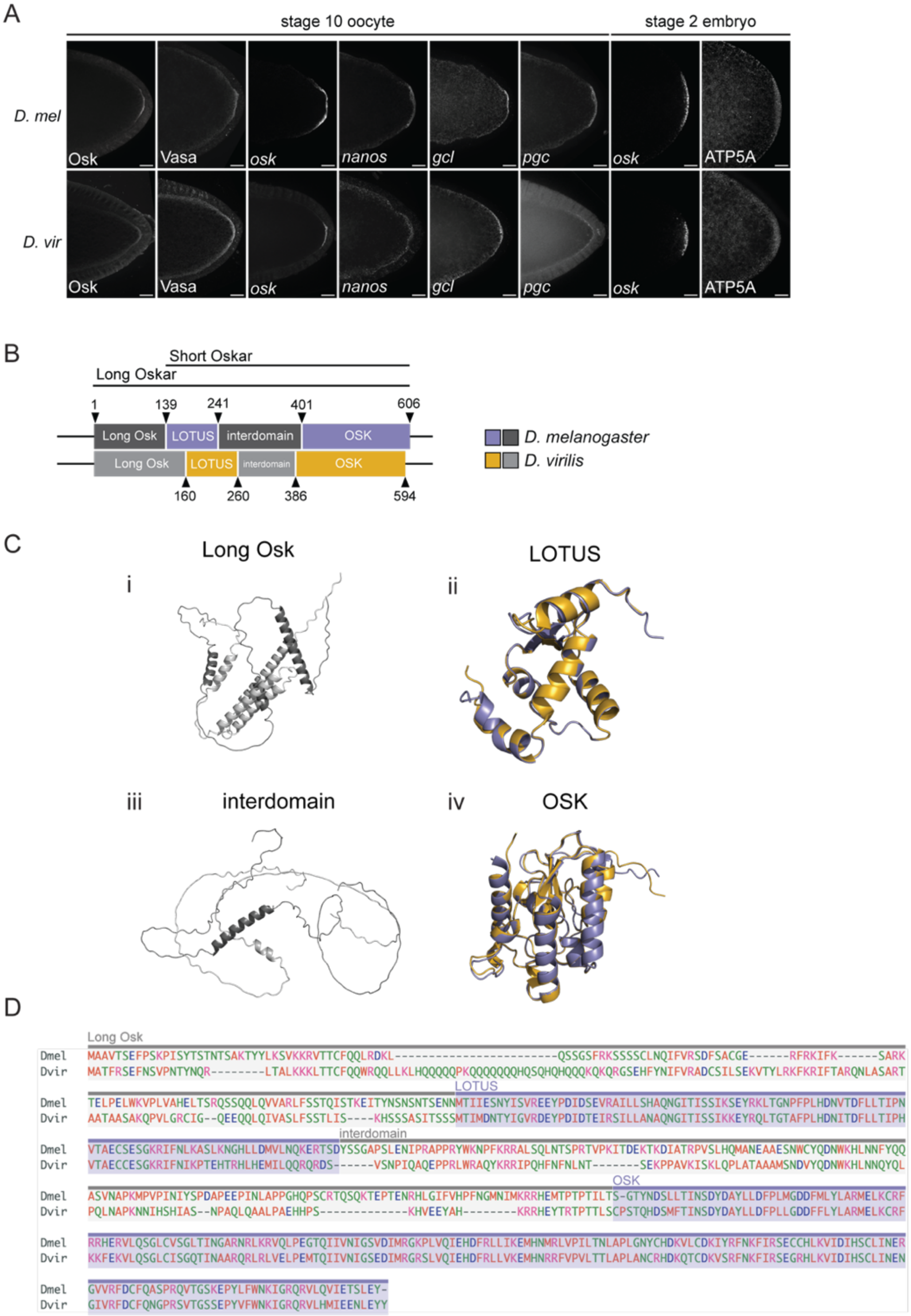


**Figure S1. Comparison of germ plasm, Oskar structure and sequence between *D. melanogaster* and *D. virilis*. A)** *D. melanogaster* (*D. mel*, top row) and *D. virilis* (*D. vir*, bottom row) stage 10 oocytes with Oskar, Vasa, *oskar*, *nanos*, *germ cell-less* (*gcl*), and *polar granule component* (*pgc*) labeled, and stage 2 embryos with *oskar* and mitochondria labeled (with anti-ATP5A antibody staining). All tissues are oriented with posterior to the right. Micrographs are maximum intensity projections of 20 optical sections (step size = 0.5 $\mu$m) and scale bars represent 20 $\mu$m. **B)** Schematic of *D. melanogaster* (top) and *D. virilis* (bottom) Oskar protein domain structures with amino acid residue number positions demarcating boundaries between domains. **C)** Individual *D. melanogaster* and *D. virilis* Oskar domain structural predictions generated with AlphaFold3 [1] and aligned in PyMOL (*D. melanogaster*: dark gray disordered regions, purple folded regions; *D. virilis*: light gray disordered regions, gold folded regions). **D)** Alignment of *D. melanogaster* and *D. virilis* Oskar protein sequences generated with MUSCLE. Domains shown in (**B**) are designated with colored background boxes.

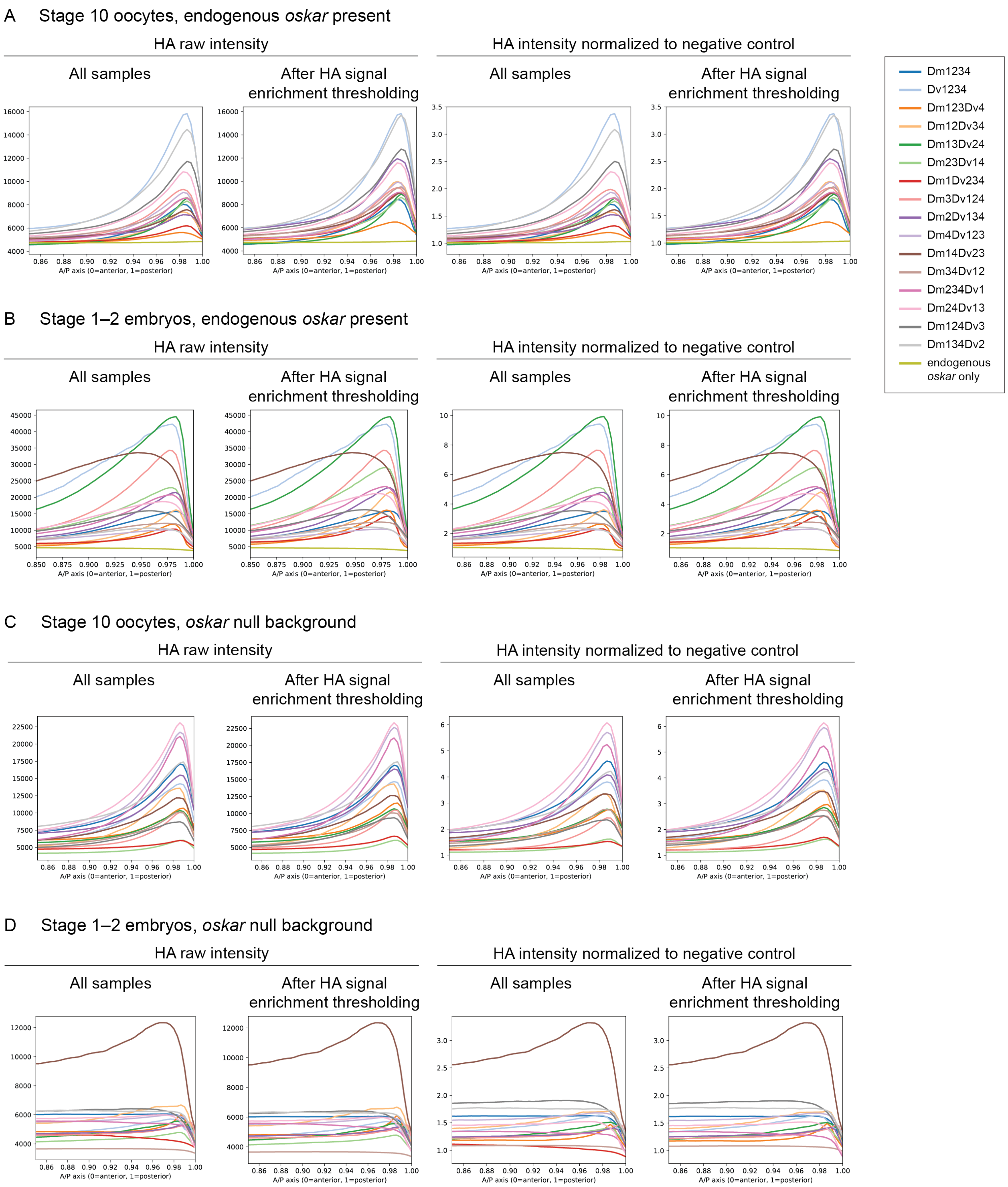


**Figure S2. Line scans of posterior HA signal. A–D)** Mean raw intensity (left) and intensity normalized to the average of negative control samples (right) are plotted for the posterior 15% of oocytes and embryos. Mean intensities are plotted for all samples that were included in the germ plasm enrichment analyses performed in this paper, based on thresholding for HA signal two standard deviations over the mean posterior signal for negative control samples. **A–B)** Negative control samples for normalization were samples expressing no transgene (endogenous *oskar* only). **C–D)** Negative control samples for normalization were embryos of the same genotype stained with no primary antibody. **A)** Stage 10 oocytes in a background with endogenous *oskar*. **B)** Stage 1–2 embryos in a background with endogenous *oskar***. C)** Stage 10 oocytes in an *oskar* null background. **D)** Stage 1–2 embryos in an *oskar* null background.

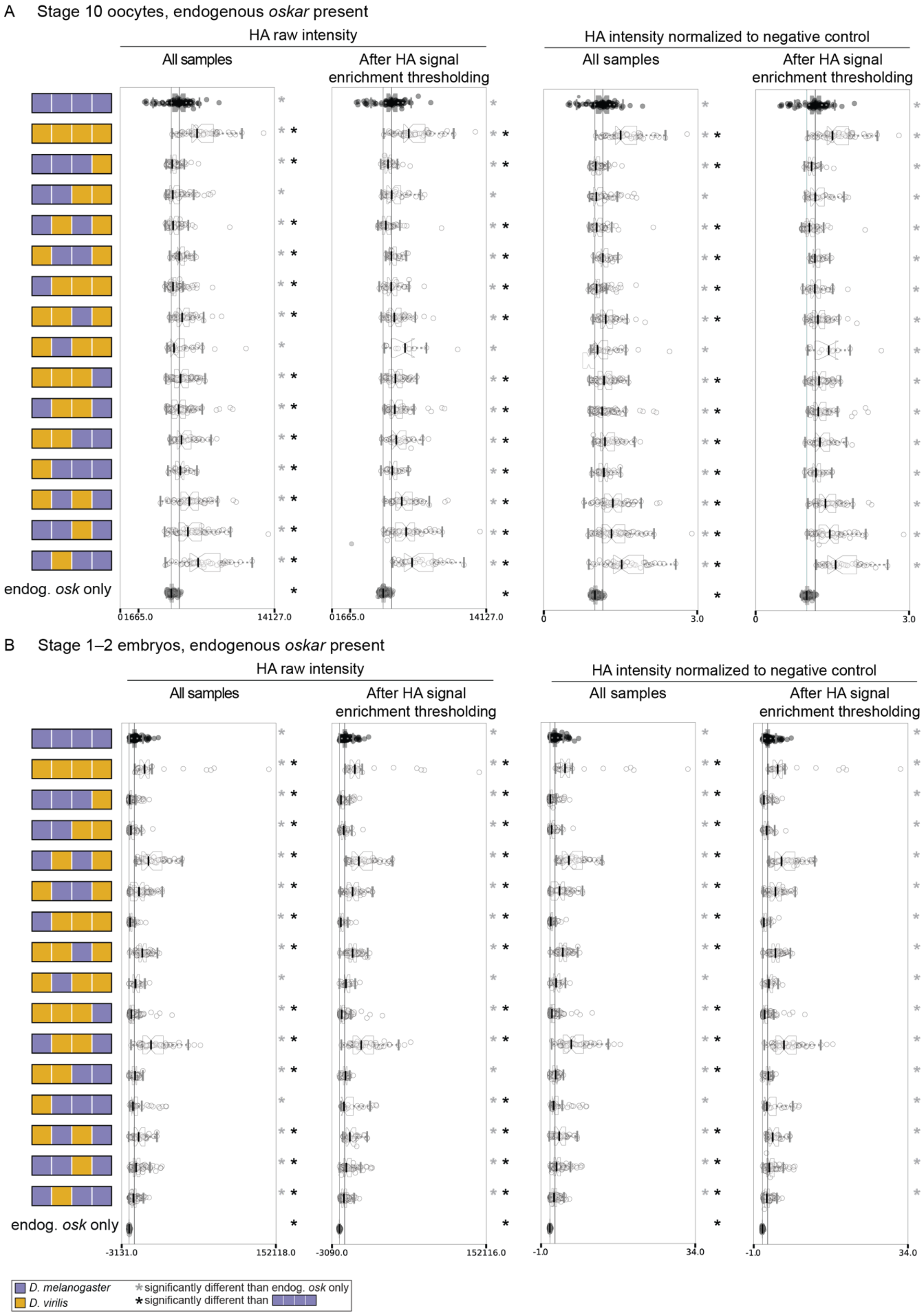
A Stage 10 oocytes, endogenous oskar present
HA raw intensity
All samples
After HA signal enrichment thresholding
HA intensity normalized to negative control
endog. osk only
0 1665.0
14127.0
0
3.0
B Stage 1–2 embryos, endogenous oskar present
-3131.0
152118.0
-3090.0
152116.0
-1.0
34.0
D. melanogaster
D. virilis
significantly different than endog. osk only
significantly different than

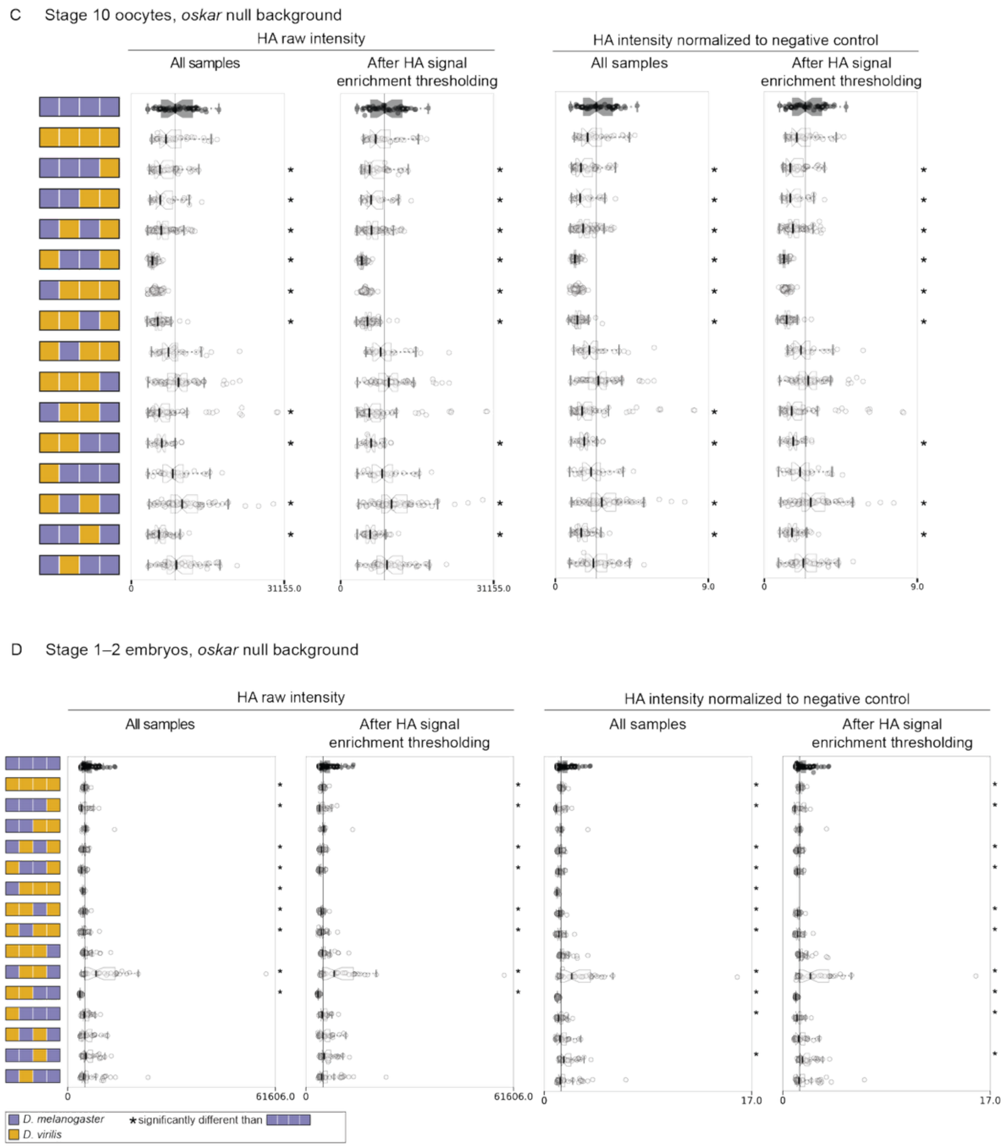


**Figure S3. Box plot of integrated posterior HA signal. A–D)** Integrated raw intensity (left) and intensity normalized to the average of negative control samples (right) are plotted for the posterior 15% of oocytes and embryos. Mean intensities are plotted for all samples that were included in the germ plasm enrichment analyses performed in this paper, included based on thresholding for HA signal two standard deviations over the mean posterior signal for negative control samples. **A–B)** Negative control samples for normalization were samples expressing no transgene (endogenous *oskar* only). **C–D)** Negative control samples for normalization were embryos of the same genotype stained with no primary antibody. **A)** Stage 10 oocytes in a background with endogenous *oskar*. **B)** Stage 1–2 embryos in a background with endogenous *oskar*. **C)** Stage 10 oocytes in an *oskar* null background. **D)** Stage 1–2 embryos in an *oskar* null background.

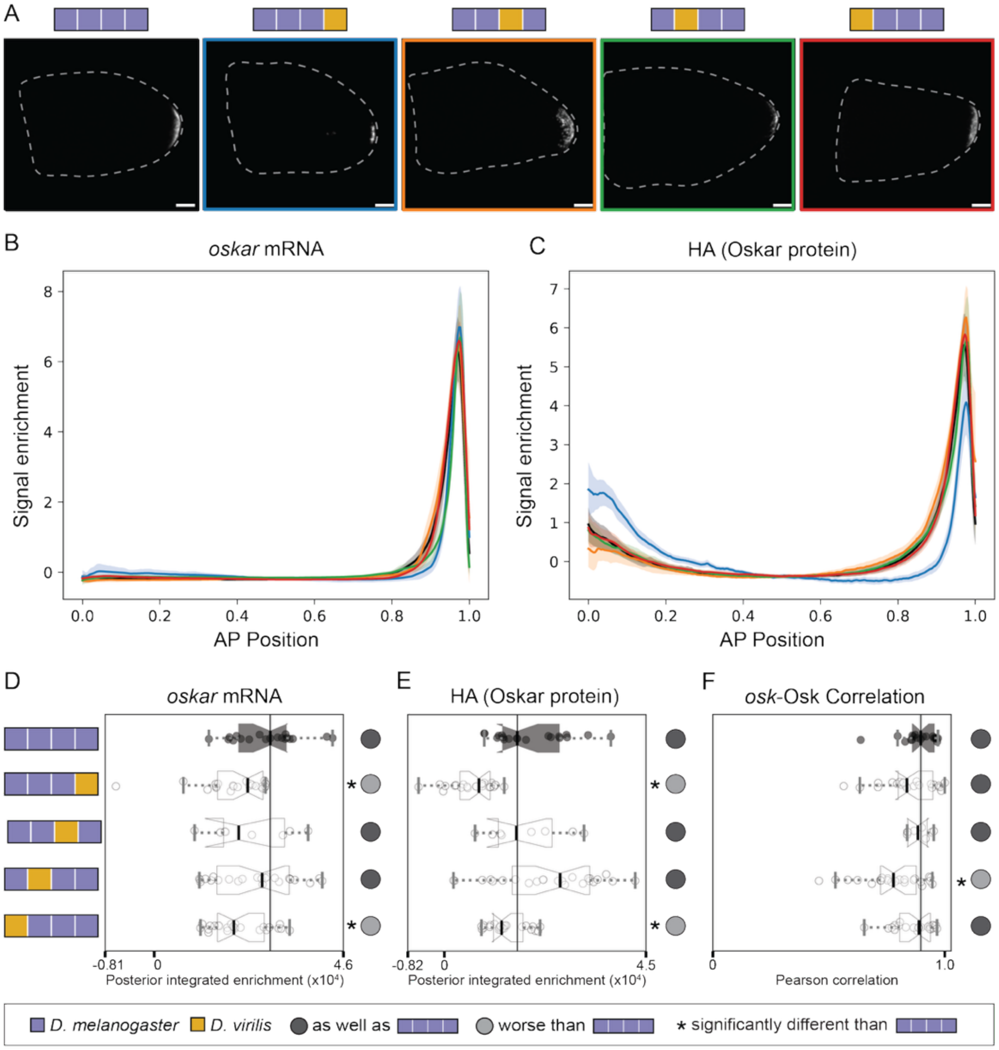


**Figure S4. Chimeric *oskar* mRNA successfully localizes to the posterior during oogenesis. A)** Representative images of oocytes from *D. melanogaster* mothers expressing chimeric Oskar sequences in an *oskar* RNA null background. HCR *in situ* hybridization marks the localization of the transgenic *oskar* mRNA. Images are oriented with posterior to the right. Micrographs are maximum intensity projections of 41 optical sections (step size = 1 $\mu$m) and scale bars = 20 $\mu$m. **B)** Line scans displaying *oskar* mRNA signal enrichment (z-score) across stage 10 oocytes. **C)** Line scans displaying HA (as a proxy for Oskar protein) signal enrichment (z-score) across stage 10 oocytes. For plots B) and C), on the X axis, 0 represents the anterior and 1 the posterior end of the oocyte. Mean (solid lines) and 95% confidence intervals (shaded regions) are shown. Line colors correspond to the chimeras shown in A). **D)** Posterior integrated enrichment of *oskar* mRNA in stage 10 oocytes from *D. melanogaster* mothers expressing chimeric Oskar proteins in an *oskar* RNA null background. **E)** Posterior integrated enrichment of HA (as a proxy for Oskar protein) in stage 10 oocytes from *D. melanogaster* mothers expressing chimeric Oskar proteins in an *oskar* RNA null background. **F)** Pearson correlation scores between intensities of the *oskar* mRNA and HA signals quantified in A) and B) respectively. For plots D-F), the distribution for eggs from flies expressing *D. melanogaster oskar* (positive control) is in black with the median marked with a black line. Significant difference from positive control distribution (p-value < 0.05, estimated from the distribution of difference of means between sample and control from 100,000 simulated samples) indicated by asterisk. Circles to the right of the plots indicate data interpretation as follows: dark gray = enriches the germ plasm molecule at least as well as *D. melanogaster oskar*; light gray = enriches the germ plasm molecule worse than *D. melanogaster oskar*. Sample sizes ≥8 for each condition; see Table S1.

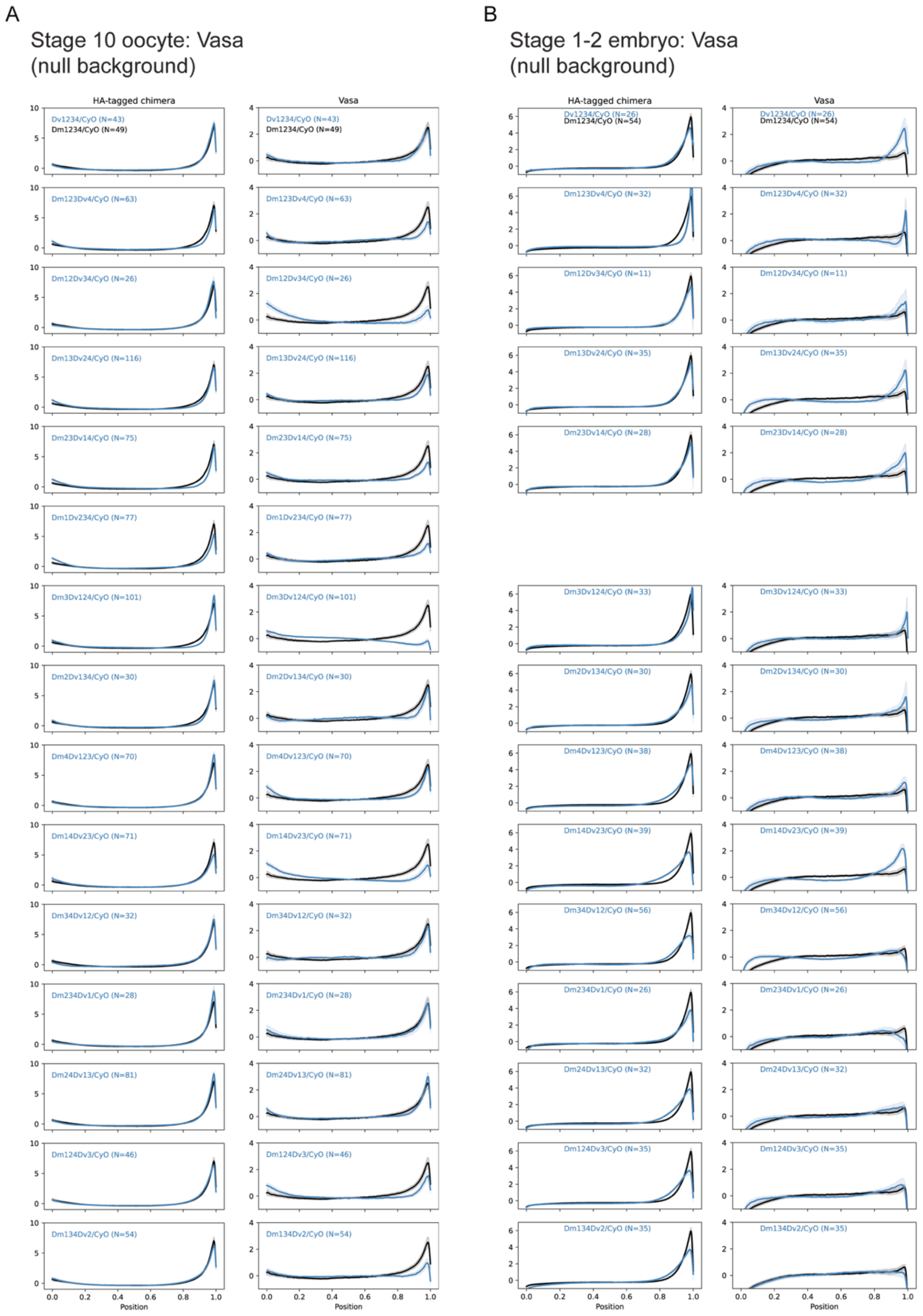

A
Stage 10 oocyte: Vasa
(null background)
HA-tagged chimera
Vasa
Dv1234/CyO (N=43)
Dm1234/CyO (N=49)
Dm123Dv4/CyO (N=63)
Dm12Dv34/CyO (N=26)
Dm13Dv24/CyO (N=116)
Dm23Dv14/CyO (N=75)
Dm1Dv234/CyO (N=77)
Dm3Dv124/CyO (N=101)
Dm2Dv134/CyO (N=30)
Dm4Dv123/CyO (N=70)
Dm14Dv23/CyO (N=71)
Dm34Dv12/CyO (N=32)
Dm234Dv1/CyO (N=28)
Dm24Dv13/CyO (N=81)
Dm124Dv3/CyO (N=46)
Dm134Dv2/CyO (N=54)
Position
B
Stage 1-2 embryo: Vasa
(null background)
HA-tagged chimera
Vasa
Dv1234/CyO (N=26)
Dm1234/CyO (N=54)
Dm123Dv4/CyO (N=32)
Dm12Dv34/CyO (N=11)
Dm13Dv24/CyO (N=35)
Dm23Dv14/CyO (N=28)
Dm3Dv124/CyO (N=33)
Dm2Dv134/CyO (N=30)
Dm4Dv123/CyO (N=38)
Dm14Dv23/CyO (N=39)
Dm34Dv12/CyO (N=56)
Dm234Dv1/CyO (N=26)
Dm24Dv13/CyO (N=32)
Dm124Dv3/CyO (N=35)
Dm134Dv2/CyO (N=35)
Position

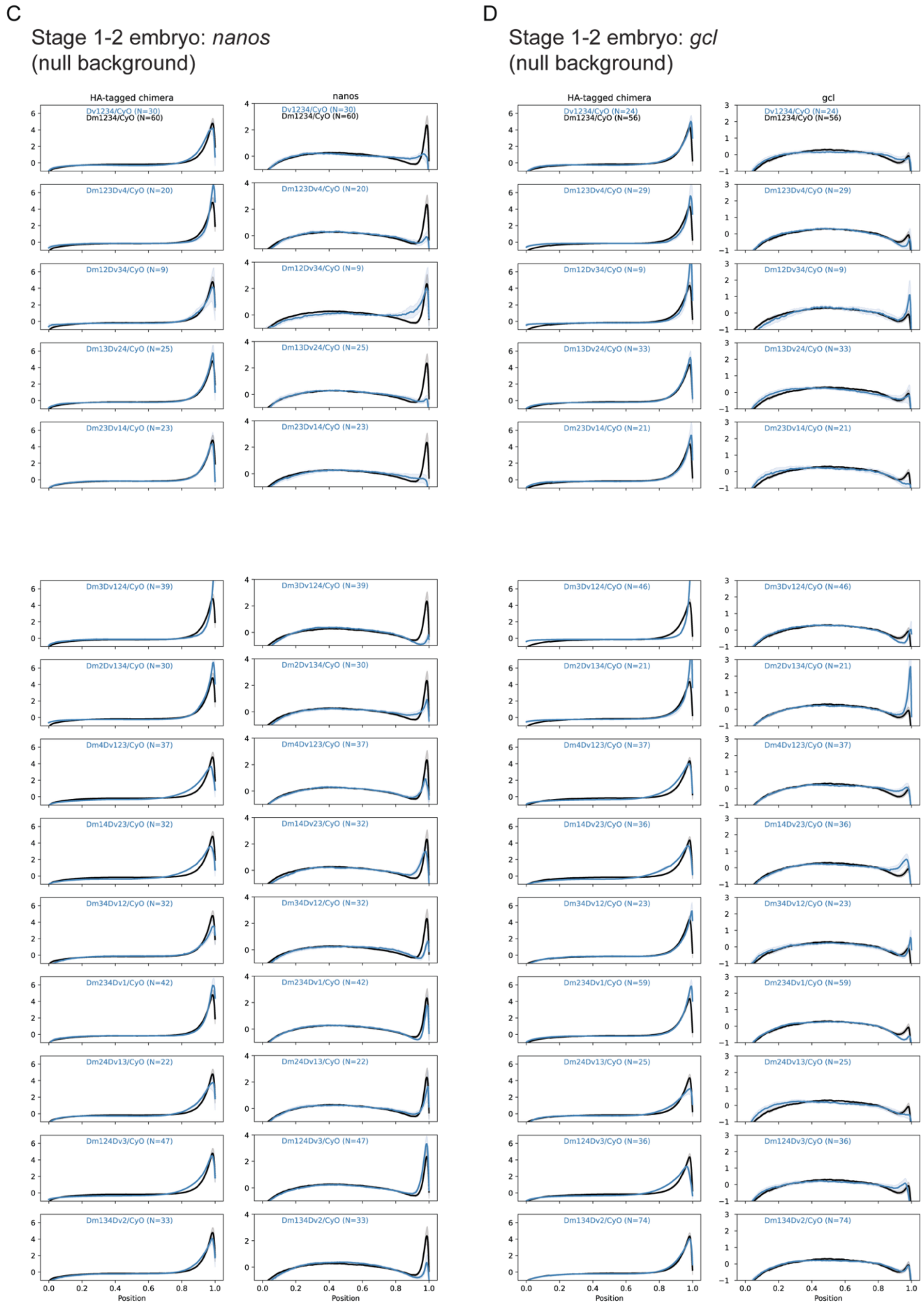


**Figure S5. A, B)** Vasa, **C)** *nanos,* and **D)** *gcl* signal enrichment (z-score) across stage 10 oocytes (A) and stage 1–2 embryos (B-D) in the *oskar* RNA null background *y[1] sc[*] v[1]; Kr[If-1]/CyO; TI{w[+mC]=TI}osk[0.EGFP]*. On the X axis, 0 designates the anterior and 1 the posterior. Mean (solid lines) and 95% confidence intervals (shaded regions) are shown. Black lines are measurements from positive control animals expressing the *D. melanogaster oskar* construct. Blue lines are measurements from each chimera genotype. Sample sizes for the measurements in each plot are noted in parentheses.

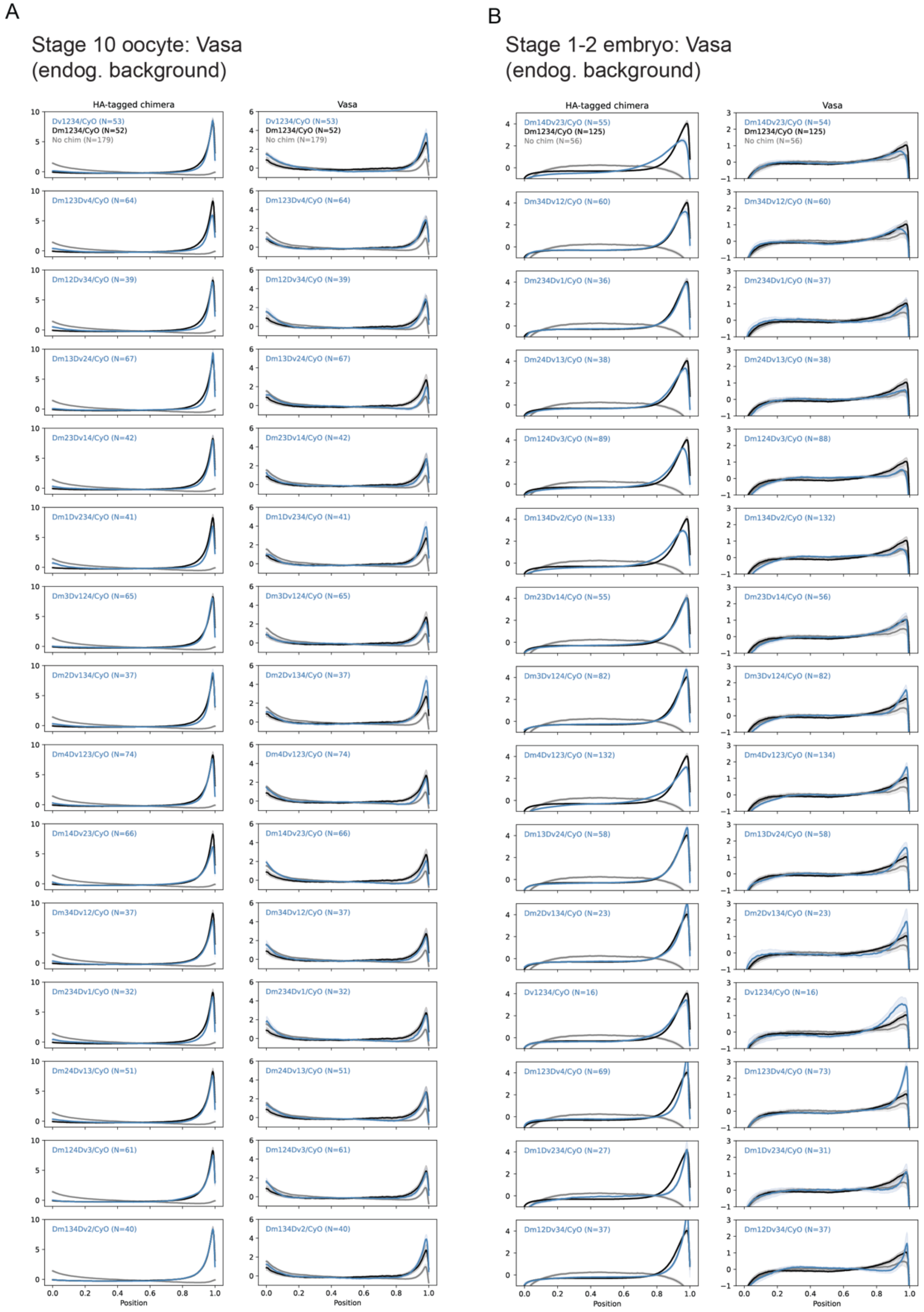
A
Stage 10 oocyte: Vasa
(endog. background)
HA-tagged chimera
Vasa
Dv1234/CyO (N=53)
Dm1234/CyO (N=52)
No chim (N=179)
Dm123Dv4/CyO (N=64)
Dm12Dv34/CyO (N=39)
Dm13Dv24/CyO (N=67)
Dm23Dv14/CyO (N=42)
Dm1Dv234/CyO (N=41)
Dm3Dv124/CyO (N=65)
Dm2Dv134/CyO (N=37)
Dm4Dv123/CyO (N=74)
Dm14Dv23/CyO (N=66)
Dm34Dv12/CyO (N=37)
Dm234Dv1/CyO (N=32)
Dm24Dv13/CyO (N=51)
Dm124Dv3/CyO (N=61)
Dm134Dv2/CyO (N=40)
Position
B
Stage 1-2 embryo: Vasa
(endog. background)
HA-tagged chimera
Vasa
Dm14Dv23/CyO (N=55)
Dm1234/CyO (N=125)
No chim (N=56)
Dm14Dv23/CyO (N=54)
Dm34Dv12/CyO (N=60)
Dm234Dv1/CyO (N=36)
Dm234Dv1/CyO (N=37)
Dm24Dv13/CyO (N=38)
Dm124Dv3/CyO (N=89)
Dm124Dv3/CyO (N=88)
Dm134Dv2/CyO (N=133)
Dm134Dv2/CyO (N=132)
Dm23Dv14/CyO (N=55)
Dm23Dv14/CyO (N=56)
Dm3Dv124/CyO (N=82)
Dm4Dv123/CyO (N=132)
Dm4Dv123/CyO (N=134)
Dm13Dv24/CyO (N=58)
Dm2Dv134/CyO (N=23)
Dv1234/CyO (N=16)
Dm123Dv4/CyO (N=69)
Dm123Dv4/CyO (N=73)
Dm1Dv234/CyO (N=27)
Dm1Dv234/CyO (N=31)
Dm12Dv34/CyO (N=37)
Position

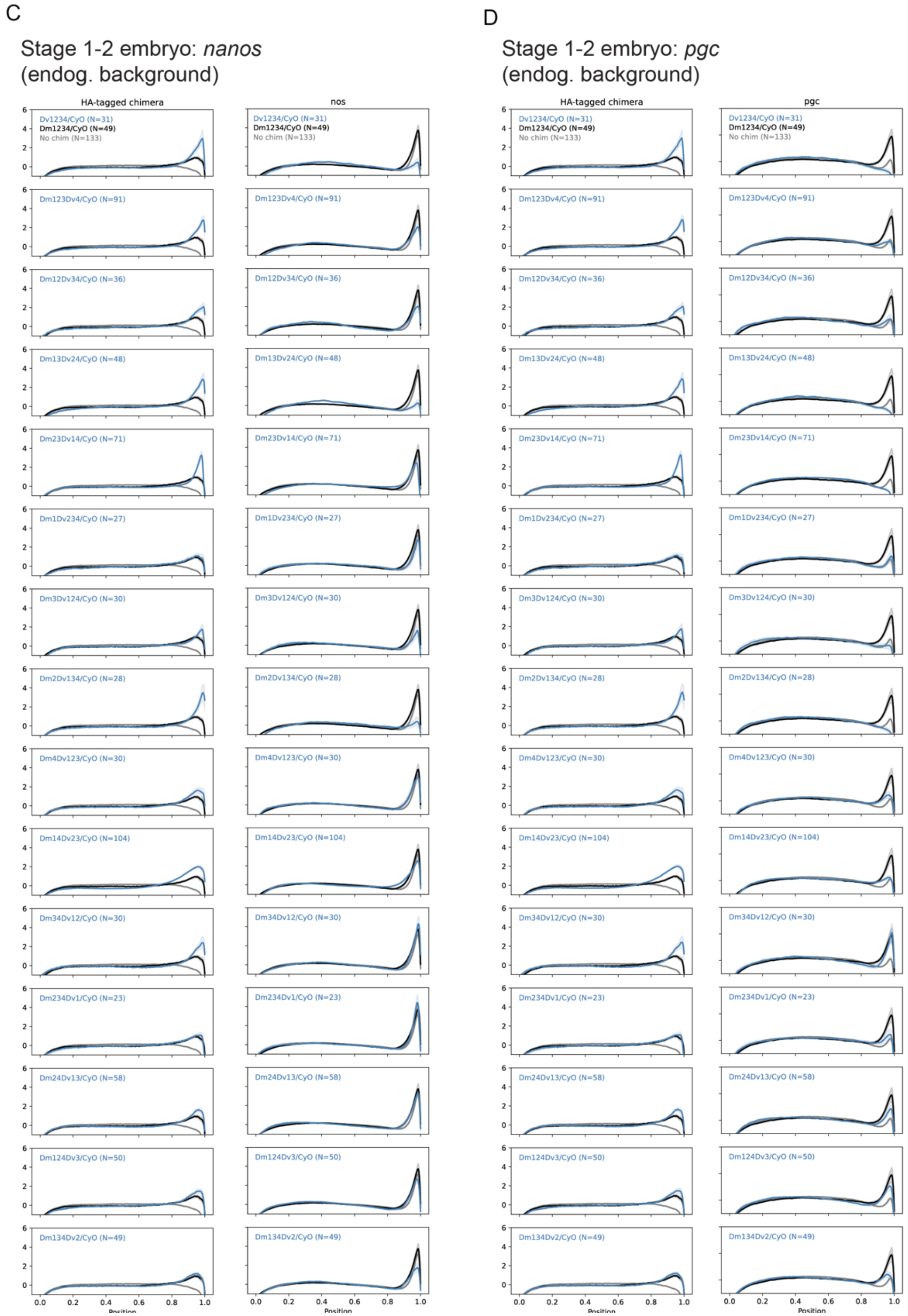


**Figure S6.** Vasa, *nanos,* and *pgc* signal enrichment (z-score) across stage 10 oocytes and stage 1–2 embryos for chimeras expressed in a background with endogenous *oskar*. On the X axis, 0 represents the anterior and 1 the posterior. Mean (solid lines) and 95% confidence intervals (shaded regions) are shown. Gray lines are measurements from flies not expressing a chimeric construct (only endogenous Oskar is expressed). Black lines are measurements from positive control animals expressing the *D. melanogaster oskar* construct. Blue lines are measurements from each chimera genotype. Sample sizes for the measurements in each plot are noted in parentheses.

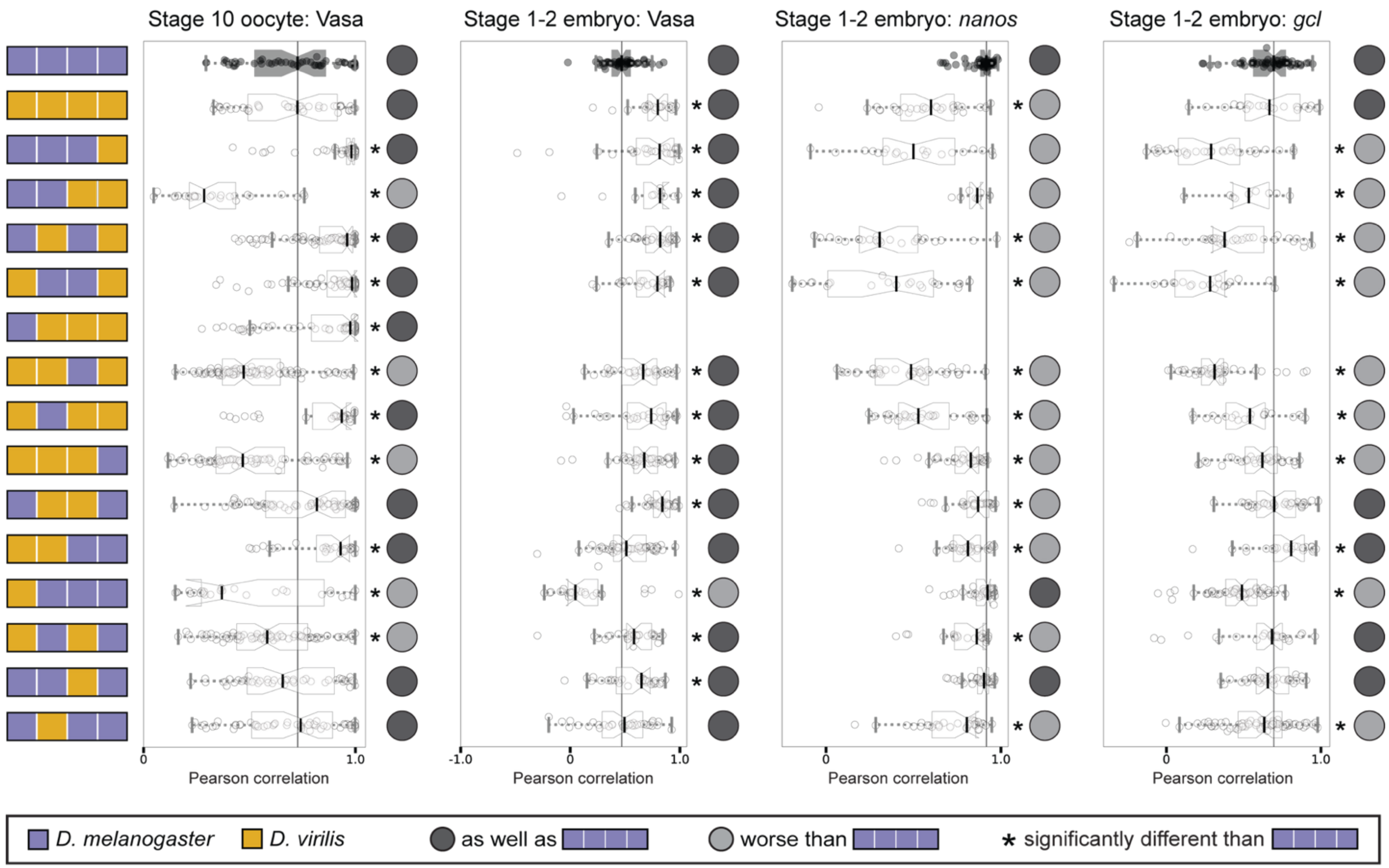


**Figure S7.** Pearson correlation scores between intensities of HA and Vasa, *nanos*, or *gcl* in stage 10 oocytes and stage 1–2 embryos from mothers expressing chimeric *oskar* in an *oskar* RNA null background (*y[1] sc[*] v[1]; Kr[If-1]/CyO; TI{w[+mC]=TI}osk[0.EGFP]).* For the chimeric genotype with the Long Osk domain from *D. melanogaster* and the other three domains from *D. virilis*, sufficient posterior HA enrichment was found in oocytes to pass our threshold for analysis of germ plasm enrichment, but no embryos had sufficient enrichment to be included in the analysis. Distribution for eggs laid by *D. melanogaster* mothers expressing the *D. melanogaster oskar* chimera (positive control) is in black with the median marked with a black line. Significant difference from positive control distribution (p-value < 0.05, estimated from the distribution of difference of means between sample and control from 100,000 simulated samples) indicated by asterisk. Circles to the right of the plots indicate data interpretation as follows: dark gray = chimera enriches the germ plasm molecule at least as well as *D. melanogaster oskar*; light gray = chimera enriches the germ plasm molecule worse than *D. melanogaster oskar*.

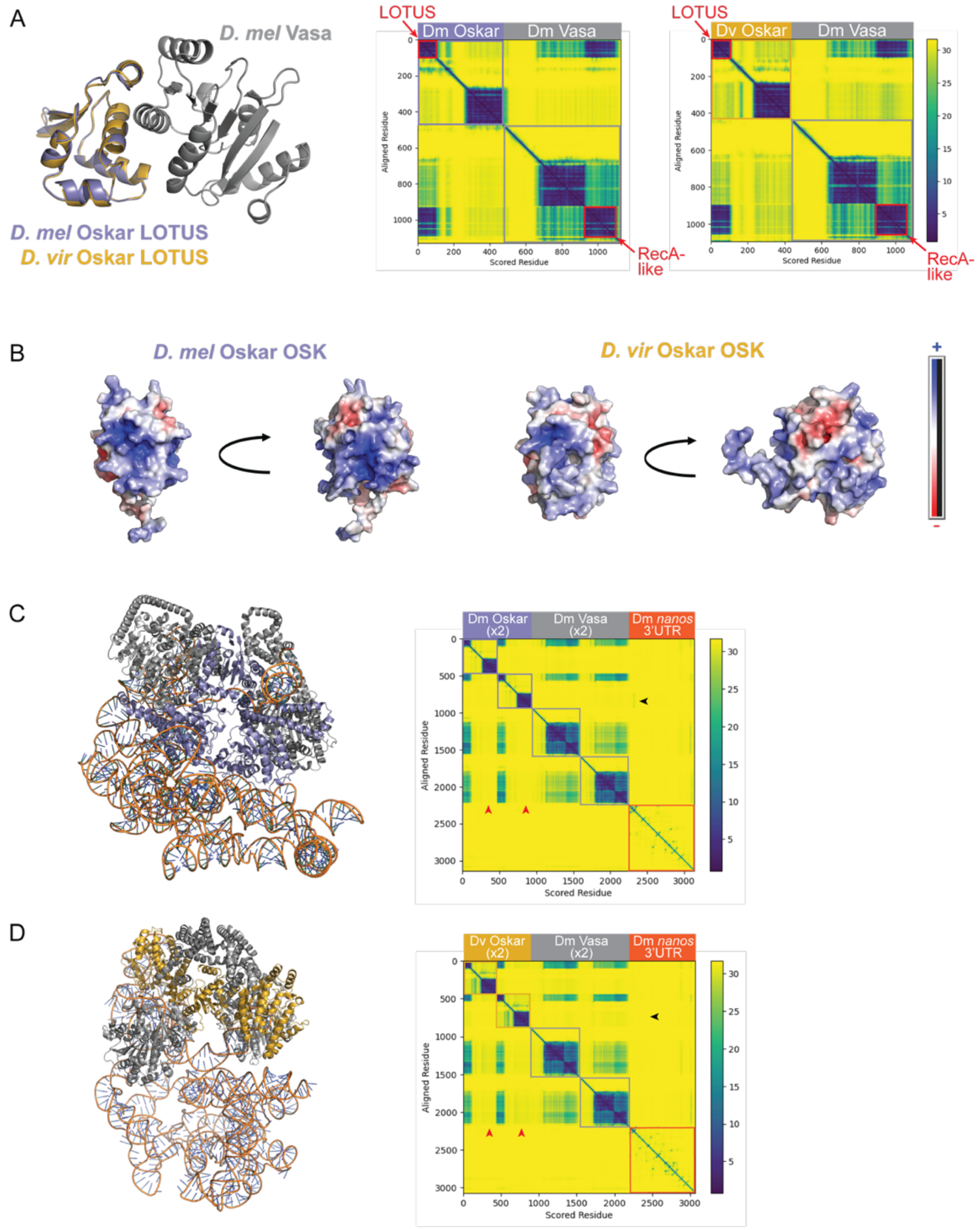


**Figure S8**. **A)** AlphaFold3 [1] predictions for *D. melanogaster* (purple) and *D. virilis* Short Oskar interactions with *D. melanogaster* Vasa (gray). The LOTUS domains of *D. melanogaster* and *D. virilis* Oskar, as well as *D. melanogaster* Vasa, are shown in structural predictions on the left. On the right, predicted aligned error plots reflect the confidence in the relative placement of residues in the predicted structure. Blue regions mark areas with low error and therefore higher confidence. Colored boxes designate the boundaries between the different proteins in each prediction. **B)** Electrostatic maps for AlphaFold3 predictions of the OSK domain from *D. melanogaster* (left) and *D. virilis* (right) Oskar protein. Blue indicates more positive and red indicates more negative surface charge. **C, D)** Structural predictions and accompanying predicted aligned error plots for two copies of *D. melanogaster* (C) and *D. virilis* (D) Short Oskar in complex with two copies of *D. melanogaster* Vasa and the *D. melanogaster nanos* 3'UTR. Red arrowheads mark where the *D. virilis* Oskar OSK domain has moderate predicted confidence of interaction with Vasa while *D. melanogaster* Oskar does not.

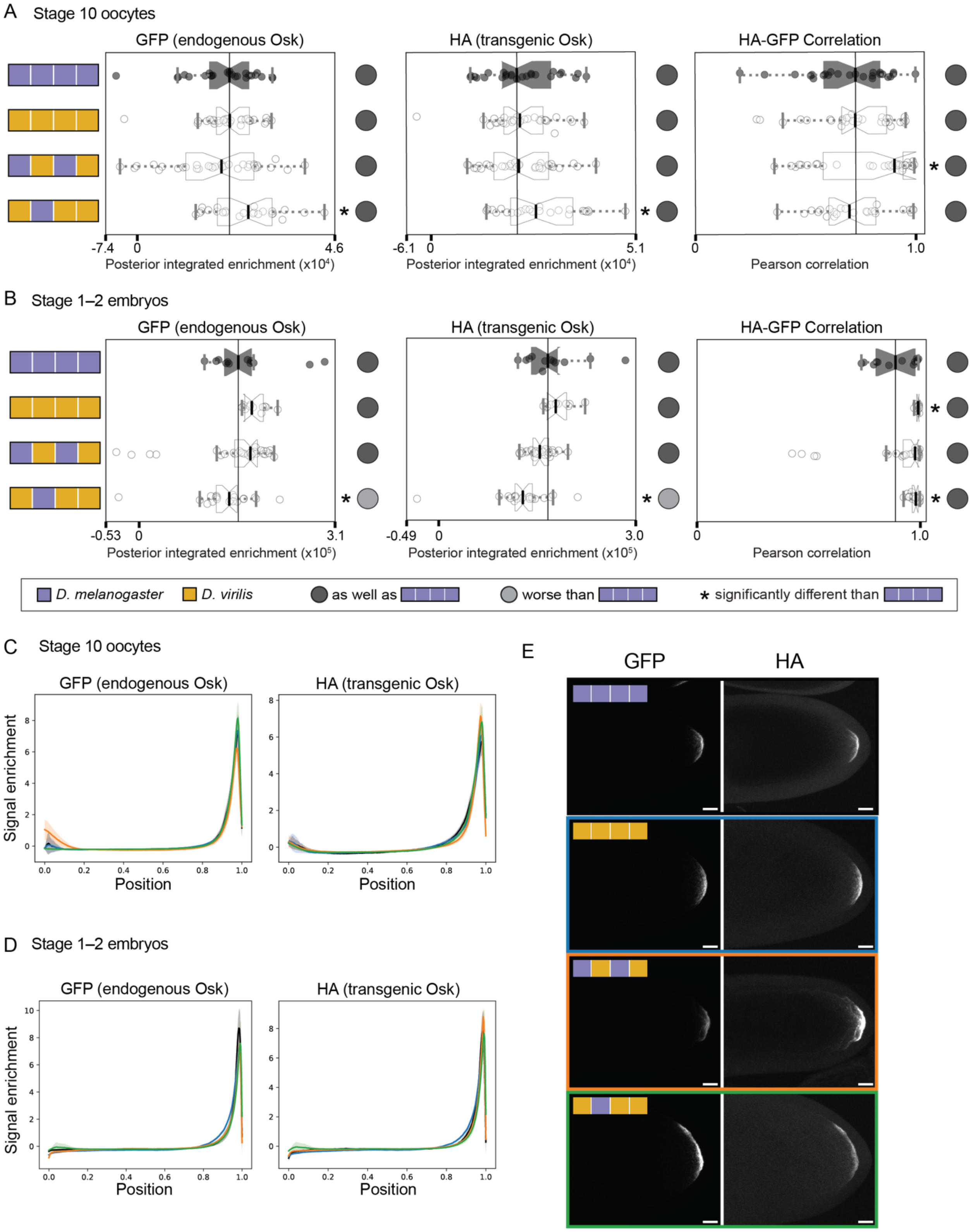

A Stage 10 oocytes
GFP (endogenous Osk)
HA (transgenic Osk)
HA-GFP Correlation
-7.4
0
4.6
Posterior integrated enrichment (x10⁴)
-6.1
0
5.1
Posterior integrated enrichment (x10⁴)
0
1.0
Pearson correlation
B Stage 1–2 embryos
GFP (endogenous Osk)
HA (transgenic Osk)
HA-GFP Correlation
-0.53
0
3.1
Posterior integrated enrichment (x10⁵)
-0.49
0
3.0
Posterior integrated enrichment (x10⁵)
0
1.0
Pearson correlation
D. melanogaster
D. virilis
as well as
worse than
* significantly different than
C Stage 10 oocytes
GFP (endogenous Osk)
HA (transgenic Osk)
Signal enrichment
Position
D Stage 1–2 embryos
GFP (endogenous Osk)
HA (transgenic Osk)
Signal enrichment
Position
E
GFP
HA

**Figure S9 (previous page). Chimeric Oskar co-localizes with endogenous Oskar. A–B)** Posterior integrated enrichment (left and middle plots) of GFP (endogenous Oskar) and HA (transgenic Oskar) in stage 10 oocytes **(A)** and stage 1–2 embryos **(B)** from *D. melanogaster* mothers expressing chimeric Oskar proteins in a background with endogenous Oskar labeled with GFP. Rightmost plot displays Pearson correlation scores between intensities of HA and GFP signal. For all plots, the distribution for eggs from flies expressing *D. melanogaster oskar* (positive control) is in black with the median marked with a black line. Significant difference from positive control distribution ($p$-value $< 0.05$, estimated from the distribution of difference of means between sample and control from 100,000 simulated samples) indicated by asterisk. Circles to the right of the plots indicate data interpretation as follows: dark gray = enriches the germ plasm molecule at least as well as *D. melanogaster oskar*; light gray = enriches the germ plasm molecule worse than *D. melanogaster oskar*. Sample sizes ≥12 for each condition; see Table S6. **C–D)** Line scans displaying GFP and HA signal enrichment (z-score) across stage 10 oocytes **(C)** and stage 1–2 embryos **(D)**. On the X axis, 0 represents the anterior and 1 the posterior. Mean (solid lines) and 95% confidence intervals (shaded regions) are shown. E) Representative images of oocytes from *D. melanogaster* mothers expressing chimeric Oskar sequences in a background with endogenous Oskar labeled with GFP. Anti-HA antibody staining marks the localization of transgenic Oskar and anti-GFP antibody staining marks the localization of endogenous Oskar protein. Images are oriented with posterior to the right. Micrographs are maximum intensity projections of 41 optical sections (step size = 1 $\mu$m) and scale bars = 20 $\mu$m.

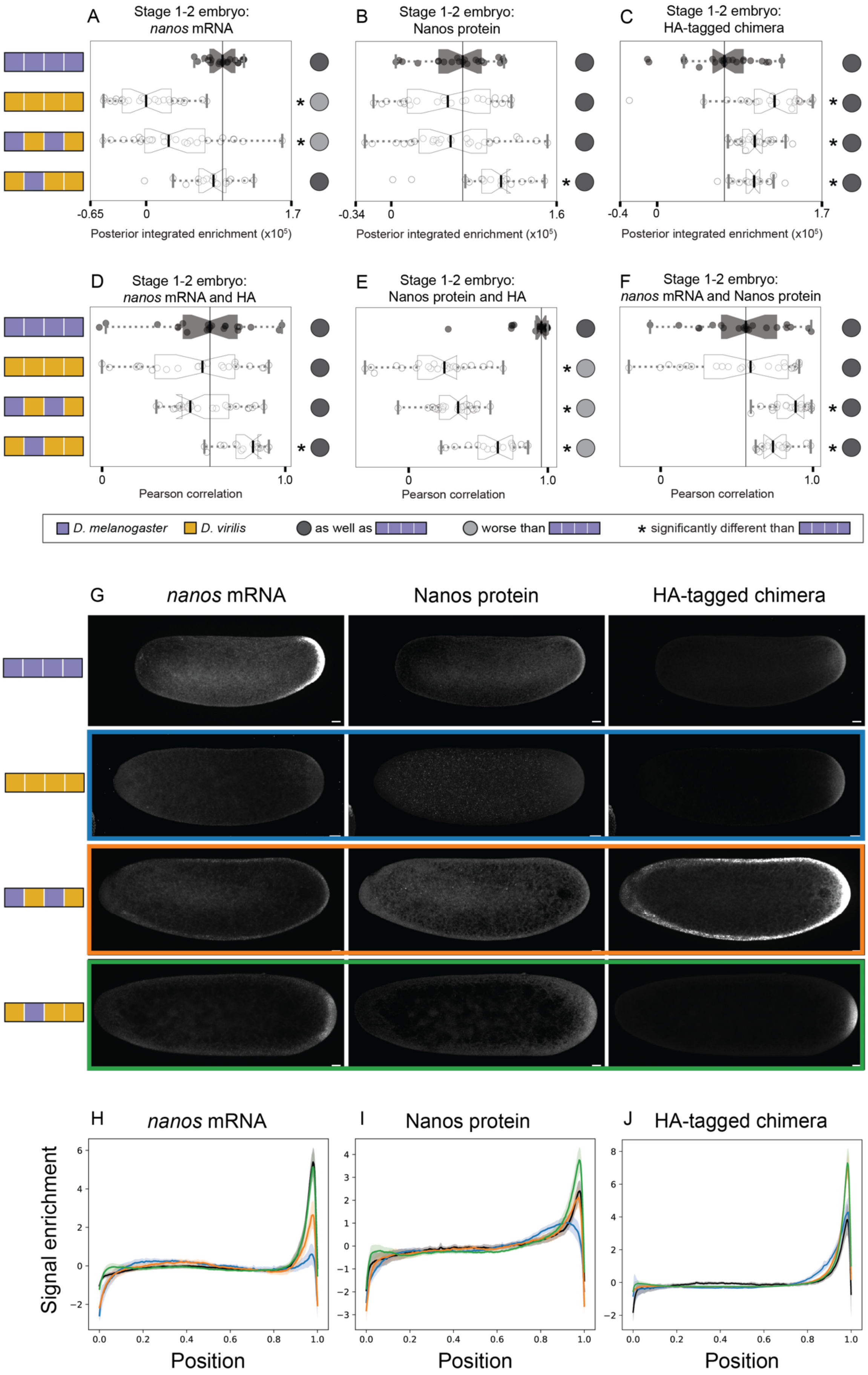

A
Stage 1-2 embryo: nanos mRNA
B
Stage 1-2 embryo: Nanos protein
C
Stage 1-2 embryo: HA-tagged chimera
Posterior integrated enrichment (x10^5)
D
Stage 1-2 embryo: nanos mRNA and HA
E
Stage 1-2 embryo: Nanos protein and HA
F
Stage 1-2 embryo: nanos mRNA and Nanos protein
Pearson correlation
D. melanogaster
D. virilis
as well as
worse than
* significantly different than
G
nanos mRNA
Nanos protein
HA-tagged chimera
H
nanos mRNA
I
Nanos protein
J
HA-tagged chimera
Signal enrichment
Position

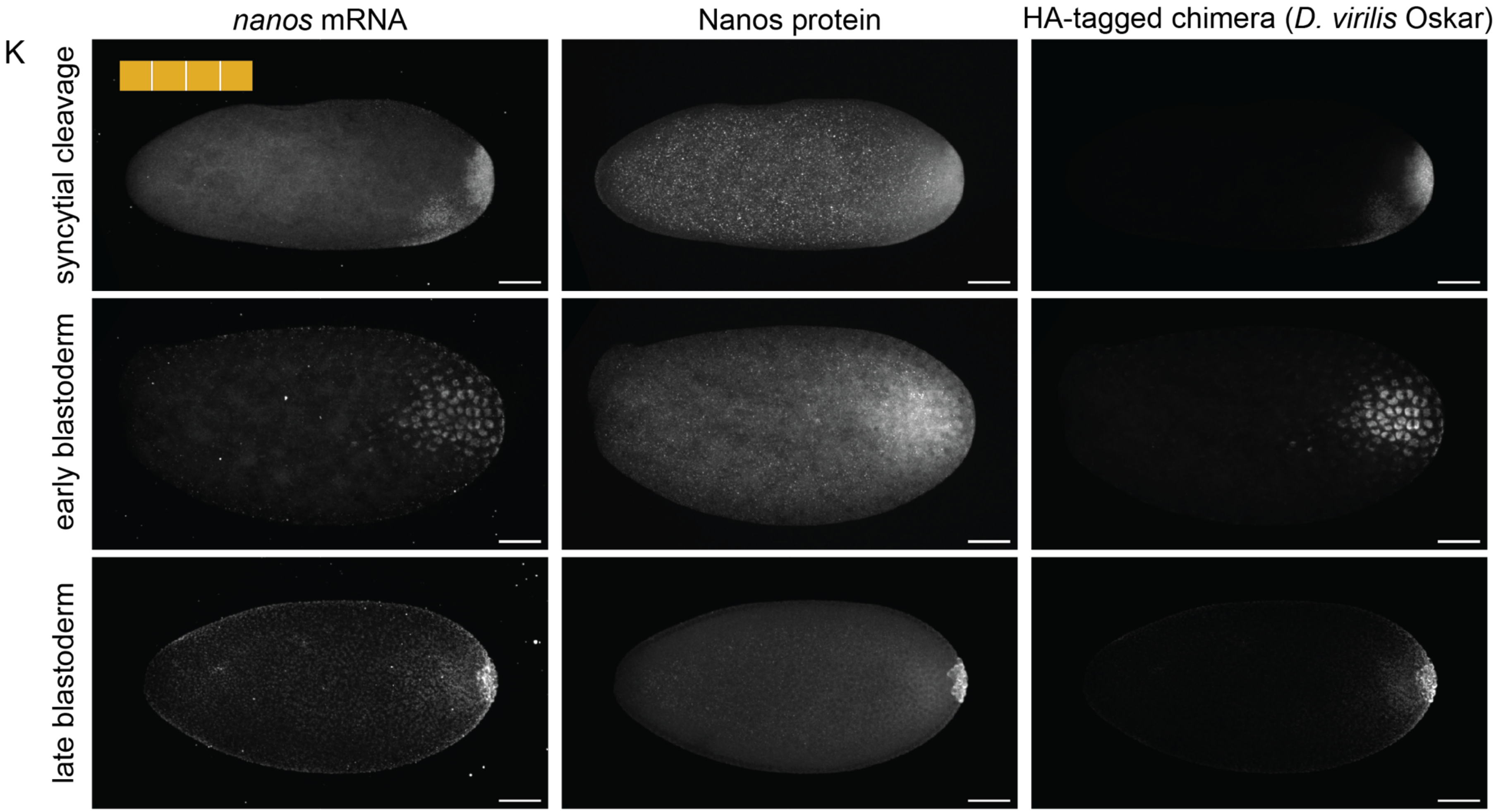
nanos mRNA
Nanos protein
HA-tagged chimera (D. virilis Oskar)
K
syncytial cleavage
early blastoderm
late blastoderm

**Figure S10 (previous pages). Quantification of the relationships between chimeric Oskar proteins, *nanos* mRNA and Nanos protein. A)** Posterior integrated enrichment of *nanos* mRNA. **B)** Posterior integrated enrichment of Nanos protein. **C)** Posterior integrated enrichment of HA-tagged chimeric Oskar protein. A-C) were quantified in stage 1–2 embryos from *D. melanogaster* mothers expressing chimeric Oskar proteins in a background with endogenous *oskar* present. **D)** Pearson correlation scores between intensities of HA and *nanos* mRNA signal. **E)** Pearson correlation scores between intensities of HA and Nanos protein signal. **F)** Pearson correlation scores between intensities of *nanos* mRNA and Nanos protein signal. All correlation scores are calculated for pixels in the 15% posterior-most end of the embryo with HA intensity values two standard deviations over the mean. For both integrated enrichment and correlation plots, the distribution for eggs from flies expressing D. *melanogaster oskar* (positive control) is in black with the median marked with a black line. Significant difference from positive control distribution (p-value < 0.05, estimated from the distribution of difference of means between sample and control from 100,000 simulated samples) indicated by asterisk. Circles to the right of the plots indicate data interpretation as follows: dark gray = enriches the germ plasm molecule at least as well as *D. melanogaster oskar*; light gray = enriches the germ plasm molecule worse than *D. melanogaster oskar*. Sample sizes ≥17 for each condition; see Table S7. **G)** Representative images of embryos from *D. melanogaster* mothers expressing chimeric Oskar sequences in a background with endogenous *oskar* present and stained with anti-Nanos antibody, *nanos* HCR *in situ* probes, and anti-HA to mark the chimeric Oskar proteins. Images are oriented with posterior to the right. Micrographs are maximum intensity projections of 41 optical sections (step size = 1 $\mu$m) and scale bars = 20 $\mu$m. **H)** Line scans displaying *nanos* mRNA, **I)** Nanos protein, and **J)** HA-tagged chimeric Oskar protein signal enrichment (Y axis; z-score) across stage 1–2 embryos. On the X axes, 0 represents the anterior and 1 the posterior. Mean (solid lines) and 95% confidence intervals (shaded regions) are shown. **K)** Representative images of embryos at distinct early stages from *D. melanogaster* mothers expressing *D. virilis* Oskar in a background with endogenous *oskar* present and stained with anti-Nanos antibody, *nanos* HCR *in situ* probes, and anti-HA to mark the *D. virilis* Oskar protein. Images are oriented with posterior to the right. Micrographs are maximum intensity projections of 30 optical sections (step size = 2 $\mu$m). Scale bars = 50 $\mu$m.

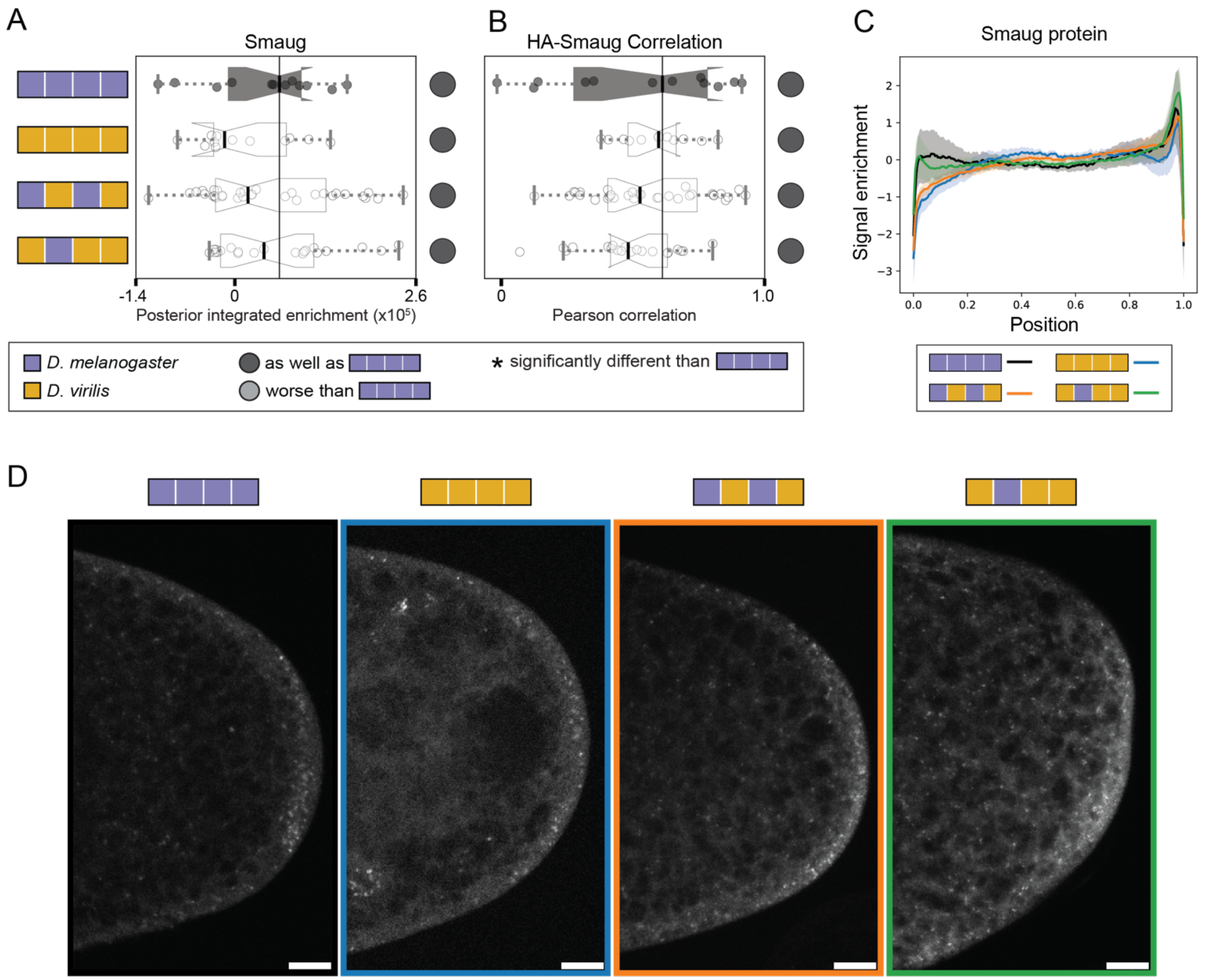


**Figure S11. Smaug remains localized to the germ plasm when expressing chimeric *oskar* proteins containing the *D. virilis* OSK domain in a background with endogenous *oskar*. A)** Posterior integrated enrichment of Smaug and HA in stage 1–2 embryos from *D. melanogaster* mothers expressing chimeric Oskar proteins in a background with endogenous *oskar* present. **B)** Pearson correlation scores between intensities of HA and Smaug signal. For both plots, the distribution for eggs from flies expressing D. *melanogaster oskar* (positive control) is in black with the median marked with a black line. Significant difference from positive control distribution ($p$-value < 0.05, estimated from the distribution of difference of means between sample and control from 100,000 simulated samples) indicated by asterisk. Circles to the right of the plots indicate data interpretation as follows: dark gray = enriches the germ plasm molecule at least as well as *D. melanogaster oskar*; light gray = enriches the germ plasm molecule worse than *D. melanogaster oskar*. Sample sizes ≥12 for each condition; see Table S8. **C)** Line scans displaying Smaug signal enrichment (z-score) across stage 2 embryos. On the X axis, 0 represents the anterior and 1 is the posterior. Mean (solid lines) and 95% confidence intervals (shaded regions) are shown. **D)** Representative images of embryos from *D. melanogaster* mothers expressing chimeric Oskar sequences in a background with endogenous *oskar* present and stained with anti-Smaug antibody (white). Images are oriented with posterior to the right. Micrographs are single optical sections and scale bars = 20 $\mu$m.

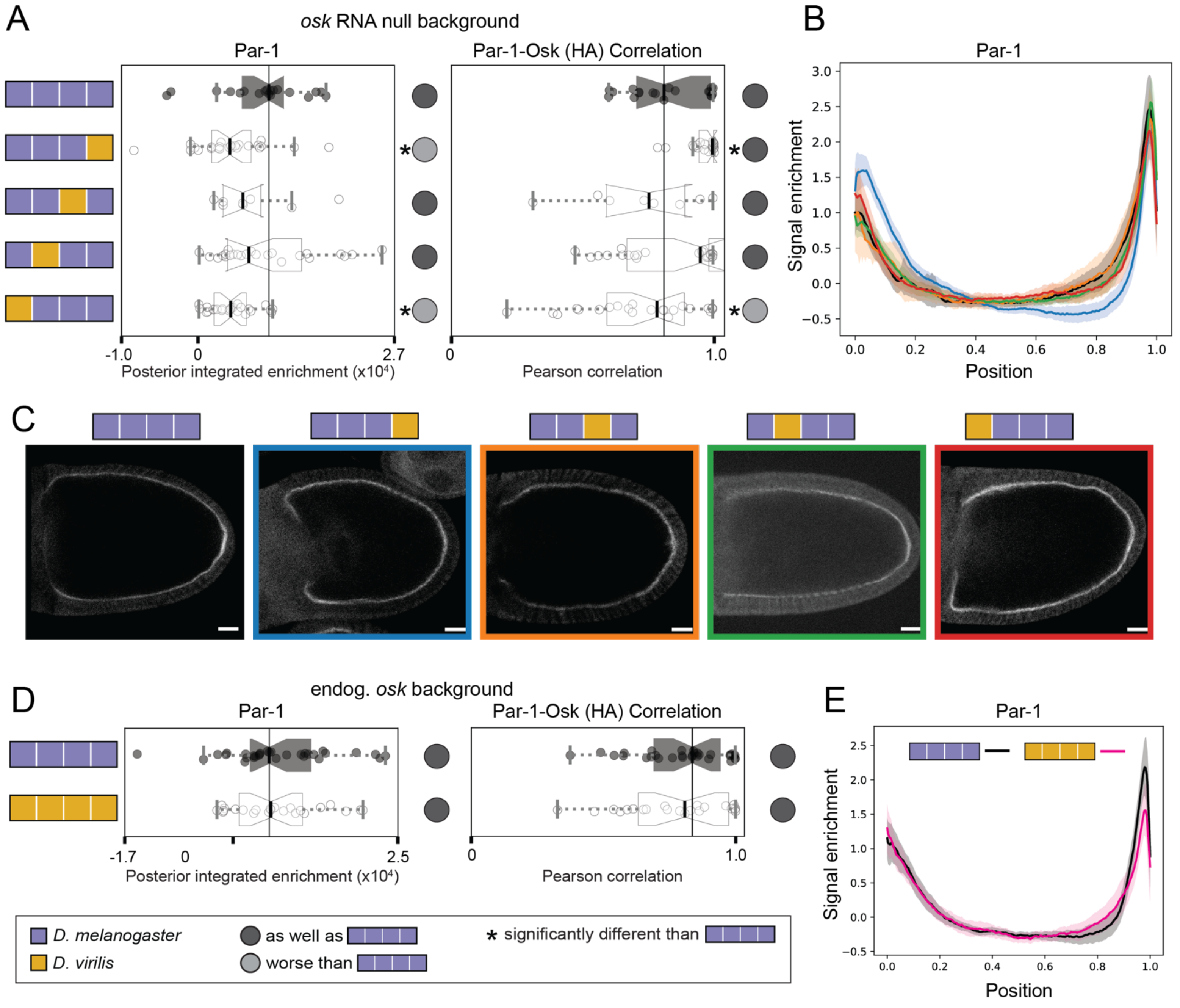

A
osk RNA null background
Par-1
Par-1-Osk (HA) Correlation
-1.0
0
2.7
Posterior integrated enrichment (x10⁴)
0
1.0
Pearson correlation
B
Par-1
Signal enrichment
Position
C
D
endog. osk background
Par-1
Par-1-Osk (HA) Correlation
-1.7
0
2.5
Posterior integrated enrichment (x10⁴)
0
1.0
Pearson correlation
D. melanogaster
D. virilis
as well as
worse than
* significantly different than
E
Par-1
Signal enrichment
Position

**Figure S12 (previous page). Par-1 exhibits some variation in posterior enrichment in stage 10 oocytes for chimeras expressed in an *oskar* RNA null background. A)** Posterior integrated enrichment (left) of Par-1 protein in stage 10 oocytes from *D. melanogaster* mothers expressing chimeric Oskar proteins in an *oskar* RNA null background. The plot on the right displays Pearson correlation scores between intensities of HA and Par-1 signal. For both plots, the distribution for eggs from flies expressing D. *melanogaster oskar* (positive control) is in black with the median marked with a black line. Significant difference from positive control distribution (p-value < 0.05, estimated from the distribution of difference of means between sample and control from 100,000 simulated samples) indicated by asterisk. Circles to the right of the plots indicate data interpretation as follows: dark gray = enriches the germ plasm molecule at least as well as *D. melanogaster oskar*; light gray = enriches the germ plasm molecule worse than *D. melanogaster oskar*. Sample sizes ≥8 for each condition; see Table S9. **B)** Line scans displaying Par-1 signal enrichment (z-score) across *oskar* RNA null stage 10 oocytes. On the X axis, 0 represents the anterior and 1 the posterior. Mean (solid lines) and 95% confidence intervals (shaded regions) are shown. **C)** Representative images of oocytes from *D. melanogaster* mothers expressing chimeric Oskar sequences in an *oskar* RNA null background. Anti-Par1 antibody staining (white) marks its localization. Images are oriented with posterior to the right. Micrographs depict single optical sections and scale bars = 20 $\mu$m. **D)** Posterior integrated enrichment (left) of Par-1 protein in stage 10 oocytes from *D. melanogaster* mothers expressing chimeric Oskar proteins in a background with endogenous *oskar* present. The plot on the right displays Pearson correlation scores between intensities of HA and Par-1 signal. For both plots, the distribution for eggs from flies expressing D. *melanogaster oskar* (positive control) is in black with the median marked with a black line. Significant difference from positive control distribution (p-value < 0.05, estimated from the distribution of difference of means between sample and control from 100,000 simulated samples) indicated by asterisk. Circles to the right of the plots designate the same interpretations described above. Sample sizes ≥23 for each condition; see Table S10. **E)** Line scans displaying Par-1 signal enrichment (z-score) across stage 10 oocytes in a background with endogenous *oskar* present. On the X axis, 0 represents the anterior and 1 the posterior. Mean (solid lines) and 95% confidence intervals (shaded regions) are shown.

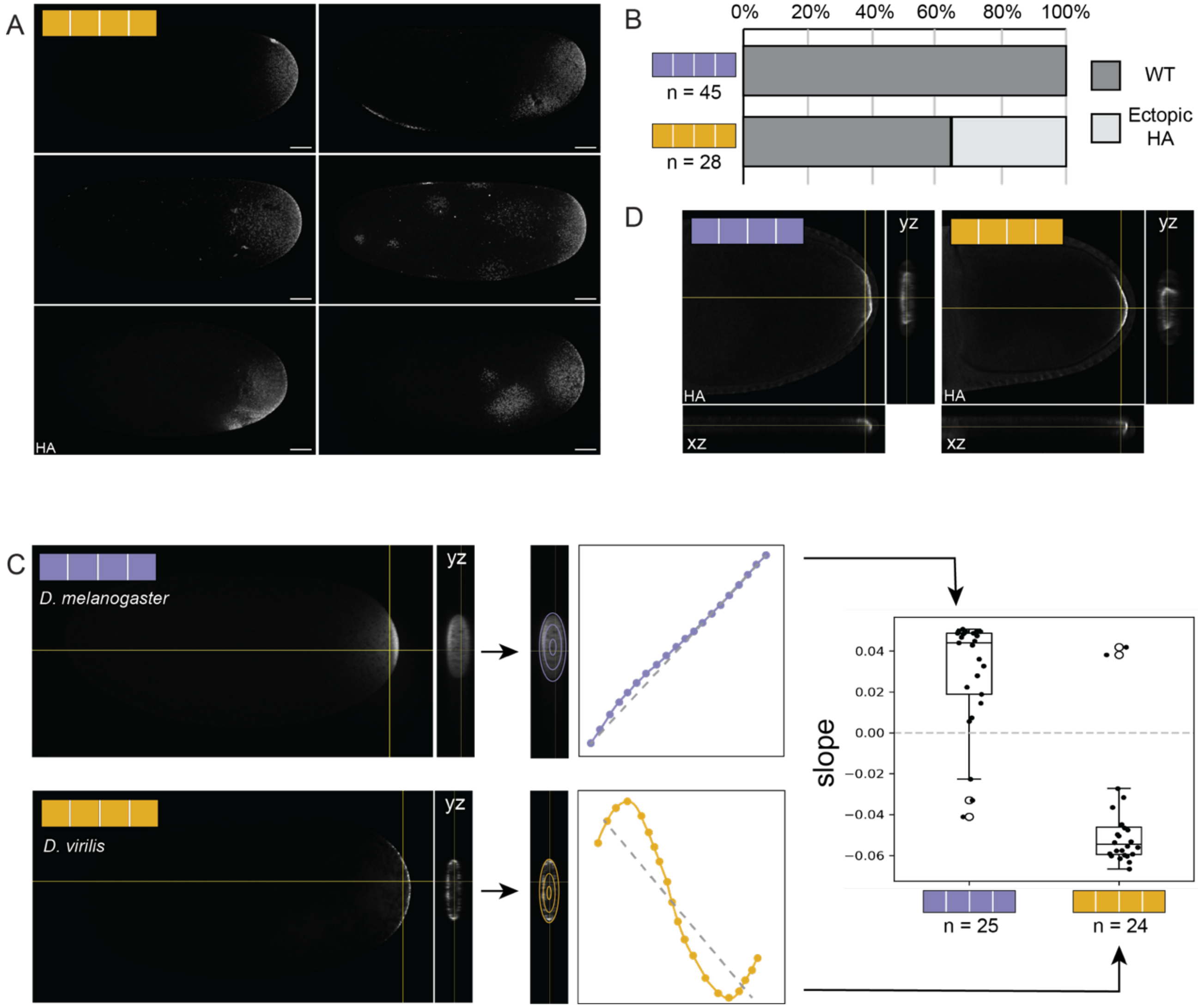


**Figure S13. A)** Cleavage stage embryos from *D. melanogaster* mothers expressing *D. virilis* Oskar exhibiting representative *D. virilis* Oskar localization patterns seen with anti-HA antibody stains. Micrographs are maximum intensity projections of 30 optical sections (step size = 2 $\mu$m), scale bars represent 50 $\mu$m and posterior is to the right. **B)** Frequency of ectopic patches of HA appearing away from the posterior pole in embryos from mothers expressing either *D. melanogaster* (left) or *D. virilis* (right) *oskar.* **C)** Schematic demonstrating how the plot in Fig. 6C was made. The yz projections from image stacks of anti-HA-stained embryos were eroded and fluorescent signal intensity measured 20 times. We drew a line of best fit for the curve of fluorescent signal intensity and plotted the slope of this line. **D)** Stage 10 oocytes from mothers expressing *D. melanogaster oskar* and *D. virilis oskar* stained with anti-HA antibody. Orthogonal views in the yz and xz orientations flank each image.

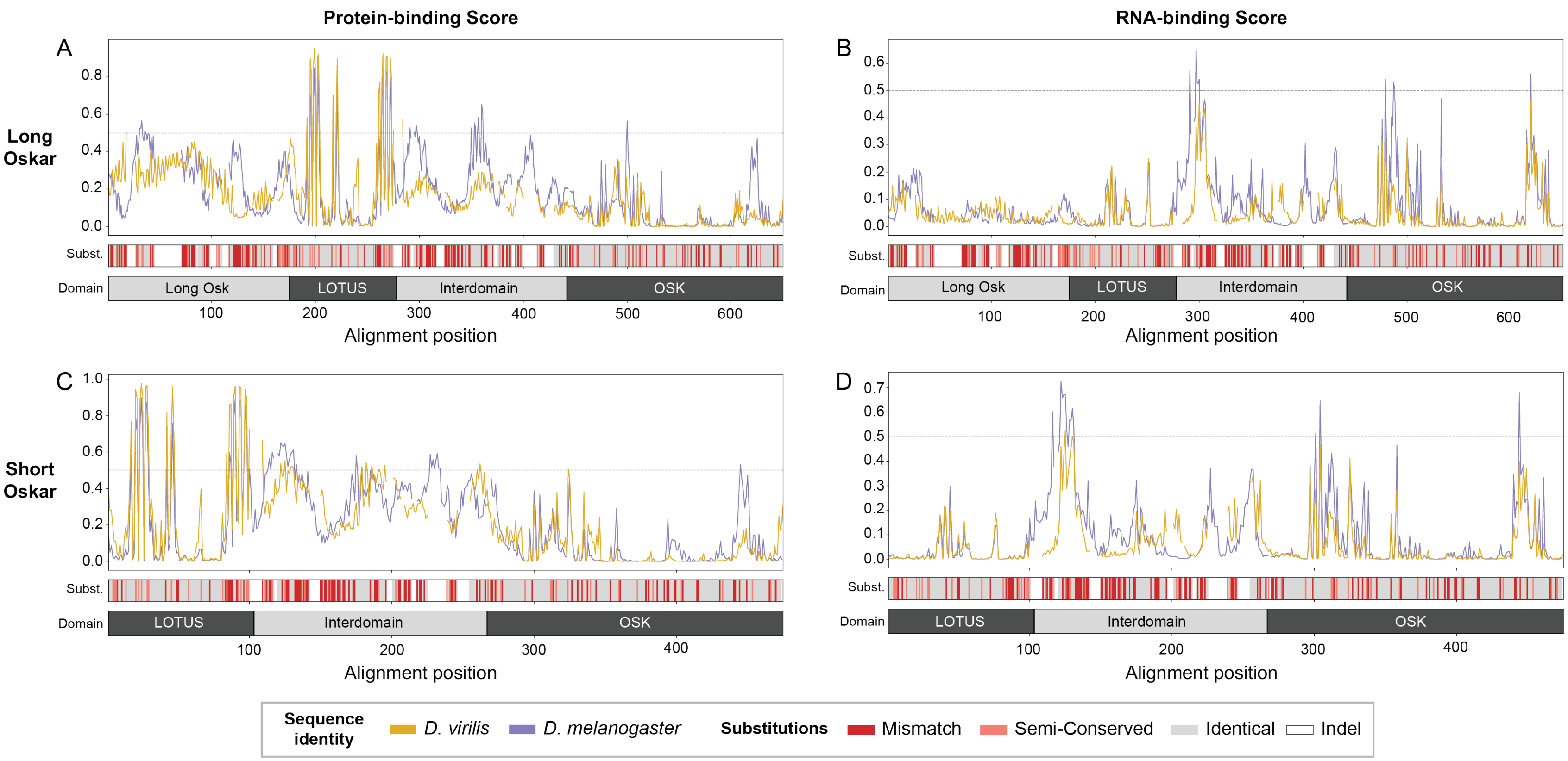


**Figure S14**. **A)** Line scans exhibiting the GPSite protein-binding score by residue for *D. melanogaster* (purple) and *D. virilis* (gold) Long Oskar sequences. **B)** Line scans exhibiting the GPSite RNA-binding score by residue for *D. melanogaster* (purple) and *D. virilis* (gold) Long Oskar sequences. **C)** Line scans exhibiting the GPSite protein-binding score by residue for *D. melanogaster* (purple) and *D. virilis* (gold) Short Oskar sequences. **D)** Line scans exhibiting the GPSite RNA-binding score by residue for *D. melanogaster* (purple) and *D. virilis* (gold) Short Oskar sequences. For all analyses, sequences were aligned using EMBOSS Needle [2]. A dotted line is drawn at 0.5. The bar below the line scans (Subst.) indicates where residues are identical between the orthologs in the alignment (gray), semi-conservative substitutions with shared biochemical properties (pink), and not conserved (red), while white indicates insertions/deletions. The four domains of Long Oskar (A, B), or the three domains of Short Oskar (C, D) are demarcated in the gray bar at the bottom of the figure (Domain).

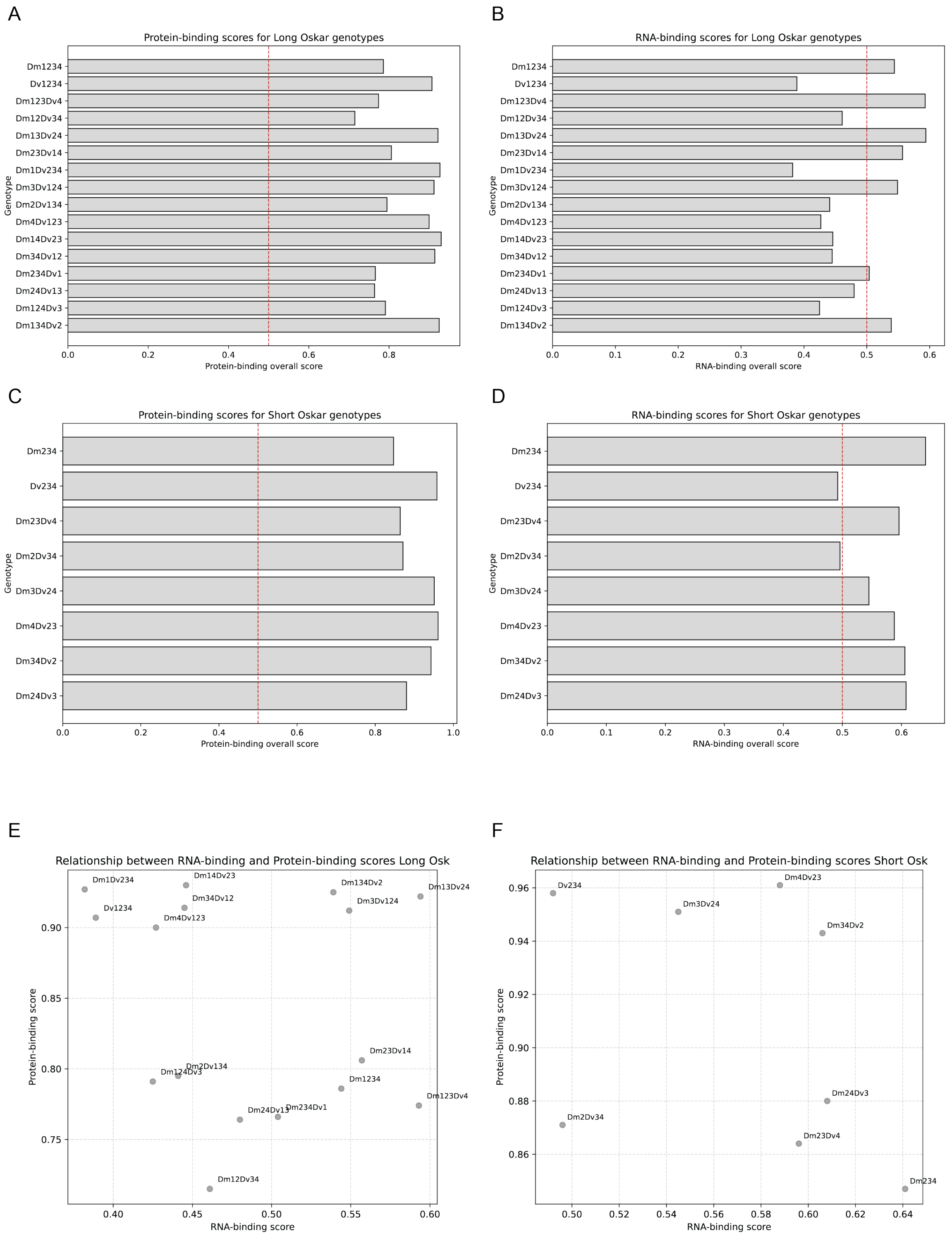


**Figure S15. A)** Overall protein-binding scores for Long Oskar chimeric proteins. **B)** Overall RNA-binding scores for Long Oskar chimeric proteins. **C)** Overall protein-binding scores for Short Oskar chimeric proteins. **D)** Overall RNA-binding scores for Short Oskar chimeric proteins. For A) through D), scores were generated using GPsite, and a red dotted line is drawn at 0.5. **E)** Scatter plot showing the correlation between predicted RNA-binding and protein-binding scores for Long Oskar chimeric proteins. **F)** Scatter plot showing the correlation between predicted RNA-binding and protein-binding scores for Short Oskar chimeric proteins. For E) and F), RNA-binding score is on the x-axis and protein-binding score is on the y-axis.

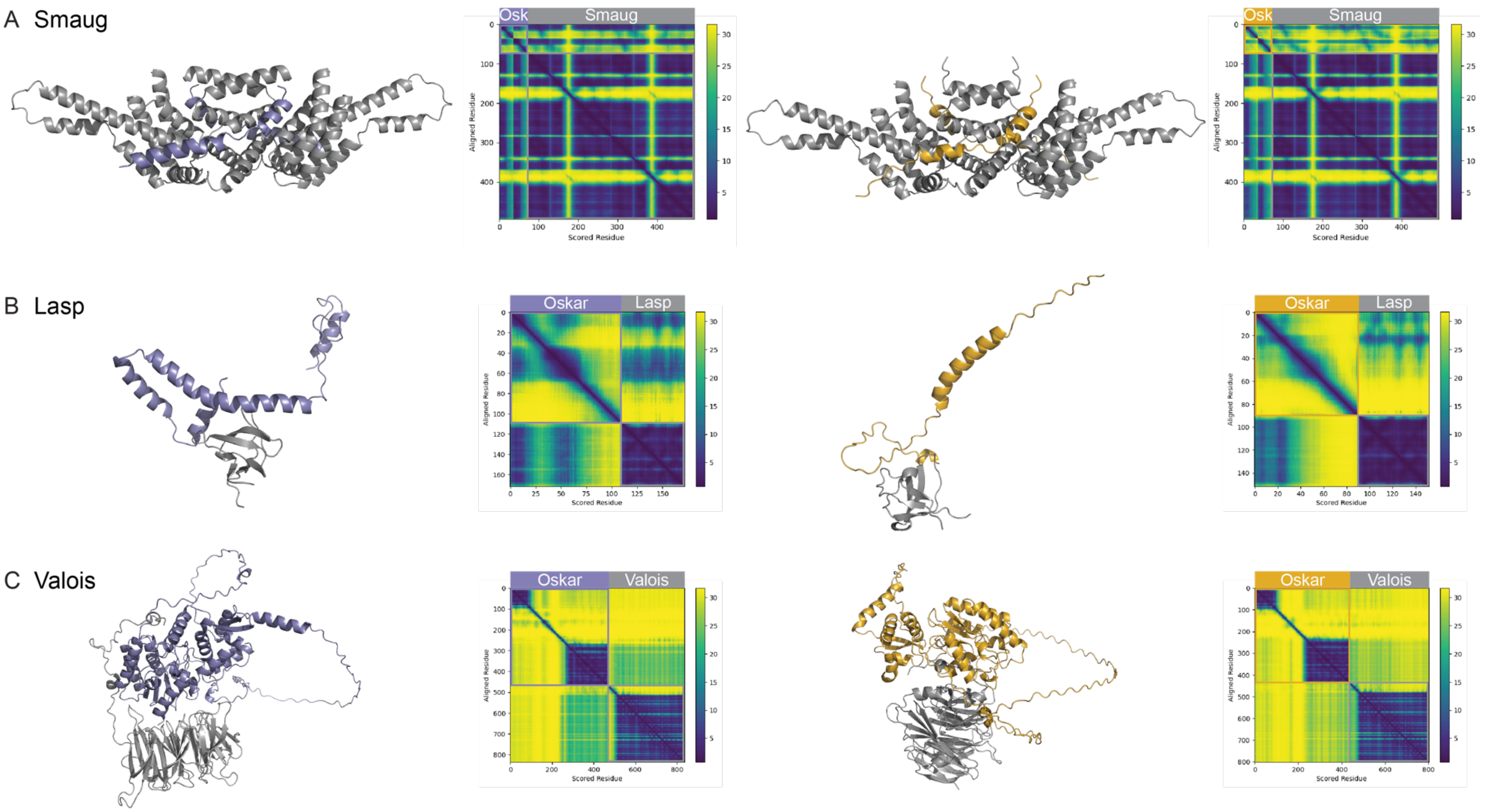


**Figure S16.** AlphaFold3 structural predictions and accompanying predicted aligned error plots for *D. melanogaster* Short Oskar (left) and *D. virilis* Short Oskar (right) with *D. melanogaster* Smaug (**A**), Lasp (**B**), and Valois (**C**). Only portions of Smaug [3], Lasp [4], and Valois [5] suggested by previous structural or *in vitro* assays to interact with Oskar, along with the corresponding portion of Oskar in this interaction, were used in the structural prediction.

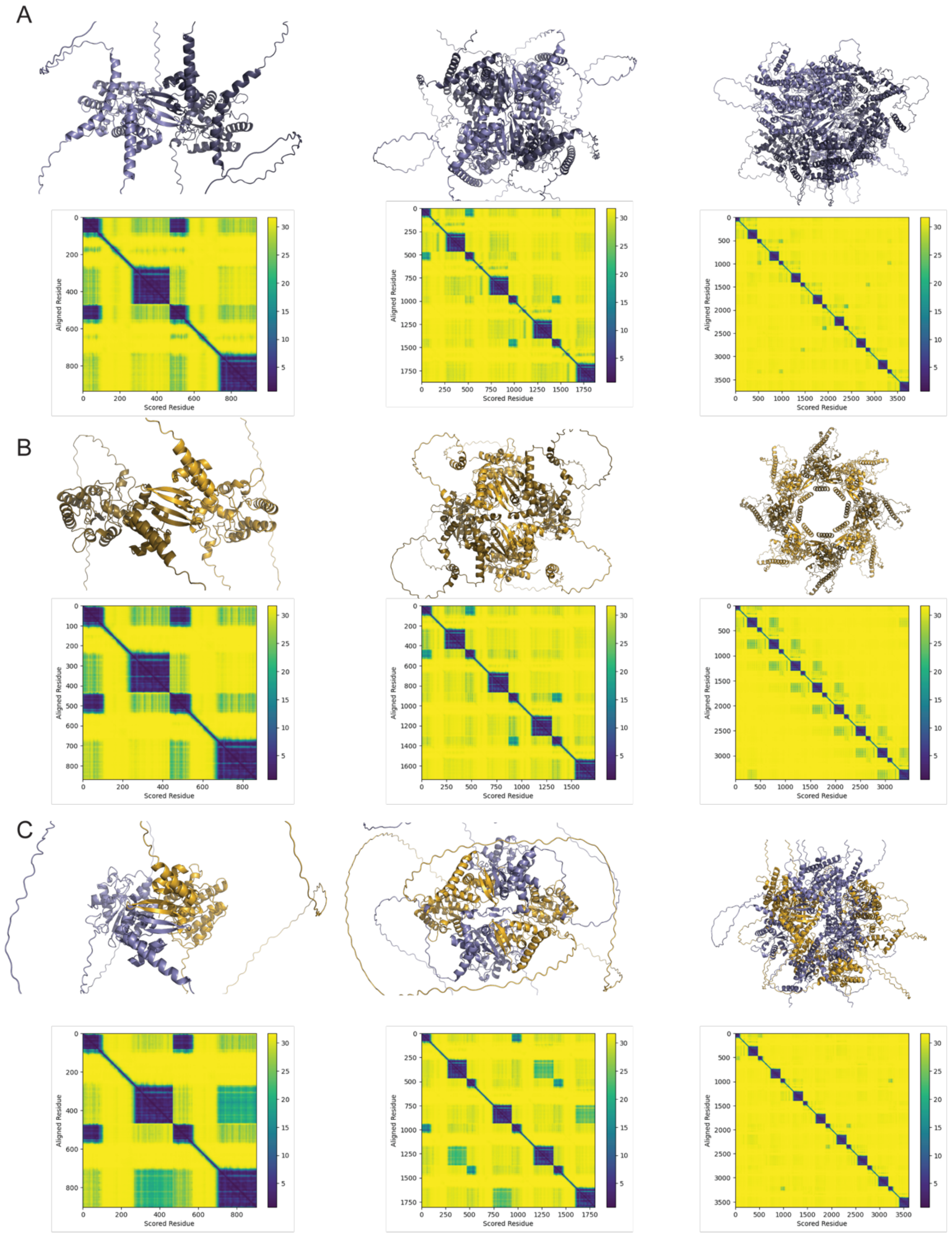
A
B
C
Aligned Residue
Scored Residue

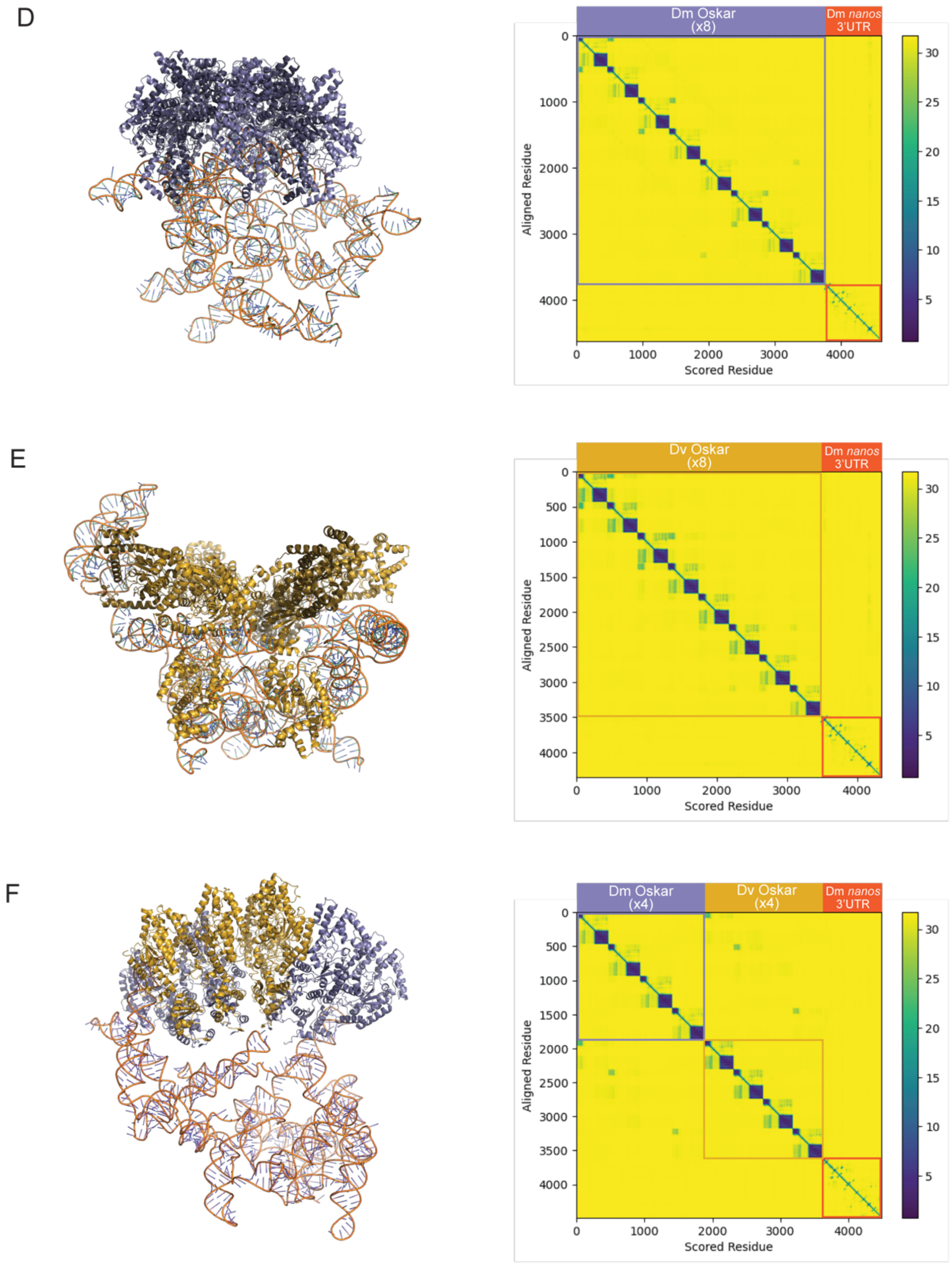


**Figure S17. A–C)** AlphaFold3 structural predictions and accompanying predicted aligned error plots for one (left column), two (middle column) or four (right column) monomers of each of *D. melanogaster* Short Oskar (**A**), *D. virilis* Short Oskar (**B**), both *D. melanogaster* and *D. virilis* Short Oskar (**C**). **D–F)** AlphaFold3 structural predictions and predicted aligned error plots for eight copies of *D. melanogaster* (**D**) *D. virilis* (**E**), and *D. melanogaster* and *D. virilis* Short Oskar (**F**) with *D. melanogaster nanos* 3'UTR.

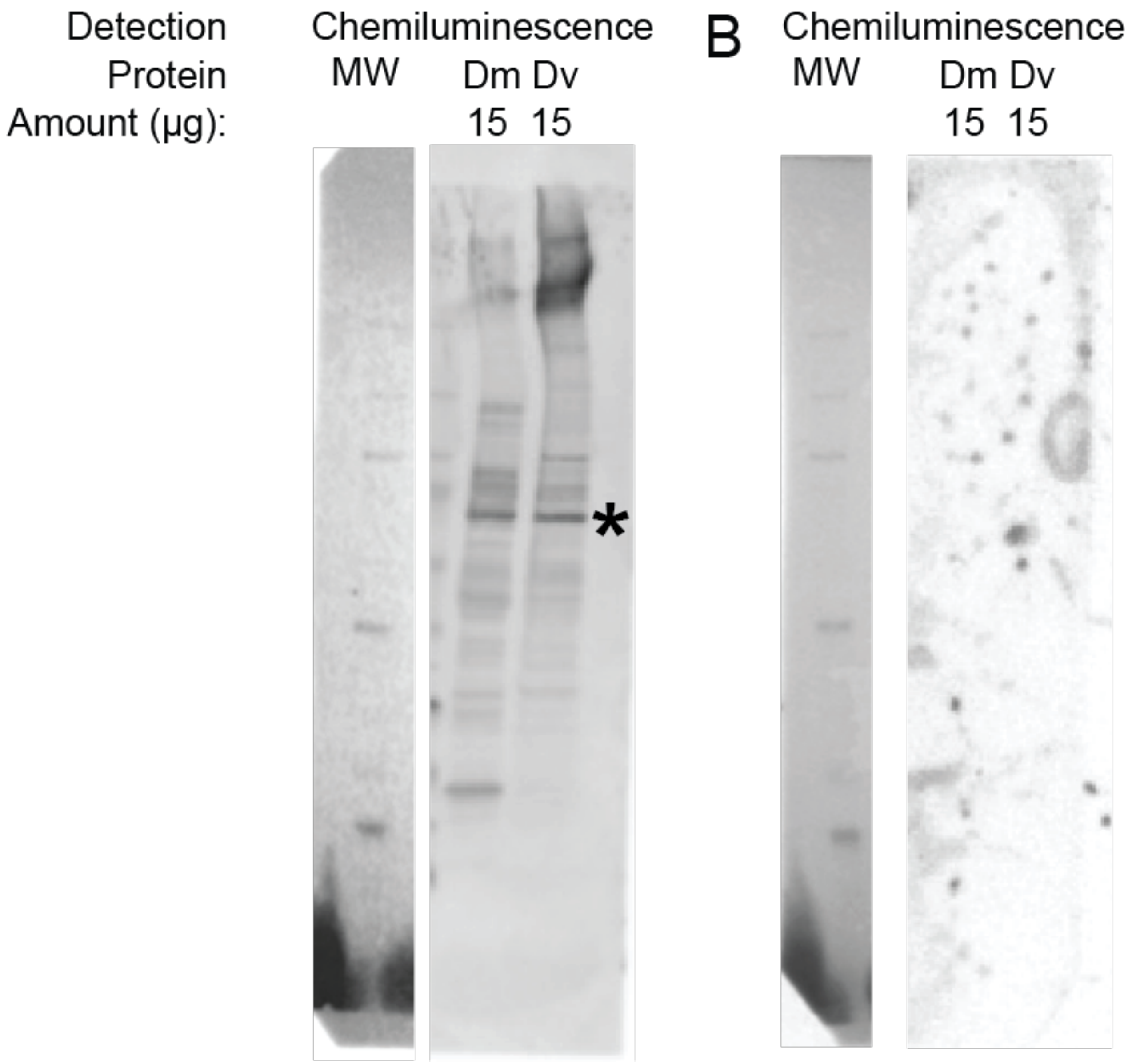


**Figure S18.** *D. virilis* anti-Oskar antibody validation. **A)** Chemiluminescent Western Blot with *D. melanogaster* (Dm) and *D. virilis* (Dv) ovary lysates incubated with anti-*Dvir* Oskar primary antibody. The amount of protein (µg) that was loaded for each lane is listed under the species identifier for each lane. Lane with molecular weight markers are marked MW. Bands at expected product sizes are marked with an asterisk (*Dmel* Oskar ~69 kDa, *Dvir* Oskar ~69 kDa). **B)** Control blot with no primary antibody. Note that this control experiment is the same one used to validate a second antibody that was generated at the same time for a different project[6] , and is reported as such in Figure S6 therein.

| **Chimeric *oskar* mRNA enrichment (*osk* null background)** | | | |
|---|---|---|---|
| Genotype | Sample size | Integrated posterior enrichment p-value | Pearson correlation p-value |
| Dm1234 | 19 | n/a | n/a |
| Dm123Dv4 | 24 | 0.00107 | 0.08475 |
| Dm124Dv3 | 8 | 0.12169 | 0.46363 |
| Dm134Dv2 | 24 | 0.14356 | 0.00017 |
| Dm234Dv1 | 24 | 0.00104 | 0.24710 |

**Table S1.** Statistical analysis of *oskar* mRNA posterior enrichment and Pearson correlation with HA in stage 10 oocytes expressing each Oskar chimera in the *oskar* RNA null background *w[1118]; Kr[If-1]/CyO; TI{w[+mC]=TI}osk[0.EGFP]*. P-values are estimated from the distribution of difference of means between sample and control from 100,000 simulated samples.

| Pole cell count (*osk* null background) | | | |
|---|---|---|---|
| **Genotype** | **Sample size** | **μ** | **p-value** |
| Dm1234 | 37 | 22.22 | n/a |
| Dv1234 | 21 | 0.00 | 0.00 |
| Dm123Dv4 | 16 | 0.00 | 0.00 |
| Dm12Dv34 | 15 | 1.13 | 0.00 |
| Dm13Dv24 | 18 | 0.00 | 0.00 |
| Dm23Dv14 | 20 | 1.55 | 0.00 |
| Dm1Dv234 | 15 | 0.00 | 0.00 |
| Dm3Dv124 | 21 | 0.00 | 0.00 |
| Dm2Dv134 | 18 | 0.00 | 0.00 |
| Dm4Dv123 | 21 | 25.19 | 0.04438 |
| Dm14Dv23 | 23 | 30.61 | 6.49E-7 |
| Dm34Dv12 | 32 | 9.88 | 4.14E-13 |
| Dm234Dv1 | 21 | 9.33 | 0.00 |
| Dm24Dv13 | 21 | 18.57 | 0.01105 |
| Dm124Dv3 | 22 | 22.95 | 0.31977 |
| Dm134Dv2 | 20 | 8.25 | 1.55E-15 |

**Table S2.** Statistical analysis of pole cell counts in stage 5 embryos expressing each Oskar chimera in the *oskar* RNA null background (*y[1] sc[*] v[1]; Kr[If-1]/CyO; TI{w[+mC]=TI}osk[0.EGFP]*). The p-values are estimated from the distribution of difference of means between sample and control from 100,000 simulated samples.

| Germ plasm enrichment (*osk* null background) | | | | |
|---|---|---|---|---|
| | **Genotype** | **Sample size** | **Integrated posterior enrichment p-value** | **Pearson correlation p-value** |
| Stage 10 oocyte: Vasa | Dm1234 | 49 | n/a | |
| | Dv1234 | 43 | 0.20058 | 0.42069 |
| | Dm123Dv4 | 63 | 6.38E-07 | 6.54E-12 |
| | Dm12Dv34 | 26 | 6.33E-12 | 7.20e-13 |
| | Dm13Dv24 | 116 | 0.00011 | 1.27e-08 |
| | Dm23Dv14 | 75 | 1.55E-08 | 7.03e-09 |
| | Dm1Dv234 | 77 | 8.99E-08 | 1.68e-05 |
| | Dm3Dv124 | 101 | 0.0 | 8.17e-07 |
| | Dm2Dv134 | 30 | 0.00121 | 0.00300 |
| | Dm4Dv123 | 70 | 0.00141 | 6.92E-06 |
| | Dm14Dv23 | 71 | 5.04E-11 | 0.08168 |
| | Dm34Dv12 | 32 | 0.04972 | 0.00017 |
| | Dm234Dv1 | 28 | 0.25847 | 0.00201 |
| | Dm24Dv13 | 81 | 0.29487 | 0.00678 |
| | Dm124Dv3 | 46 | 1.49E-06 | 0.23057 |
| | Dm134Dv2 | 54 | 3.53E-10 | 0.39111 |
| Stage 1–2 embryo: Vasa | Dm1234 | 54 | n/a | |
| | Dv1234 | 26 | 1.07E-5 | 3.93E-13 |
| | Dm123Dv4 | 32 | 0.03668 | 0.00014 |
| | Dm12Dv34 | 11 | 0.12219 | 0.01178 |
| | Dm13Dv24 | 35 | 0.00141 | 0.00 |
| | Dm23Dv14 | 28 | 5.69E-5 | 7.67E-08 |
| | Dm1Dv234 | See Table Legend & Methods | | |
| | Dm3Dv124 | 33 | 0.25939 | 0.00023 |
| | Dm2Dv134 | 30 | 0.03191 | 0.00245 |
| | Dm4Dv123 | 38 | 0.25513 | 8.31E-06 |
| | Dm14Dv23 | 39 | 1.50E-8 | 0.00 |
| | Dm34Dv12 | 56 | 0.12130 | 0.10232 |
| | Dm234Dv1 | 24 | 0.49020 | 8.26E-7 |
| | Dm24Dv13 | 32 | 0.10103 | 0.03315 |
| | Dm124Dv3 | 35 | 0.07229 | 0.01482 |
| | Dm134Dv2 | 35 | 0.41964 | 0.33832 |
| Stage 1–2 embryo: *nanos* | Dm1234 | 60 | n/a | |
| | Dv1234 | 30 | 0.17960 | 7.08E-14 |

| | | | | |
|---|---|---|---|---|
| | Dm123Dv4 | 20 | 0.00476 | 1.50E-09 |
| | Dm12Dv34 | 9 | 2.01E-06 | 0.04266 |
| | Dm13Dv24 | 25 | 0.00028 | 7.80E-25 |
| | Dm23Dv14 | 23 | 0.14142 | 1.86E-15 |
| | Dm1Dv234 | See Table Legend & Methods | | |
| | Dm3Dv124 | 39 | 3.40E-15 | 4.97E-34 |
| | Dm2Dv134 | 30 | 0.24130 | 3.31E-19 |
| | Dm4Dv123 | 37 | 0.15120 | 1.73E-06 |
| | Dm14Dv23 | 32 | 0.00104 | 0.02030 |
| | Dm34Dv12 | 32 | 0.00113 | 0.00112 |
| | Dm234Dv1 | 42 | 8.97e-05 | 0.19292 |
| | Dm24Dv13 | 22 | 0.39319 | 0.00110 |
| | Dm124Dv3 | 47 | 7.80E-08 | 0.25014 |
| | Dm134Dv2 | 33 | 1.73E-11 | 1.41E-07 |
| Stage 1–2 embryo: *gcl* | Dm1234 | 56 | n/a | |
| | Dv1234 | 24 | 0.03136 | 0.49283 |
| | Dm123Dv4 | 29 | 0.07679 | 1.33E-11 |
| | Dm12Dv34 | 9 | 0.00527 | 0.01495 |
| | Dm13Dv24 | 33 | 0.44230 | 1.345E-05 |
| | Dm23Dv14 | 21 | 0.03223 | 2.67E-14 |
| | Dm1Dv234 | See Table Legend & Methods | | |
| | Dm3Dv124 | 46 | 0.06752 | 5.54E-21 |
| | Dm2Dv134 | 21 | 0.00394 | 0.00545 |
| | Dm4Dv123 | 37 | 0.00205 | 0.02295 |
| | Dm14Dv23 | 36 | 0.00023 | 0.09993 |
| | Dm34Dv12 | 23 | 0.08356 | 0.02337 |
| | Dm234Dv1 | 59 | 0.00136 | 7.82E-11 |
| | Dm24Dv13 | 25 | 0.14451 | 0.14037 |
| | Dm124Dv3 | 36 | 0.06186 | 0.47409 |
| | Dm134Dv2 | 74 | 0.19011 | 0.01637 |

**Table S3.** Statistical analysis of Vasa, *nanos*, and *gcl* posterior enrichment and Pearson correlation with HA in stage 10 oocytes and stage 1–2 embryos expressing each Oskar chimera in the *oskar* RNA null background *w[1118]; Kr[If-1]/CyO; TI{w[+mC]=TI}osk[0.E+]*. For the chimeric genotype with the Long Osk domain from *D. melanogaster* and the other three domains from *D. virilis* (Dm1Dv234), sufficient posterior HA enrichment was found in oocytes to pass our threshold for analysis of germ plasm enrichment, but no embryos had sufficient enrichment to be included in the analysis. P-values are estimated from the distribution of difference of means between sample and control from 100,000 simulated samples.

| Pole cell count and axial patterning (endog. *osk* background) | | | | | | | |
|---|---|---|---|---|---|---|---|
| Genotype | Stage 5 embryo: Pole cell counts | | | | 1st instar larva: Cuticle patterning | | |
| | Sample size | $\mu$ | With negative control p-value | With positive control p-value | Sample size | JSD negative | JSD positive |
| Dm1234 | 21 | 62.05 | 5.99E-18 | n/a | 46 | 0 | n/a |
| Dv1234 | 21 | 27.86 | 1.95E-7 | 0.00 | 31 | 1 | 1 |
| Dm123Dv4 | 20 | 37.15 | 0.04990 | 2.10E-15 | 60 | 0.04297 | 0.04297 |
| Dm12Dv34 | 26 | 35.23 | 0.01277 | 0.00 | 53 | 0.00950 | 0.00950 |
| Dm13Dv24 | 20 | 3.80 | 0.00 | 0.00 | 51 | 1 | 1 |
| Dm23Dv14 | 20 | 49.80 | 0.00330 | 1.72E-5 | 47 | 0.09083 | 0.09083 |
| Dm1Dv234 | 20 | 40.70 | 0.28557 | 2.77E-15 | 51 | 0.04037 | 0.04037 |
| Dm3Dv124 | 20 | 37.80 | 0.06714 | 7.77E-16 | 45 | 0 | 0 |
| Dm2Dv134 | 20 | 45.35 | 0.17532 | 1.25E-6 | 54 | 0.55964 | 0.55964 |
| Dm4Dv123 | 20 | 51.55 | 0.00020 | 0.00010 | 53 | 0.00950 | 0.00950 |
| Dm14Dv23 | 20 | 59.30 | 1.41E-9 | 0.18236 | 57 | 0.01777 | 0.01777 |
| Dm34Dv12 | 30 | 57.03 | 3.47E-13 | 0.01352 | 50 | 0 | 0 |
| Dm234Dv1 | 22 | 46.68 | 0.03862 | 8.50E-9 | 52 | 0.04983 | 0.04983 |
| Dm24Dv13 | 20 | 64.30 | 1.39E-20 | 0.19133 | 51 | 0 | 0 |
| Dm124Dv3 | 33 | 44.42 | 0.15442 | 1.80E-13 | 60 | 0.06094 | 0.06094 |
| Dm134Dv2 | 20 | 38.30 | 0.07193 | 0.00 | 57 | 0.02683 | 0.02683 |
| endog. *osk* only | 34 | 42.15 | n/a | 0.00 | 55 | n/a | 0 |

**Table S4.** Left: Statistical analysis of pole cell counts in stage 5 embryos expressing one copy of each Oskar chimera in a background with endogenous *oskar* present. Right: Jensen–Shannon Divergence (JSD) calculated for the distribution of "WT," "Strong AP Defects," and "Partial AP Defects" phenotypes in each chimeric background compared to the distribution of these phenotypes in the negative (endogenous *oskar* only) and positive control (Dm1234).

| Germ plasm enrichment (endog. *osk* background) | | | | |
|---|---|---|---|---|
| | Genotype | Sample size | With negative control p-value | With positive control p-value |
| Stage 10 oocyte: Vasa | Dm1234 | 52 | 1.48E-10 | n/a |
| | Dv1234 | 53 | 4.10E-21 | 0.11051 |
| | Dm123Dv4 | 64 | 6.46E-15 | 0.37868 |
| | Dm12Dv34 | 39 | 6.55E-8 | 0.20238 |
| | Dm13Dv24 | 67 | 0.03216 | 3.17E-5 |
| | Dm23Dv14 | 42 | 4.17E-7 | 0.07693 |
| | Dm1Dv234 | 41 | 2.58E-18 | 0.08042 |
| | Dm3Dv124 | 65 | 4.96E-7 | 0.14429 |
| | Dm2Dv134 | 37 | 1.86E-31 | 0.00423 |
| | Dm4Dv123 | 74 | 1.36E-5 | 0.00587 |
| | Dm14Dv23 | 66 | 0.00073 | 0.00093 |
| | Dm34Dv12 | 37 | 4.15E-5 | 0.10557 |
| | Dm234Dv1 | 32 | 4.68E-6 | 0.01924 |
| | Dm24Dv13 | 51 | 2.25E-8 | 0.11377 |
| | Dm124Dv3 | 61 | 1.78E-9 | 0.10925 |
| | Dm134Dv2 | 40 | 1.34E-22 | 0.03775 |
| | endog. *osk* only | 179 | n/a | 1.36E-10 |
| Stage 1–2 embryo: Vasa | Dm1234 | 125 | 0.00056 | n/a |
| | Dv1234 | 16 | 1.47E-6 | 0.00238 |
| | Dm123Dv4 | 73 | 8.02e-9 | 0.00131 |
| | Dm12Dv34 | 37 | 0.31201 | 0.01605 |
| | Dm13Dv24 | 58 | 0.00016 | 0.20395 |
| | Dm23Dv14 | 56 | 0.02356 | 0.21459 |
| | Dm1Dv234 | 31 | 0.29723 | 0.03034 |
| | Dm3Dv124 | 82 | 0.00159 | 0.44346 |
| | Dm2Dv134 | 23 | 0.00578 | 0.24946 |
| | Dm4Dv123 | 134 | 1.36e-5 | 0.15072 |
| | Dm14Dv23 | 54 | 0.24605 | 0.00250 |
| | Dm34Dv12 | 60 | 0.03906 | 0.15357 |
| | Dm234Dv1 | 37 | 0.21932 | 0.02844 |
| | Dm24Dv13 | 38 | 0.36906 | 0.00322 |
| | Dm124Dv3 | 88 | 0.30718 | 9.40E-6 |
| | Dm134Dv2 | 132 | 0.43041 | 1.13e-5 |
| | endog. *osk* only | 56 | n/a | 0.00052 |
| Stage 1–2 embryo: *nanos* | Dm1234 | 49 | 4.03e-7 | n/a |
| | Dv1234 | 31 | 4.44E-16 | 0.00 |
| | Dm123Dv4 | 91 | 0.49110 | 2.66E-6 |
| | Dm12Dv34 | 36 | 0.43363 | 0.00062 |

| | | | | |
|---|---|---|---|---|
| | Dm13Dv24 | 48 | 0.00 | 0.00 |
| | Dm23Dv14 | 71 | 0.0927 | 0.00041 |
| | Dm1Dv234 | 27 | 0.26759 | 0.00028 |
| | Dm3Dv124 | 30 | 0.00459 | 2.84E-8 |
| | Dm2Dv134 | 28 | 7.89E-7 | 1.28E-14 |
| | Dm4Dv123 | 30 | 0.00275 | 0.18410 |
| | Dm14Dv23 | 104 | 5.80E-7 | 0.29581 |
| | Dm34Dv12 | 30 | 0.00017 | 0.39860 |
| | Dm234Dv1 | 23 | 0.00167 | 0.23828 |
| | Dm24Dv13 | 58 | 2.34E-5 | 0.13648 |
| | Dm124Dv3 | 50 | 0.17035 | 0.00110 |
| | Dm134Dv2 | 49 | 0.21680 | 6.42E-6 |
| | endog. *osk* only | 133 | n/a | 3.78E-7 |
| Stage 1–2 embryo: *pgc* | Dm1234 | 49 | 1.38E-18 | n/a |
| | Dv1234 | 31 | 4.24E-8 | 0.00 |
| | Dm123Dv4 | 91 | 0.01503 | 1.04E-12 |
| | Dm12Dv34 | 36 | 0.01208 | 1.62e-8 |
| | Dm13Dv24 | 48 | 3.42E-6 | 0.00 |
| | Dm23Dv14 | 71 | 1.13E-13 | 0.00 |
| | Dm1Dv234 | 27 | 0.09671 | 6.85E-10 |
| | Dm3Dv124 | 30 | 0.11079 | 5.83E-14 |
| | Dm2Dv134 | 28 | 1.97E-5 | 0.00 |
| | Dm4Dv123 | 30 | 1.09E-7 | 2.35E-5 |
| | Dm14Dv23 | 104 | 6.03E-9 | 1.12E-8 |
| | Dm34Dv12 | 30 | 5.70E-13 | 0.06107 |
| | Dm234Dv1 | 23 | 0.00412 | 1.27E-5 |
| | Dm24Dv13 | 58 | 4.75E-8 | 1.54E-5 |
| | Dm124Dv3 | 50 | 2.55E-11 | 0.00144 |
| | Dm134Dv2 | 49 | 2.76E-6 | 4.26E-6 |
| | endog. *osk* only | 133 | n/a | 0.00 |

**Table S5.** Statistical analysis of Vasa, *nanos,* and *pgc* enrichment in stage 10 *D. melanogaster* oocytes and stage 1–2 embryos expressing each Oskar chimera in a background with endogenous *oskar* present.

| | Oskar-GFP enrichment (endog. *osk* background) | | | |
|---|---|---|---|---|
| | Genotype | Sample size | Integrated posterior enrichment p-value | Pearson correlation GFP-HA p-value |
| Stage 10 oocyte: GFP | Dm1234 | 25 | n/a | n/a |
| | Dv1234 | 31 | 0.20564 | 0.28918 |
| | Dm13Dv24 | 33 | 0.24297 | 0.02389 |
| | Dm2Dv134 | 24 | 0.01444 | 0.28044 |
| Stage 10 oocyte: HA | Dm1234 | | n/a | |
| | Dv1234 | | 0.48413 | |
| | Dm13Dv24 | | 0.49845 | |
| | Dm2Dv134 | | 0.02964 | |
| Stage 1–2 embryo: GFP | Dm1234 | 12 | n/a | n/a |
| | Dv1234 | 12 | 0.21432 | 5.92E-5 |
| | Dm13Dv24 | 32 | 0.19108 | 0.29510 |
| | Dm2Dv134 | 21 | 0.03168 | 0.00045 |
| Stage 1–2 embryo: HA | Dm1234 | | n/a | |
| | Dv1234 | | 0.28852 | |
| | Dm13Dv24 | | 0.06833 | |
| | Dm2Dv134 | | 0.00187 | |

**Table S6.** Statistical analysis of Oskar-GFP posterior enrichment and Pearson correlation with HA in stage 10 oocytes and stage 1–2 embryos expressing each Oskar chimera in a background with endogenous *oskar* present and tagged with Oskar-GFP. P-values are estimated from the distribution of difference of means between sample and control from 100,000 simulated samples.

| | Nanos enrichment (endog. *osk* bacgkround) | | | |
|---|---|---|---|---|
| | Genotype | Sample size | Integrated posterior enrichment p-value | Pearson correlation p-value |
| *nanos* mRNA | Dm1234 | 21 | n/a | n/a |
| | Dv1234 | 24 | 2.53E-14 | 0.23452 |
| | Dm13Dv24 | 27 | 8.23E-5 | 0.44211 |
| | Dm2Dv134 | 17 | 0.25754 | 0.00032 |
| Nanos protein | Dm1234 | 21 | n/a | n/a |
| | Dv1234 | 24 | 0.19438 | 4.75E-27 |
| | Dm13Dv24 | 27 | 0.33927 | 3.73E-38 |
| | Dm2Dv134 | 17 | 0.00469 | 4.08E-7 |

**Table S7.** Statistical analysis of Nanos protein and *nanos* mRNA posterior enrichment and Pearson correlation with HA (proxy for chimeric Oskar protein) in stage 1–2 embryos laid by females expressing chimeric *oskar* in a background with endogenous *oskar* present. P-values are estimated from the distribution of difference of means between sample and control from 100,000 simulated samples.

| **Smaug enrichment (endog. *osk* background)** | | | |
|---|---|---|---|
| Genotype | Sample size | Integrated posterior enrichment p-value | Pearson correlation p-value |
| Dm1234 | 12 | n/a | n/a |
| Dv1234 | 12 | 0.15866 | 0.26754 |
| Dm13Dv24 | 32 | 0.44401 | 0.35755 |
| Dm2Dv134 | 21 | 0.28508 | 0.33941 |

**Table S8.** Statistical analysis of Smaug posterior enrichment and Pearson correlation with HA in stage 1–2 embryos expressing each Oskar chimera in a background with endogenous *oskar* present. P-values are estimated from the distribution of difference of means between sample and control from 100,000 simulated samples.

| Par-1 enrichment (endog. *osk* background) | | | |
|---|---|---|---|
| Genotype | Sample size | Integrated posterior enrichment p-value | Pearson correlation p-value |
| Dm1234 | 31 | n/a | n/a |
| Dv1234 | 23 | 0.43724 | 0.31283 |

**Table S9.** Statistical analysis of Par-1 posterior enrichment and Pearson correlation with HA in stage 10 oocytes expressing each Oskar chimera in a background with endogenous *oskar* present. P-values are estimated from the distribution of difference of means between sample and control from 100,000 simulated samples.

| Par-1 enrichment (*osk* null background) | | | |
|---|---|---|---|
| Genotype | Sample size | Integrated posterior enrichment p-value | Pearson correlation p-value |
| Dm1234 | 19 | n/a | n/a |
| Dm123Dv4 | 24 | 0.00517 | 3.45E-5 |
| Dm124Dv3 | 8 | 0.30397 | 0.11840 |
| Dm134Dv2 | 24 | 0.39665 | 0.49251 |
| Dm234Dv1 | 24 | 0.00297 | 0.04158 |

**Table S10.** Statistical analysis of Par-1 posterior enrichment and Pearson correlation with HA in stage 10 oocytes expressing each Oskar chimera in the *oskar* RNA null background *w[1118]; Kr[If-1]/CyO; TI{w[+mC]=TI}osk[0.EGFP]*. P-values are estimated from the distribution of difference of means between sample and control from 100,000 simulated samples.

| Radial Oskar localization | |
|---|---|
| **Genotype** | **Sample size** |
| *D. mel osk* null + *D. mel osk* transgene | 25 |
| *D. mel osk* null + *D. vir osk* transgene | 24 |
| *D. melanogaster* (Oregon R) | 25 |
| *D. virilis* | 56 |
| *D. mel osk* null + *Dm123Dv4* transgene | 34 |
| | **p-value** |
| *D. mel osk* null + *D. mel osk* transgene vs. *D. mel osk* null + *D. vir osk* transgene | 1.21E-15 |
| *D. mel osk* null + *D. mel osk* transgene vs. *D. melanogaster* (Oregon R) | 0.05499 |
| *D. mel osk* null + *D. mel osk* transgene vs. *D. virilis* | 0.00103 |
| *D. mel osk* null + *D. mel osk* transgene vs. *D. mel osk* null + *Dm123Dv4 osk* transgene | 0.30309 |
| *D. mel osk* null + *D. vir osk* transgene vs. *D. melanogaster* (Oregon R) | 0.00 |
| *D. mel osk* null + *D. vir osk* transgene vs. *D. virilis* | 4.71E-8 |
| *D. mel osk* null + *D. vir osk* transgene vs. *D. mel osk* null + *Dm123Dv4 osk* transgene | 0.00 |
| *D. melanogaster* (Oregon R) vs. *D. virilis* | 2.40E-09 |
| *D. melanogaster* (Oregon R) vs. *D. mel osk* null + *Dm123Dv4 osk* transgene | 0.00202 |
| *D. virilis* vs.*D. mel osk* null + *Dm123Dv4 osk* transgene | 0.00127 |

**Table S11.** Statistical analysis of radial HA localization in cleavage stage embryos. The p-values are estimated from the distribution of difference of means between sample and control from 100,000 simulated samples.

| **Genetic background** | **Experiment** | **Primary antibody** | **Conc.** | **Secondary antibody/ stain** | **Conc.** |
|---|---|---|---|---|---|
| Chimeras with endogenous *oskar* present | Vasa and HA in stage 10 oocytes | Rabbit anti-Vasa (gift from Akira Nakamura, Kumamoto University, Japan) | 1:3000 | Goat anti-rabbit Alexa 488 (Jackson ImmunoResearch) | 1:500 |
| | | Mouse anti-HA (AbClonal) | 1:50 | Goat anti-mouse Alexa 647 (ThermoFisher) | 1:500 |
| | | — | | Hoechst 33342 (Sigma Aldrich) | 1:2500 of 10 mg/mL stock |
| | Vasa and HA in stage 1–2 embryos | Chicken anti-Vasa (this paper) | 1:1000 | Goat anti-chicken Alexa 568 (ThermoFisher) | 1:500 |
| | | Rat anti-HA (Millipore Sigma) | 1:50 | Goat anti-rat Alexa 647 (ThermoFisher) | 1:500 |
| | | | | Hoechst 33342 (Sigma Aldrich) | 1:2500 |
| | *nanos, pgc,* and HA in stage 1–2 embryos | Mouse anti-HA (AbClonal) | 1:50 | Goat anti-mouse Alexa 488 (ThermoFisher) | 1:500 |
| | | | | Hoechst 33342 (Sigma Aldrich) | 1:2500 |
| | Vasa, phalloidin, and HA in stage 5 embryos | Chicken anti-Vasa (this paper) | 1:1000 | Goat anti-chicken Alexa 568 (ThermoFisher) | 1:500 |
| | | Mouse anti-HA (AbClonal) | 1:50 | Goat anti-mouse Alexa 647 (ThermoFisher) | 1:500 |
| | | — | | Alexa 488 phalloidin (ThermoFisher) | 1:500 |
| | | | | Hoechst 33342 (Sigma Aldrich) | 1:2500 |
| | Osk-GFP, HA in stage 10 oocytes | Rabbit Anti-GFP conjugated to Alexa Fluor 488 | 1:250 | — | — |
| | | Rat anti-HA (Millipore Sigma) | 1:50 | Goat anti-rat Alexa 568 (ThermoFisher) | 1:500 |
| | Osk-GFP, HA, and Smaug in stage 1–2 embryos | Rabbit Anti-GFP conjugated to Alexa Fluor 488 | 1:250 | — | — |
| | | Rat anti-HA (Millipore Sigma) | 1:50 | Goat anti-rat Alexa 568 (ThermoFisher) | 1:500 |
| | | Guinea pig anti-Smaug (gift from Howard Lipshitz, University of Toronto, Canada) | 1:500 | Donkey anti-guinea pig Cy5 (ThermoFisher) | 1:500 |
| | HA, *nanos,* and Nanos in stage 1–2 embryos | Rabbit anti-Nanos (gift from Akira Nakamura, Kumamoto University, Japan) | 1:1000 | Donkey anti-rabbit Alexa 647 (ThermoFisher) | 1:500 |
| | | Rat anti-HA (Millipore Sigma) | 1:50 | Goat anti-rat Alexa 488 (ThermoFisher) | 1:500 |
| Chimeras in *oskar* RNA null background | Vasa and HA in stage 10 oocytes | Rabbit anti-Vasa (gift from Akira Nakamura, Kumamoto University, Japan) | 1:3000 | Donkey anti-rabbit Alexa 647 (ThermoFisher) | 1:500 |
| | | Mouse anti-HA (AbClonal) | 1:50 | Goat anti-mouse Alexa 568 (ThermoFisher) | 1:500 |
| | | — | | Hoechst 33342 (Sigma Aldrich) | 1:2500 |
| | Vasa and HA in stage 1–2 embryos | Rabbit anti-Vasa (gift from Akira Nakamura, Kumamoto University, Japan) | 1:3000 | Goat anti-rabbit Alexa 568 (ThermoFisher) | 1:500 |
| | | Rat anti-HA (Millipore Sigma) | 1:50 | Goat anti-rat Alexa 647 (ThermoFisher) | 1:500 |

| | | | | | |
|---|---|---|---|---|---|
| | | — | | Hoechst 33342 (Sigma Aldrich) | 1:2500 |
| | *nanos* and HA in stage 1–2 embryos | Rat anti-HA (Millipore Sigma) | 1:50 | Goat anti-rat Alexa 647 (ThermoFisher) | 1:500 |
| | | — | | Hoechst 33342 (Sigma Aldrich) | 1:2500 |
| | *gcl* and HA in stage 1–2 embryos | Rat anti-HA (Millipore Sigma) | 1:50 | Goat anti-rat Alexa 647 (ThermoFisher) | 1:500 |
| | Vasa, phalloidin, and HA in stage 5 embryos | Chicken anti-Vasa (this paper) | 1:1000 | Goat anti-chicken Alexa 568 (ThermoFisher) | 1:500 |
| | | Rat anti-HA (Millipore Sigma) | 1:50 | Goat anti-rat Alexa 647 (ThermoFisher) | 1:500 |
| | | — | | Alexa 488 phalloidin (ThermoFisher) | 1:500 |
| | | — | | Hoechst 33342 (Sigma Aldrich) | 1:2500 |
| | Vasa and HA in stage 14 embryos | Rabbit anti-Vasa (gift from Akira Nakamura, Kumamoto University, Japan) | 1:3000 | Goat anti-rabbit Alexa 568 (ThermoFisher) | 1:500 |
| | | Rat anti-HA (Millipore Sigma) | 1:50 | Goat anti-rat Alexa 647 (ThermoFisher) | 1:500 |
| | | — | | Hoechst 33342 (Sigma Aldrich) | 1:2500 |
| | Par-1, *osk,* and HA in stage 10 oocytes | Rabbit anti-Par-1 (gift from Daniel St. Johnston, Gurdon Institute, UK) | 1:2500 | Donkey anti-rabbit Alexa 647 (ThermoFisher) | 1:500 |
| | | Mouse anti-HA (AbClonal) | 1:50 | Goat anti-mouse Alexa 488 (ThermoFisher) | 1:500 |
| Wild-type *D. melanogaster* | Vasa, Oskar, and actin in stage 10 oocytes | Rat anti-Oskar (Extavour Lab) | 1:250 | Goat anti-rat Alexa 647 (ThermoFisher) | 1:500 |
| | | Rabbit anti-Vasa (gift from Akira Nakamura, Kumamoto University, Japan) | 1:3000 | Goat anti-rabbit Alexa 488 (Jackson ImmunoResearch) | 1:500 |
| | | — | | Rhodamine phalloidin (ThermoFisher) | 1:250 |
| | | | | Hoechst 33342 (Sigma Aldrich) | 1:2500 |
| | Vasa and Oskar in stage 1–2 embryos | Rabbit anti-Oskar (gift from Anne Ephrussi, EMBL, Germany) | 1:3000 | Donkey anti-rabbit Alexa 647 (ThermoFisher) | 1:500 |
| | | Chicken anti-Vasa (this paper) | 1:1000 | Goat anti-chicken Alexa 568 (ThermoFisher) | 1:500 |
| | | — | | Hoechst 33342 (Sigma Aldrich) | 1:2500 |
| | Vasa in stage 5 embryos | Chicken anti-Vasa (this paper) | 1:1000 | Goat anti-chicken Alexa 568 (ThermoFisher) | 1:500 |
| | | — | | Alexa 488 phalloidin (ThermoFisher) | 1:500 |
| | | | | Hoechst 33342 (Sigma Aldrich) | 1:2500 |
| Wild-type *D. virilis* | Vasa, Oskar, and actin in stage 10 oocytes | Chicken anti-Oskar (this paper) | 1:1000 | Goat anti-chicken Alexa 647 (ThermoFisher) | 1:500 |
| | | Rabbit anti-Vasa (gift from Akira Nakamura, Kumamoto University, Japan) | 1:3000 | Goat anti-rabbit Alexa 488 (Jackson ImmunoResearch) | 1:500 |
| | | — | | Rhodamine phalloidin (ThermoFisher) | 1:250 |
| | | | | Hoechst 33342 (Sigma Aldrich) | 1:2500 |
| | Vasa and Oskar in stage 1–2 embryos | Chicken anti-Oskar (this paper) | 1:1000 | Goat anti-chicken Alexa 568 (ThermoFisher) | 1:500 |
| | | Rabbit anti-Vasa (gift from Akira Nakamura, Kumamoto University, Japan) | 1:3000 | Goat anti-rabbit Alexa 488 (Jackson ImmunoResearch) | 1:500 |
| | | — | | Hoechst 33342 (Sigma Aldrich) | 1:2500 |
| | Vasa in stage 5 embryos | Rabbit anti-Vasa (gift from Akira Nakamura, Kumamoto University, Japan) | 1:3000 | Goat anti-rabbit Alexa 488 (ThermoFisher) | 1:500 |

| | | | | |
|---|---|---|---|---|
| | | — | Rhodamine phalloidin (ThermoFisher) | 1:500 |
| | | | Hoechst 33342 (Sigma Aldrich) | 1:2500 |

**Table S12.** Antibodies and stains used in immunofluorescence experiments. “Conc.” = antibody or stain concentration.

| Hybridization chain reaction probes | | | |
|---|---|---|---|
| **Species** | **Probe** | **Amplifier** | **Fluorophore** |
| *D. melanogaster* | *oskar* | B1 | Alexa 488 |
| | *Nanos* | B5 | Alexa 555 |
| | *germ cell-less* | B3 | Alexa 594 |
| | *polar granule component* | B4 | Alexa 647 |
| | *oskar 3'UTR* | B5 | Alexa 555 |
| *D. virilis* | *oskar* | B1 | Alexa 488 |
| | *nanos* | B5 | Alexa 555 |
| | *germ cell-less* | B3 | Alexa 594 |
| | *polar granule component* | B4 | Alexa 647 |

**Table S13.** List of hybridization chain reaction (HCR) probes used in fluorescence *in situ* hybridization experiments.

| Primer sequences | |
|---|---|
| **Name** | **Sequence** |
| pVAL22_gOsk_bridge2_R | CACCCTCAGCTCCAAAGCCTCGAGTGTTTTGCCATTTTACA |
| pVAL22_gOsk_bridge1_R | CACTAGTAGTACCAGCTTATAACTTCGTATAATGTGAATTGGGTACCGGGCC |
| pVAL22_gOsk_bridge2_F | TGTAAAATGGCAAAACACTCGAGGCTTTGGAGCTGAGGGTG |
| pVAL22_gOsk_bridge1_F | GGCCCGGTACCCAATTCACATTATACGAAGTTATAAGCTGGTACTACTAGTG |
| pVALIUM22_R6 | GCTTTGGAGCTGAGGGTG |
| pVALIUM22_F6 | ACATTATACGAAGTTATAAGCTGGTACTACTAGTG |
| Dmel_gOsk_HA_F1 | GATGTTCCGGATTACGCTTAAGTTGGGTTCTTAATCAAGATACATATATG |
| Dmel_gOsk_HA_R1 | GTATGGGTAACTACCATACTCCAGACTCGTTTCAATAACTTGC |
| HA_F1 | GGTAGTTACCCATACGATGTTCCGGATTACGCT |
| Dmel_Dvir4_R1 | TAATCCGGAACATCGTATGGGTAACTACCATAATACTCCAAGTTTTCTTCAATCATGTGC |
| Dmel_upstream_F1 | CGCTTGGAGGAAAGTGATCCAATATTCTTG |
| Dmel_Dvir1_F1 | GAATATTGGATCACTTTCCTCCAAGCGATGGCGACATTCAGAAGTGAATTC |
| Dmel_upstream_F2 | CTGTTGAATTCACTTCTGAATGTCGCCATCGCTTGGAGGAAAGTGATCCAATATTCTTG |
| Dmel_Dvir1_F2 | ATGGCGACATTCAGAAGTGAATTCAACAGCGTGCCGAACACTTACAACC |
| Dvir_Losk_F161 | ATGACCATCATGGACAACACCTATATT |
| Dvir_int_R1 | CCAGCAGTATGGAGCGTATTTCAG |
| Dvir_Losk_F261 | GACAGTGTGAGCAACCCAATC |
| Dvir_Losk_F401 | TCGGACTATGATGCATATCTATTGG |
| Dvir_Losk_F387 | TCGTGTCCATCAACGCAAC |
| Dvir_int_F1 | GTACATGGTTATGATTTCGAGGCT |
| Dvir_int_F2 | CAAATCAACCGACTCCATTCCC |
| Dmel_gOsk_insert_F1 | AACTGCTCACCGTTACTTCTGTTG |
| Dmel_Losk_R414 | GTTAATCGTCAGCAGAGAATCGTTG |
| Dmel_Losk_R400 | TAAAATCGTTGGCGTGGGC |
| Dmel_Losk_F140 | ACCATCATCGAGAGCAACTATATATCCG |

| | |
|---|---|
| Dmel_Losk_gOsk_R1 | CATCGCTTGGAGGAAAGTGATC |
| Dmel_Dvir4_R2 | CCGGAACATCGTATGGGTAACTACCATAATACTCCAAGTTTTCTTCAATCATGTGC |
| Dmel_upstream_F3 | GAATTCACTTCTGAATGTCGCCATCGCTTGGAGGAAAGTGATCCAATATTCTTG |
| Dmel_Dvir1_F2 | ATGGCGACATTCAGAAGTGAATTCAACAGCGTGCCGAACACTTACAACC |
| Dvir_Losk_R386 | CAACGTTGTTGGCGTGCGCG |
| Dm4_Dv3_F1 | CGCGCACGCCAACAACGTTGACCAGTGGAACCTACAACGATTCTC |
| Dm4_HA_R1 | CCGGAACATCGTATGGGTAACTACCATACTCCAGACTCGTTTCAATAACTTGC |
| Dvir_Losk_R260 | GCGTTGTCGCTGCTGCAG |
| Dm3_Dv2_F1 | TACTGCAGCAGCGACAACGCTACAGCAGCGGAGCTCCG |
| Dm3_Dv4_R1 | TGTTGCGTTGATGGACACGATAAAATCGTTGGCGTGGGC |
| Dm2_Dv1_F1 | GTGCAAGTATCACCAGCTCCAGCATGACCATCATCGAGAGCAACTATATATC |
| Dvir_Losk_R160 | GCTGGAGCTGGTGATACTTGCAC |
| Dm2_Dv3_R1 | TGGATTGGGTTGCTCACACTGTCGTCGCTGGTGCGCTCTTTC |
| Dm1_Dv2_R1 | CCAATATAGGTGTTGTCCATGATGGTCATGTTGTTCTCGCTGGTGTTGCTG |
| Dm1_gOsk_F1 | GAATATTGGATCACTTTCCTCCAAGCGATGGCCGCAGTCACAAGTG |

**Table S14**. Primer sequences used for cloning *oskar* chimeras.
